# VICTree - a Variational Inference method for Clonal Tree reconstruction

**DOI:** 10.1101/2024.02.14.580312

**Authors:** Harald Melin, Vittorio Zampinetti, Andrew McPherson, Jens Lagergren

## Abstract

Clonal tree inference brings crucial insights to the analysis of tumor heterogeneity and cancer evolution. Recent progress in single cell sequencing has prompted a demand for more advanced probabilistic models of copy number evolution, coupled with inference methods which can account for the noisy nature of the data along with dependencies between adjacent sites in copy number profiles. We present VICTree, a variational inference based algorithm for joint Bayesian inference of clonal trees, together with a novel Tree-structured Mixture Hidden Markov Model (TSMHMM) which combines HMMs related through a tree with a mixture model. For the tree inference, we introduce a new algorithm, LARS, for sampling directed labeled multifurcating trees. To evaluate our proposed method, we conduct experiments on simulated data and on samples of multiple myeloma and breast cancer. We demonstrate VICTree’s capacity for reliable clustering, clonal tree reconstruction, copy number evolution and the utility of the ELBO for model selection. Lastly, VICTree’s results are compared in terms of quality and speed of inference to other state-of-the-art methods. The code for VICTree is available on GitHub: github.com/Lagergren-Lab/victree.

## 1 Introduction

The well-established clonal theory of cancer [27] describes cancer development as a process of phenotypically distinct cell sub-populations, referred to as clones, undergoing Darwinian evolution driven by somatic genetic mutations. The resulting tumor heterogeneity is a substantial barrier for successful treatment [10]. To analyze the genetic content of a tumor, the recent single cell whole genome sequencing (scWGS) technology Direct Library Preparation (DLP) [35], has shown proficiency in producing data of thousands of single cells suitable for copy number (CN) analysis [30, 35]. Yet, the data exhibits imperfections and the experimental process is inherently noisy. These limitations motivates the necessity of adopting phylogenetic methods able to incorporate uncertainty in the inferences.

CN deletion and duplication events may effect multiple genomic sites. This has sparked recent surge in research of models able to accommodate for dependencies within the CN profile (CNP), e.g., in form of clustering methods [29], and phylogenetic methods which also account for site-dependent CN evolution [16, 25, 30, 32, 34]. Despite their demonstrated strengths, the phylogenetic methods either rely on heuristic approaches [16, 32, 34] which lack the rigor of probabilistic modeling, or on simplifying assumptions and pre-processing steps that break the full Bayesian aspiration [25, 30].

Markov chain Monte Carlo (MCMC) has played a pivotal role in Bayesian phylogenetics, both in classical setting [7,12] and in the CN tumor setting [25,30]. However, the introduction of dependencies within the CNPs causes the size of the hidden state-space to be subject to combinatorial explosion, which has proven difficult for MCMC in classical phylogenetics [13], where the marginalization over the hidden states to evaluate the likelihood becomes intractable. In the CN clonal tree setting, a cluster assignment and CN state must be given in order to evaluate the likelihood. Thus, MCMC algorithms dedicated to fully explore this state-space will be limited. Therefore, previous tumor phylogenetic CN based methods circumvent this problem by requiring the unobservable CNPs as input [30], these are obtained e.g. by running HMMCopy [22], and by implicitly segmenting the CNPs through a transformation [30] or user-input to select potential breakpoints [25]. Unfortunately, the decisions taken in such preprocessing cannot be reversed later and may lead to artifacts. Variational inference [2, 14] is alternative to MCMC and has delivered substantial speed-up of inference in a variety of settings [1], including in the context of classical phylogenetics [13, 18, 26, 36, 37], but is yet to enter the tumor phylogenetic arena.

We present VICTree, a method that utilizes VI for joint Bayesian inference of site-dependent CN evolution, clonal tree reconstruction, cell-to-clone clustering and cell specific read baselines, using only GC-correction and normalization for data pre-processing. VICTree draws inspiration from both the tumor and classical phylogenetic literature.

To model site-dependent CN evolution, we extend the dependency graph between hidden states of the phylo-HMM model [31], which describes the evolution of site-dependent chromatin’s in a phylogenetic tree, and infuse it with a transition model tailored to tumor phylogenetics. To this we add a mixture model and an emission model, extending the Mixture Hidden Markov Model (MHMM) [11, 28] of CopyMix [29]. The combination yields a Tree structured MHMM (TSMHMM), a novel type of HMM, suitable for tumor phylogeny from the clonal perspective.

In order to achieve efficient and accurate inference, we employ a coordinate ascent VI (CAVI) approach and derive update equations to the associated variational distributions. To estimate the non-standard expected value w.r.t tree topologies, we employ Importance sampling [17] (IS) as in [18]. Furthermore, we introduce a novel algorithm for labeled arborescences sampling, LARS, to use as proposal distribution for our IS procedure. Lastly, we identify that and describe why VI based clonal analysis is haunted by the problem of component degeneracy, i.e., high similarity between clones. We resolve the problem by extending our VI algorithm with a novel domain specific split-and-merge-operation, thereby, taking VI based Bayesian clonal analysis from the theoretically interesting to the practically applicable. As a bonus, the use of VI equips us with a lower bound on the marginal likelihood, a metric which encodes Occam’s Razor, fit for model selection [24]. By assembling these methods and the TSMHMM, we reach our aspiration of a fully automated Bayesian inference procedure of clonal phylogenies.

### Contributions

− **VICTree** - The first framework for fully automated Bayesian inference over genomic site-dependent CN evolution, clonal tree topologies, cell-to-clone assignment, clonal prevalence and cell specific emission baseline and precision.
− **TSMHMM** - A novel type of HMM, the Tree-Structured Mixture-HMM, suitable for the tumor clonal tree context.
− **LARS** - a novel algorithm for sampling labeled multifurcating rooted trees (also known as *arborescences*) induced by a weighted graph.
− **Split-and-merge algorithm** - a novel Split-and-merge algorithm tailored to the context of clonal trees, taking VI based clonal analysis from the theoretically interesting to the practically applicable.

## 2 Background

Given observations, **y**, described by a set of latent variables and parameters, **x**, of a generative model *p*(**x, y**), Bayesian inference is concerned with finding the posterior distribution:

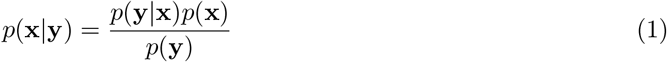

Unfortunately, the posterior is intractable due to the denominator in 1, called the marginal likelihood or evidence, and therefore an approximate Bayesian inference procedure must be applied.

Variational Inference (VI) [2, 14] addresses this by finding a surrogate distribution, called a variational distribution, *q*_*ϕ*_(**x**), parameterized by *ϕ*, which best approximates the posterior *p*(**x**|**y**), in terms of minimal KL-Divergence, *D*_*KL*_, between *q* and *p*. Furthermore, instead of minimizing *D*_*KL*_ explicitly, maximizing the Evidence Lower Bound, hereinafter referred to as *ELBO*, is preferred. The ELBO is defined as:

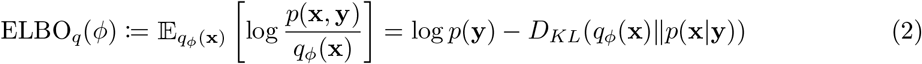

Considering that *p*(**y**) is a constant, it is straightforward to see that maximizing the ELBO w.r.t. *ϕ* is equivalent to minimizing *D*_*KL*_, which is the ultimate objective of the inference process, and, since *D*_*KL*_ *≥* 0, the ELBO lower bounds *p*(**y**). What remains is to choose a family of distributions for *q*_*ϕ*_(**x**). A popular choice is the mean-field family, which assumes that *q*_*ϕ*_(**x**) factorizes as 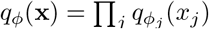 where *x*_*j*_ is a subset of **x**. This assumption leads to the coordinate ascent VI (CAVI) method [2, 14] where each 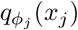is optimally updated iteratively by:

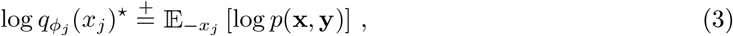

where 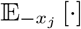 denotes the expectation over all latent variables but *x*_*j*_ and 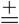 symbolizes equality up to additive constants. When simplifying 3 for a specific model, we might be able to recognize the distribution of 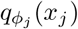 along with the optimal *ϕ*_*j*_. In the particular case where the full conditional, *p*(*x*_*j*_|*y, x*_*−j*_), is in the exponential family, for conjugate priors w.r.t. *p*(*y, x*_*−j*_|*x*_*j*_), the corresponding *q*^*\**^(*x*_*j*_) will be in the same family as the prior *p*(*x*_*j*_) [9].

Before we divulge in the derivation of *q*_*ϕ*_(**x**) of VICTree, we first describe the proposed model *p*(**x, y**) in the next section.

## 3 Proposed model

This section describes our proposed model and the related variables under analysis. The model can be split into two modules:

− *CopyTree*: a tree-structured Markov model that generates CNPs with horizontal and hierarchical dependencies, and
− the mixture emission model, an emission model which associates the observations to the hidden states of CopyTree by a mixture over the tree nodes (hereon referred to as *clones*).

In Section 4 we explain how variational inference is used for estimating the posterior distribution of the model.

### 3.1 The CopyTree model

We introduce a tree-structured Markov Model (MM) for latent CNPs linked by parent-child dependencies within a given tree topology, *T*, containing arcs *A*(*T*). Given an arc, *uv ∈ A*(*T*) the CN at site *m* for clone *v*, 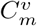, depends not only on 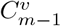, but also on the parent clone’s CNs at site *m* and *m −* 1, 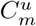 and 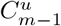. Such dependencies are depicted in Figure 2a. We can see this as a set of hierarchically interlaced Markov models where the transition probability from one CN state to the subsequent one is a function of four CN states and an arc-specific transition probability, *ε*_*uv*_ *∈* [0, 1].

**Fig. 1:**
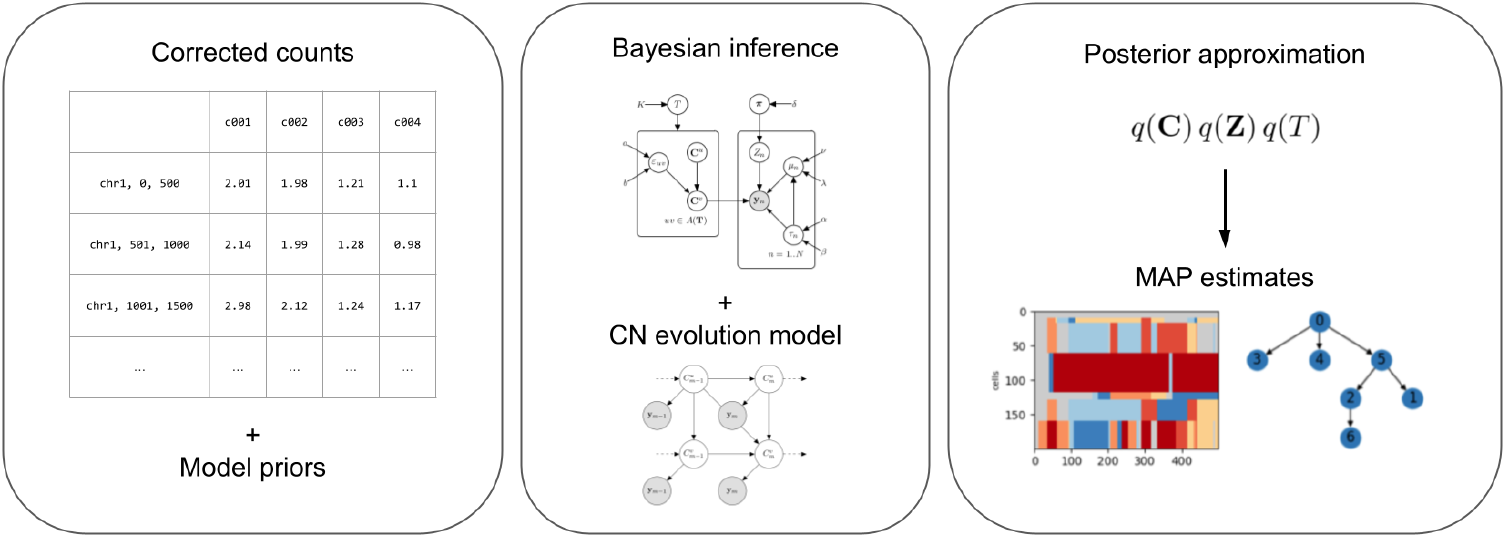
Overview of VICTree. Bin counts are the only required input data and have to be corrected for GC content and ploidy (neutral CN equals 2). The method performs Bayesian inference through CAVI updates by optimizing the parameters of the model here introduced. The output produced is the set of variational distribution, from which it is possible to retrieve MAP estimates for downstream analysis (CN calling, clustering of cells into clones and CN-aware evolution tree).

**Fig. 2:**
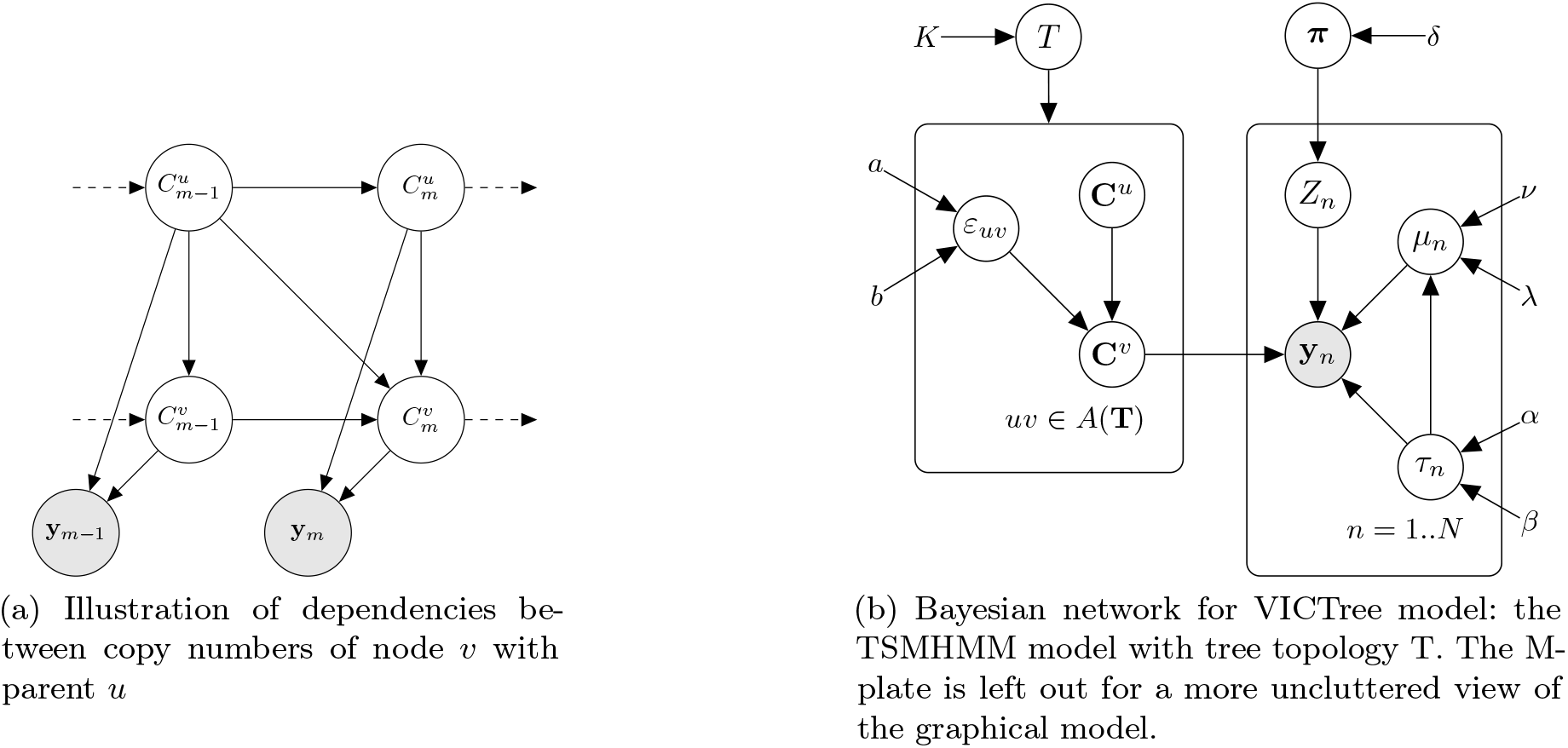
Variables dependency graphs.

We call this function the *CN coherence function* (4) and it models the CN evolution both along the clone CNP and over the ancestry according to:

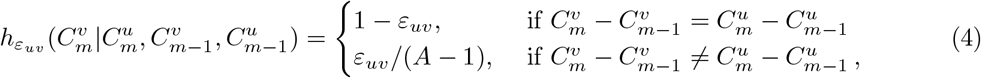

where *A* is the maximum CN. If *ε*_*uv*_ is small, the CN coherence function formalizes the intuition that if we observe a CN change in the CNP of *u*, then we are likely to observe the same CN change in *u*’s child *v*. Note, the state 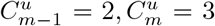 and 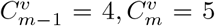 is treated as coherent despite the evident difference in CN between these two clones, e.g. between 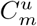 and 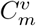. This may at first seem like a model-misspecification, however, this difference will be reflected in an incoherence in a preceding bin, i.e. a bin *m*^*′*^ *< m −* 1, of the CNPs.

We construct a first simple model by treating all CN changes that do not satisfy the CN coherence condition as equal, similar to the Jukes-Cantor model [15] in classical phylogeny. Furthermore, *ε*_*uv*_ can be interpreted as a distance between the two clones, indeed we will refer to it as *arc distance*. For what concerns the edge cases, the root node is set as a healthy clone with constant CN i.e. 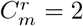 for all *m*, and the initial state probability for any internal node depends only on the initial state of the parent node.

This fully describes the generative process behind the hidden states of our TSMHMM. To associate the hidden states with observed data, we turn the emission model in the next section.

### 3.2 The mixture emission model

We relate our observations, **Y**, to the latent CopyTree state by a Mixture Hidden Markov Model (MHMM) [11, 28], inspired by the MHMM with Gaussian emissions used in [29], with cell-to-clone assignments **Z**, concentration parameter ***π***, cell baseline ***μ*** and emission precision ***τ***. More specifically, for cell *n* assigned to clone *k* through **Z**, we let the corrected reads, 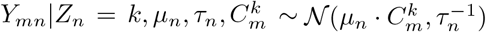, which we will refer to as *corrected reads*, since they are assumed to be corrected for GC content and normalized so that 2 is the neutral CN. GC correction is a pre-processing step that corrects for the read bias of low GC content regions and transforms the data from natural to real numbers, thus motivating a continuous distribution for the observational model.

### 3.3 Tree structured MHMM

The combined statistical model is a Tree-structured MHMM for a fixed tree *T*. Together with a uniform prior distribution over the finite set of all possible tree topologies 𝒯 _*K*_ for a fixed number of nodes *K*, this completes our generative model, *p*(**Y, C, *ε*, Z, *π***, *T*, ***μ, τ***). The Bayesian network for the full generative model can be seen in figure 2b and the corresponding distributions are:

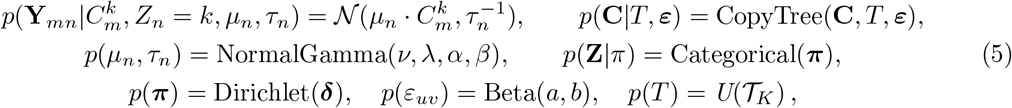

Where CopyTree(**C**, *T*, ***ε***) represents the joint distribution of the interleaved Markov model described in Section 3.1, which depends on the topology by means of its arcs, and on the arcs’ distances ***ε***. We now turn our attention to the framework for performing joint Bayesian inference on the model.

## 4 VICTree

In the Bayesian setting, the main goal is to infer the posterior distribution, that is the probability of the model parameters given the observation, as mentioned in Section 2. We assume the following factorization of the variational distribution:

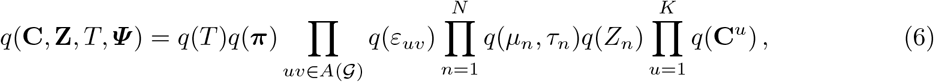

where *𝒢* is the fully connected directed graph with *K* nodes and *A*(*𝒢*) is the set of directed arcs, which does not contain self connections. With abuse of notation, each *q* over different variables has unique learnable parameters, which are left implicit for uncluttered notation.

Unsurprisingly, as our variational updates for **Z, *π, ε, μ, τ*** follow the conditional conjugacy mentioned in Section 2, our CAVI updates become:

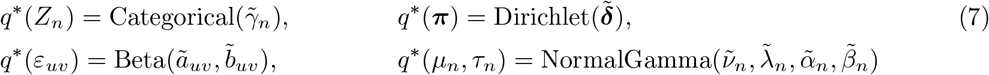

where the expressions of the variational parameters, 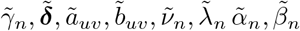 can be found in the Supplementary Material, together with their derivations.

The updates of *q*(*T*) and *q*(**C**), however, are less orthodox. For *q*(**C**), the CAVI update can be simplified to the form:

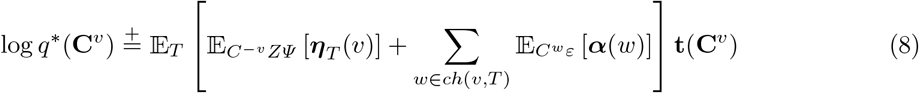

where, ***η***_*T*_ (*v*) is the vector of natural parameters associated to the sufficient statistic **t**(**C**^*v*^) of node *v*, ***α***_*T*_ (*w*) is the vector of natural parameters viewed from child *w* of *v*, which has the same form of ***η***_*T*_ (*v*) thanks to conjugacy. **t**(**C**^*v*^) is the sufficient statistic of the CNP at node *v*. These terms can be seen as variational messages coming from both parent and children of node *v* as in [33] (see Supplementary Material, Section S2). The outermost expectation value w.r.t. *T* is non-trivial and must be approximated, e.g. using a Monte Carlo sampling. Appealingly, the optimal *q*(**C**) for the TSMHMM corresponds to a distinct non-homogeneous Markov chain for each clone, where the parameter update balances two terms; one enforcing the CN evolution w.r.t. the CopyTree prior and the other forcing the distribution to adapt the observed data.

A detailed derivation and description of 8 can be found in Section S3.1.

The CAVI update of *q*(*T*) simplifies as:

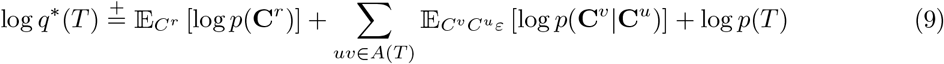

which is not on the form of some known distribution. To handle this we adopt a similar procedure as in [18]; we construct a fully connected graph, *𝒢* and let the associated weight matrix, *W* (𝒢), be determined by the second term of eq. 9 (Note: the first and third term in 9 are constant over trees), allowing us to evaluate sampled trees on *q*(*T*). Whenever an update requires an expectation w.r.t. *T*, e.g. in eq. 8, we employ IS [17], using our novel sampler LARS as proposal distribution, *g*(*T*). This concludes our VI framework, the pseudocode for the algorithm can be found in Algorithm 1.

### 4.1 LARS

Here, we describe our IS proposal distribution *g*(*T*), a distribution over labeled arborescences of *W* (*𝒢*), induced by our LARS algorithm. Ideally, we would sample a tree with respect to its normalized weight using the matrix tree theorem. However, this is costly, instead we prescribe the Diaconis heuristic for the Markov chain tree theorem [3, 6], by replacing the sum of tree weights with the maximum spanning tree (MST) weight. Consider the directed graph *G* with arcs *A*(*G*). For any subset of arcs *S ⊆ A*(*G*) and arc *e ∈ A*(*G*), let *A*(*S, e*) be the set of arborescences of *G* that contains both the arcs of *S* and *e* and, moreover, let *A*^*′*^(*S, e*) be the set of arborescences of *G* that contain the arcs of *S* but not *e*. Let *e*_1_, …, *e*_*L*_ be a random permutation of the arcs of *G*. Then the next state of the Markov chain tree is obtained by selecting the arc *e*_*i*_ with probability

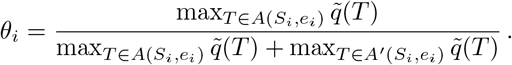

The maximization problem is solved by Edmonds’ algorithm [8]. The proposal distribution from which we sample is then constructed as a product of Bernoulli trials, i.e., 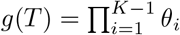, if *e*_*i*_ is accepted. Further details of the algorithm is described in Section S6 of the Supplementary Material.

### 4.2 The split-and-merge algorithm

Inference of mixture models, e.g., in the context of VI, MCMC or the EM-algorithm, is susceptible to mixture component degeneracy, resulting in convergence to poor local optima. This section examines the origin of this issue in CN based clustering and an algorithm to counteract the phenomena. The algorithm is designed to address domain specific issue of shared hidden states between the clones; segments of the CNPs of each clone may be shared between all clones, *clonal profile*, between two or more clones, *semi-clonal profile*, or fully clone specific, *subclonal profile*.

We will illustrate how this effects our VI optimization with an extreme case: one of the *K* clusters, e.g. the *k*^th^ cluster, adapts *q*(**C**^*k*^) to the signal of the clonal profile in the data before the remaining clusters. Consequently, each cell n will be assigned to the *k*^th^ cluster with a probability *q*(*Z*_*n*_) which depends on the ratio between the clonal profile and subclonal profile signal in the observations. If the ratio is high, then optimization converges to a degenerate local optima where *q*(*Z*_*n*_ = *k*) = 1 for all *n* and *q*(*C*^*k*^) learns the clonal profile and averages the subclonal part over all cells. As *q*(*Z*_*n*_ = *j*) = 0 for all *j ?*= *k* and *q*(**C**^*j*^) is updated w.r.t. *q*(*Z*_*n*_ = *j*), the result is *K −* 1 degenerate components.

To escape these local optima, we adapt a split-and-merge algorithm for VI [4] to the clonal clustering context. We merge clusters *i* and *j* based on the distance between *q*(**C**^*i*^) and *q*(**C**^*j*^). We split a cluster *i* into a degenerated cluster *j* if there exists a group of cells within *i*, that when updating *q*(**C**^*j*^) on that group, maximize the ELBO w.r.t. to all other possible splits. More details are given in section S7.

### 4.3 ELBO calculation

To monitor the status of the inference procedure and assess the fitness of the current posterior approximation we use the ELBO presented in Section 2. In the VICTree setting, the ELBO is derived as follows:

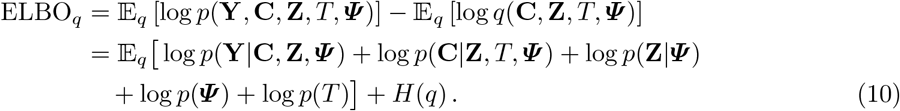

Each one of the expectation terms can be computed separately and such derivations are shown in the Supplementary Material. It should be noted that any expectation over the tree topology space is approximated as explained in Section 4.1, therefore it is useful to view each such expectation as 𝔼_*T,X*_ [*f* (*T, X*)] = 𝔼_*T*_ [𝔼_*X*_ [*f* (*T, X*)]] where *f* is some function of *T* and a generic r.v. *X*.

## 5 Experimental results

In order to evaluate our method we conduct experiments on both synthetic data, to get advantage of a ground-truth, and real data, so to assess the practical efficacy of inference. We compare our results on real data against HMMCopy [22] (in copy-number calling) and CONET [25] (in copy-number calling and clonal tree inference).

### 5.1 Simulated data

Data have been simulated from the generative model presented in Section 3. We show the results of two distinct experiments: one to assess the overall performance of VICTree and provide a benchmark on its downstream analysis capability, and one to assess robustness on the amount of copy-number mutations. In the first experiment, for three sizes of datasets, *small, medium, large* i.e. respectively (*K, M, N*) ∈ {(6, 1000, 300), (9, 3000, 600), (12, 5000, 1000)}, we compute *adjusted-rand index* (ARI) on cell assignment, copy-number calling *mean absolute deviation* (MAD), and *arc accuracy* on the reconstructed tree (i.e. the proportion of arcs in the MAP tree that also belong to the ground truth tree).

In the second experiment we simulate data of *medium* size with increasing number of mutations along the genome and compute the same measures. Mutations are generated at a higher frequency by multiplying the first parameter of the Beta distribution of *ε*_*uv*_ by a factor *≥* 1, which we denote as *mutation rate*. The results for this experiment are shown in the Supplementary Material S2.

For all these experiments, VICTree has been configured by setting the hyper parameters to fixed values, partly dependent on the size of the datasets, and mainly derived from explorative data analysis on real datasets such as the ones here under study. More specifically, the update *step-size* was set to 0.3 and LARS sample size *L* = 5. We fix the model priors (as defined in Equation 5) to ***δ*** = (3, …, 3), *ν* = 1 *λ* = 20 *× M, α* = 500, *β* = 50, *a* = 5, *b* = *M*. The value of *K* was set to the same (normally unknown) value from which the data has been generated. We motivate this choice by showing that by running VICTree with different values of *K* on synthetic data, the ELBO clearly suggests which value is best for each dataset (see Figure 4).

Our method performs well in all the above mentioned objectives, namely cell clustering, clonal CN calling, and tree reconstruction, achieving perfect score in most of the datasets as shown in Figure 3.

**Fig. 3:**
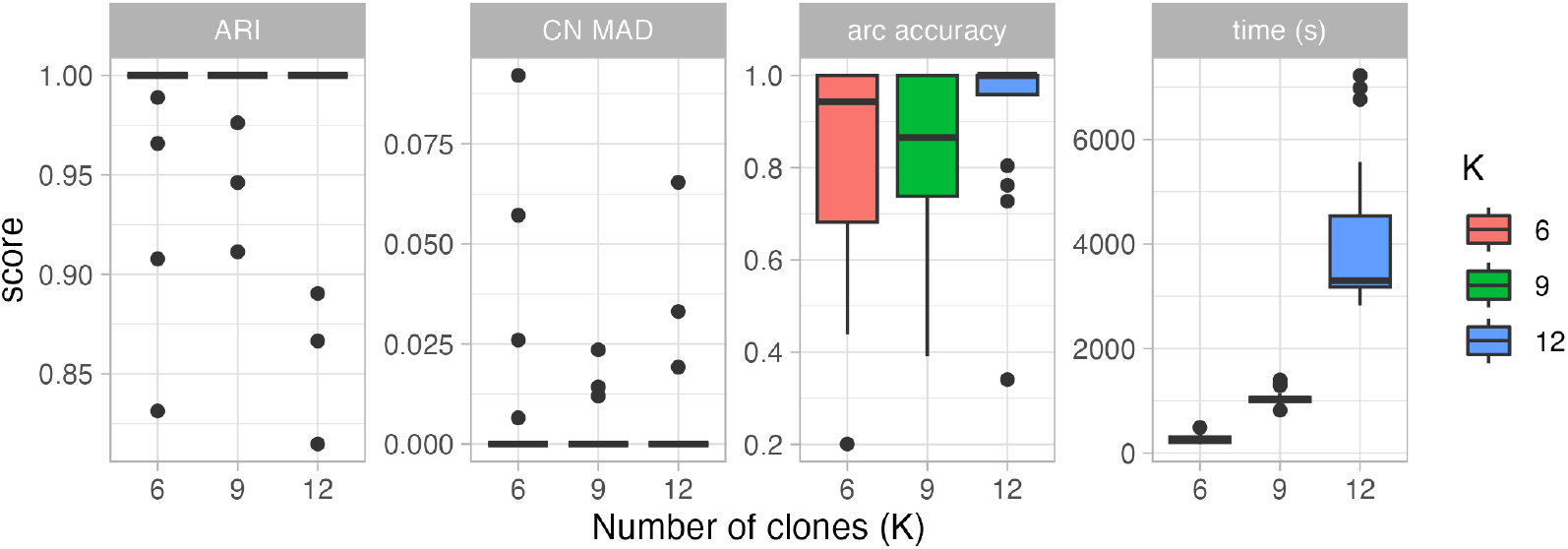
Scores against ground truth over 20 synthetic datasets for three values of *K* (number of clones). Clustering performance is quantified by the Adjusted-Rand index while CN calling performance by mean absolute deviation. Arc accuracy is the proportion of arcs in the ground truth tree that are included in the MAP estimate tree. Dots represent outliers. Time refers to the execution of VICTree on one core of a Intel Xeon Gold 6130.

**Fig. 4:**
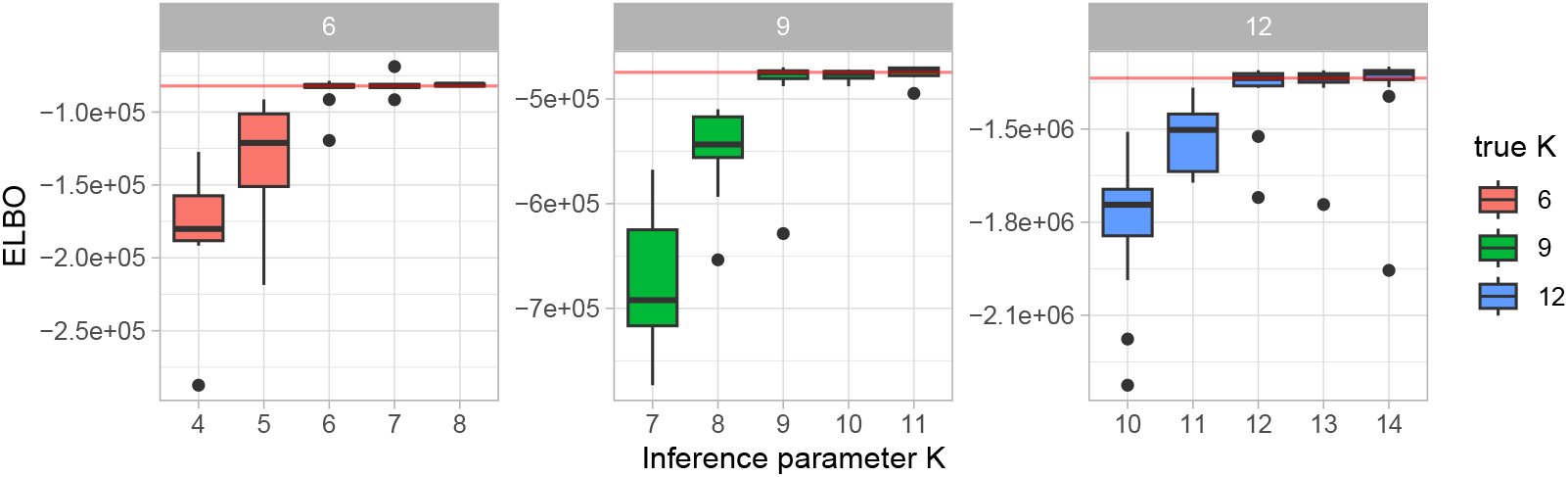
ELBO plot showing model selection on synthetic data. Each facet shows runs on equally sized datasets (i.e. small, medium, large). On the horizontal axis the number of nodes *K* chosen for inference. Boxplots show the result over 20 datasets.

### 5.2 Multiple Myeloma datasets

We apply VICTree to several datasets of multiple myeloma extracted using the 10x Genomics technology [23]. This section provides a description of the results over the sample MM29, while the results of other two samples (MM03 and MM04) are left in the Supplementary Material together with the comparison to HMMcopy CN calling (see Section S1.4).

In Figure 5 we show the clonal CNP of each cell that is the Viterbi path of *q*(**C**^*v*^), ordered by the estimated MAP cell-to-clone assignment given by the inferred *q*(***Z***), together with the inferred MAP tree. Inference has been performed over a number of different *K* values. We only present the results with *K* = 14 as it is shown to be a fair trade-off between model complexity and ELBO (see Supplementary Figure S6), although there’s evidence in support of a larger *K*.

**Fig. 5:**
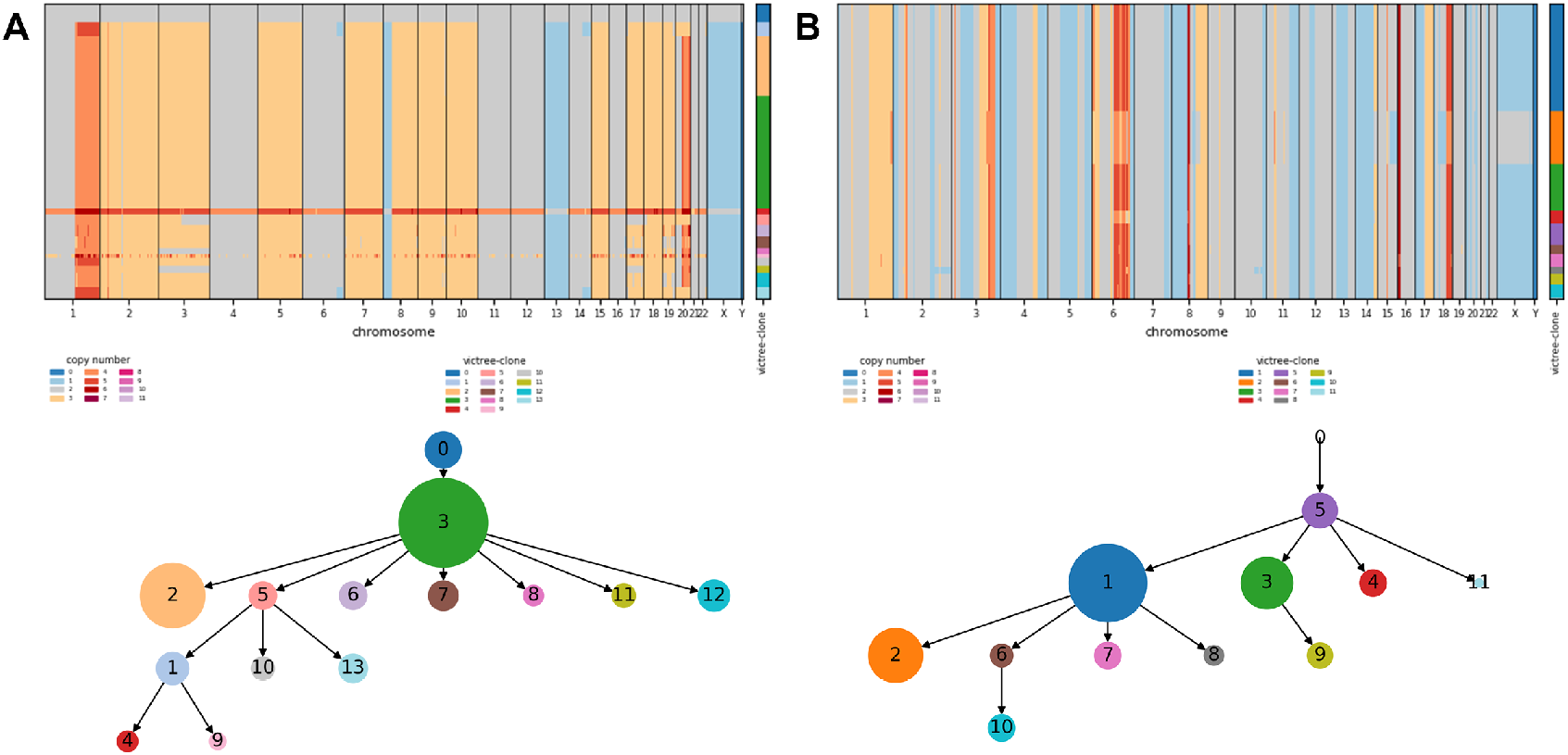
Inferred clonal copy number profile and MAP estimate of the clonal tree of two real datasets. **A** Sample MM29 (multiple myeloma) is obtained by running VICTree with *K* = 14 while **B** sample SA501X3F (breast cancer) with *K* = 12.

### 5.3 SA501X3F xenograft breast cancer dataset

We run our method on the DLP dataset SA501X3F [35] which features 260 single cells for 18175 bins of width 150kb, setting the maximum number of clones K to 12. VICTree is able to infer the clonal copy number profile, assign cells to the clones and reconstruct a clonal tree (MAP estimate) within an hour^5^, just by providing it the corrected reads matrix without any pre-processing step. Results are shown in Figure 5 and a comparison with CONET [25] CN calling and reconstructed clonal tree is left in the Supplementary Material S3.

## 6 Discussion

We have presented a framework for fully automated Bayesian inference in clonal tumor phylogenetics, requiring a minimal amount data pre-processing steps and user inputs. To achieve this, we have presented several new modular models and subroutines: CopyTree and the TSMHMM with associated CAVI updates, LARS a rooted tree sampler, and a novel domain specific split-and-merge algorithm, which may be applied independently in the field of tumor phylogenetics. The composite algorithm, VICTree, is demonstrated to infer clonal trees, CNPs and cell-to-clone assignments in high accordance with the ground truth in 5.1 and retrieves similar MAP output to that of HMMCopy in 5.2 with the addition of a distribution over trees explaining the process of CN evolution. In relation to CONET, VICTree infers trees that, while capturing the same evolution process, summarize the single-cells information in a less cluttered fashion (S3). Furthermore, VICTree’s is robust to high mutation rates in the data S2, a scenario where the user-input required by CONET is cumbersome.

The presented probabilistic model features a large number of dependencies between variables, so to deeply explain the nature of the data from its source, not relying on other intermediate steps or broad assumptions which might not hold in all cases. This substantially increases the complexity of inference (e.g. for a site-specific HMM) and our method alleviates this with the efficiency of variational inference.

Bayesian inference have been already adopted in the field of tumor phylogenetics [19, 25], and while other methods only partially engage in the inference process, often relying on additional tools and approximation, our approach offers a comprehensive Bayesian framework supported by rigorous statistical techniques.

In terms of quality of inference, VICTree demonstrates strong clustering and CN calling capabilities, while simultaneously inferring trees in the vicinity of the true underlying tree. Furthermore, the experiment in Section 5.1 demonstrates VICTree’s robustness to high rates of CN events events, where other methods currently struggle. Also, despite the large set of learned parameters, VICTree reaches convergence in short overall execution time as shown in Figure 3, making it more efficient than traditional MCMC methods.

Regarding the experimental results, we show that LARS is a robust algorithm for sampling labeled arborescences from a weighted graph. It works effectively alongside the inferred weights in the variational distribution over the tree space.As stated in the introduction, the ELBO’s lower bound on the marginal likelihood offers a unique tool for model selection in tumor phylogenetics. Indeed, the ELBO is higher for K’s in the vicinity of the true K according to 4. In real data experiments however, such as S4, the ELBO does not plateau with increasing K. In the context of clonal tree reconstruction, we can therefore choose to limit the granularity of clones based on the ELBO trend line, by identifying a trade off between a lower model complexity and a higher fitness to the data.

### 6.1 Future Work

The modular framework of VICTree facilitates a myriad of independent improvement opportunities. The proposal distribution *g*(*T*) could be swapped with a scheme that is more fit for larger K. Furthermore, ensembles or mixtures of variational distributions can be applied to better accommodate for multi-modality [20, 21]. The CN evolution model specified in Equation (4) can be improved to explicitly model specific CN events such as whole-genome duplication events. Furthermore, the observational model is unidentifiable, in that it is observationally equivalent under different combinations of *μ*_*n*_ and 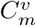. In light of this, although we still achieve accurate inference by properly setting the model priors, an observational model which includes ploidy (i.e. the average CN of a clone) could be developed so to improve the TSMHMM. Here, the unpacked update equations in Section S3 expedites such framework development.

## Supplementary Material

### S1 Additional experiments

#### S1.1 Synthetic data simulation

#### S1.2 Robustness on high mutation rate

#### S1.3 Breast cancer sample

#### S1.4 Multiple Myeloma samples

**Fig. S1:**
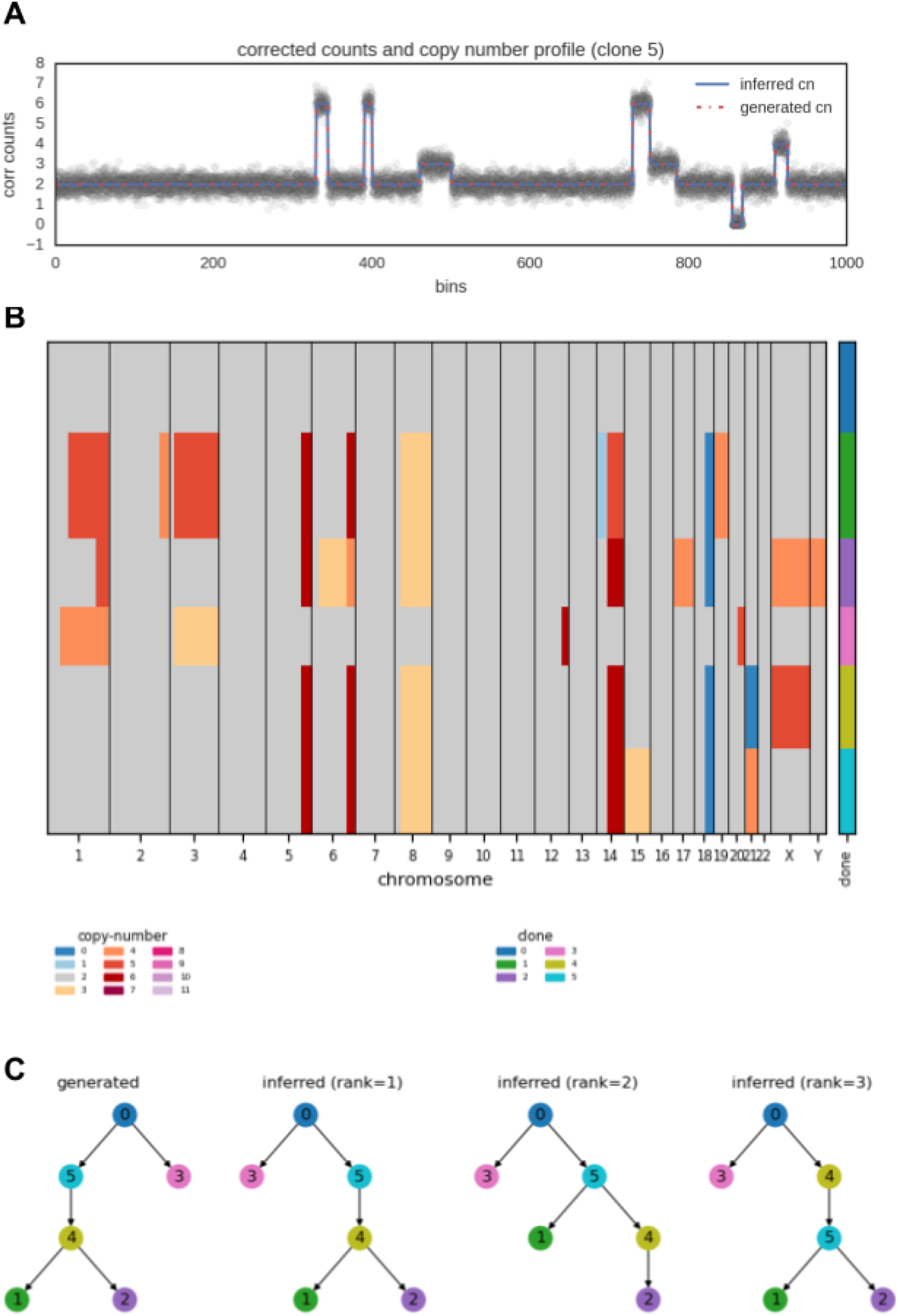
Example of synthetic dataset with *K* = 6 clones. **A** Corrected counts of cells assigned to clone 5 with ground-truth and inferred copy number profile (CN-MAD = 0.0). **B** Clonal copy number profile with cells on the vertical axis, arranged by inferred clonal assignment (ARI = 1.0). **C** Generated (ground-truth) tree and set of inferred trees, ranked by importance in the final sample from *q*(*T*) using LARS.

**Fig. S2:**
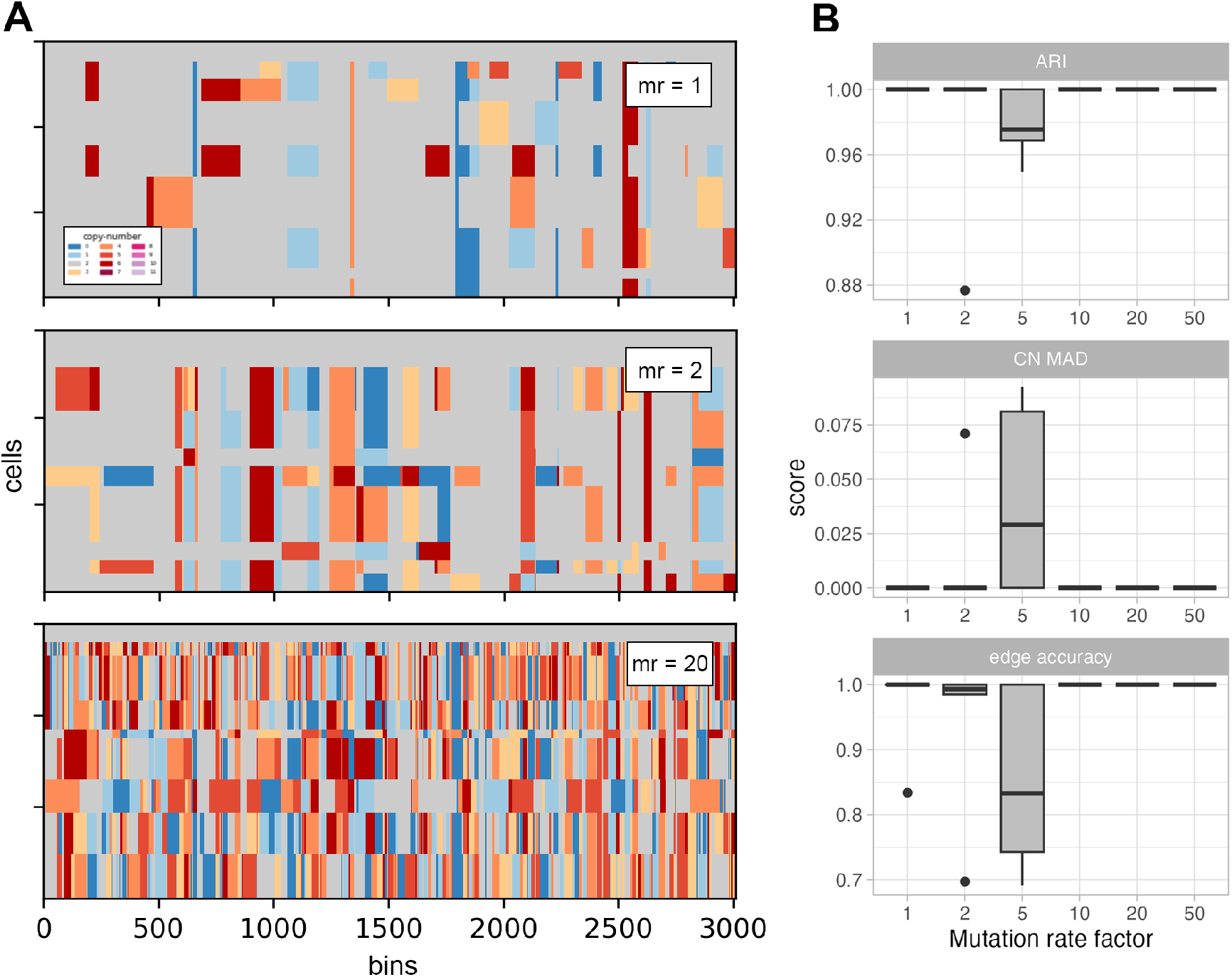
Illustration on the robustness over increasing rate of copy number mutation. **A** Simulated copy number evolution of 9 clones with three different values for the mutation rate parameter: 1 (default), 2 and 20. **B** ARI (clustering), CN MAD (copy number calling) and edge accuracy scores against ground truth on 10 synthetic datasets for 6 different values of mutation rate parameter.

**Fig. S3:**
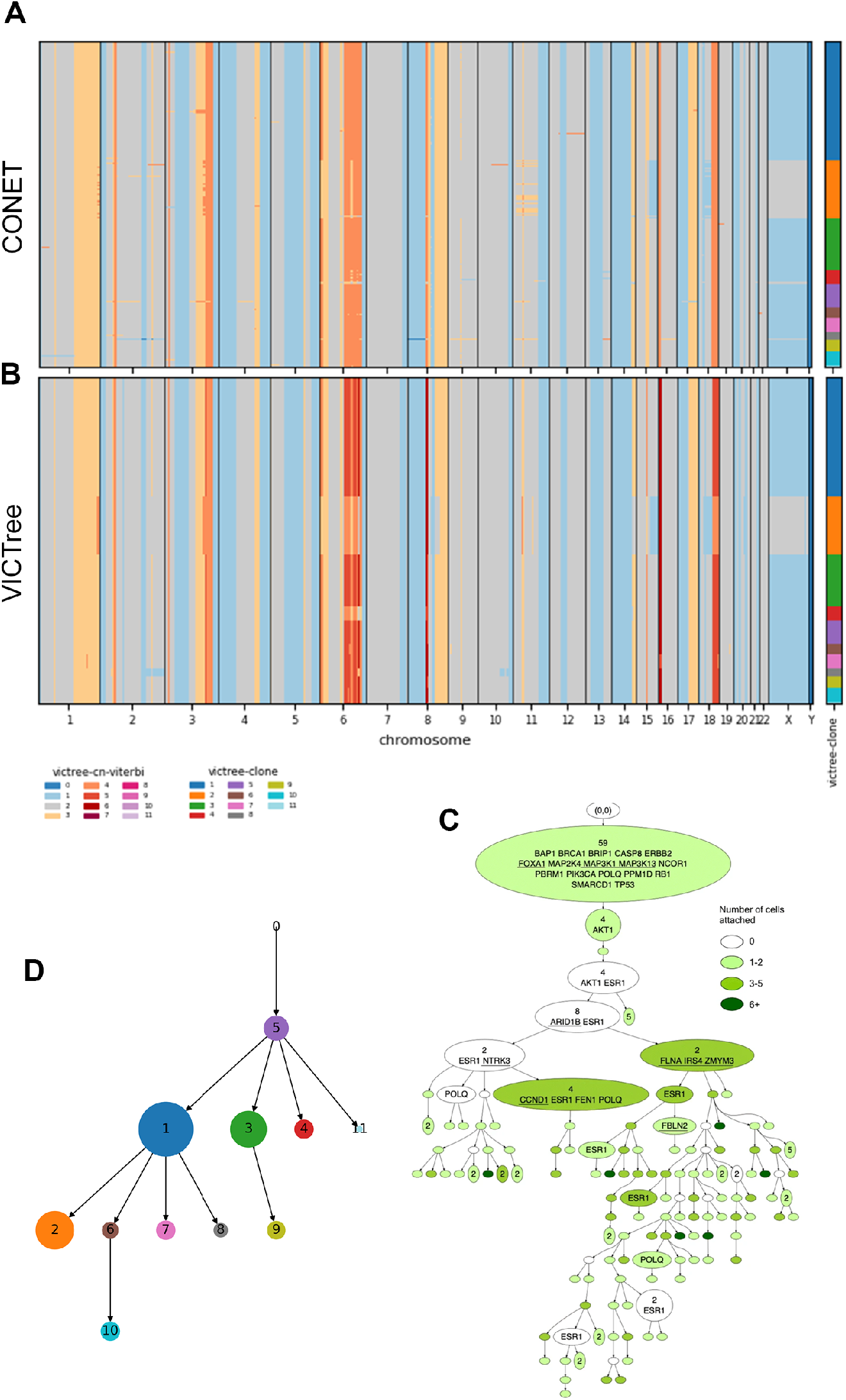
Sample of xenograft breast cancer SA501X3F. Features 260 single cells for 18175 bins of width 150kb. **A** CONET copy-number calling (sorted and annotated by VICTree clustering). **B** VICTree clonal copy-number calling and clustering of cells annotated on the right. **C** MAP tree from running VICTree with *K* = 14, where each nodes has size proportionally to the clone size according to VICTree result, and color matching the legend on the plots above. **D** CONET tree as shown in [25].

**Fig. S4:**
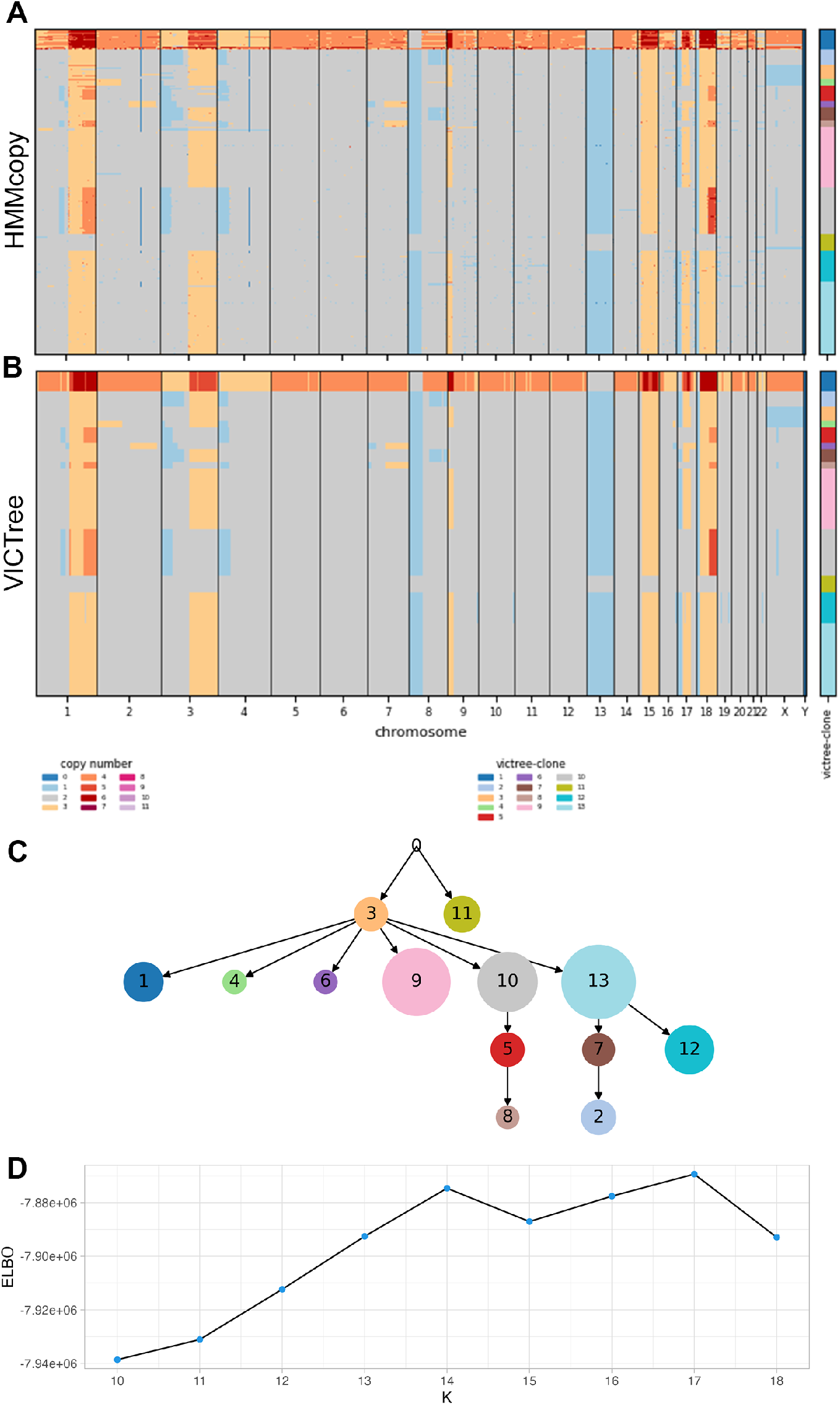
Sample of multiple myeloma MM03. Features 2208 single cells for 5298 bins of width 500kb. **A** HMMcopy copy-number calling. **B** VICTree clonal copy-number calling and clustering of cells annotated on the right. **C** MAP tree from running VICTree with *K* = 14, where each nodes has size proportionally to the clone size according to VICTree result, and color matching the legend on the plots above. **D** Plot of the ELBO for runs with several values of *K*.

**Fig. S5:**
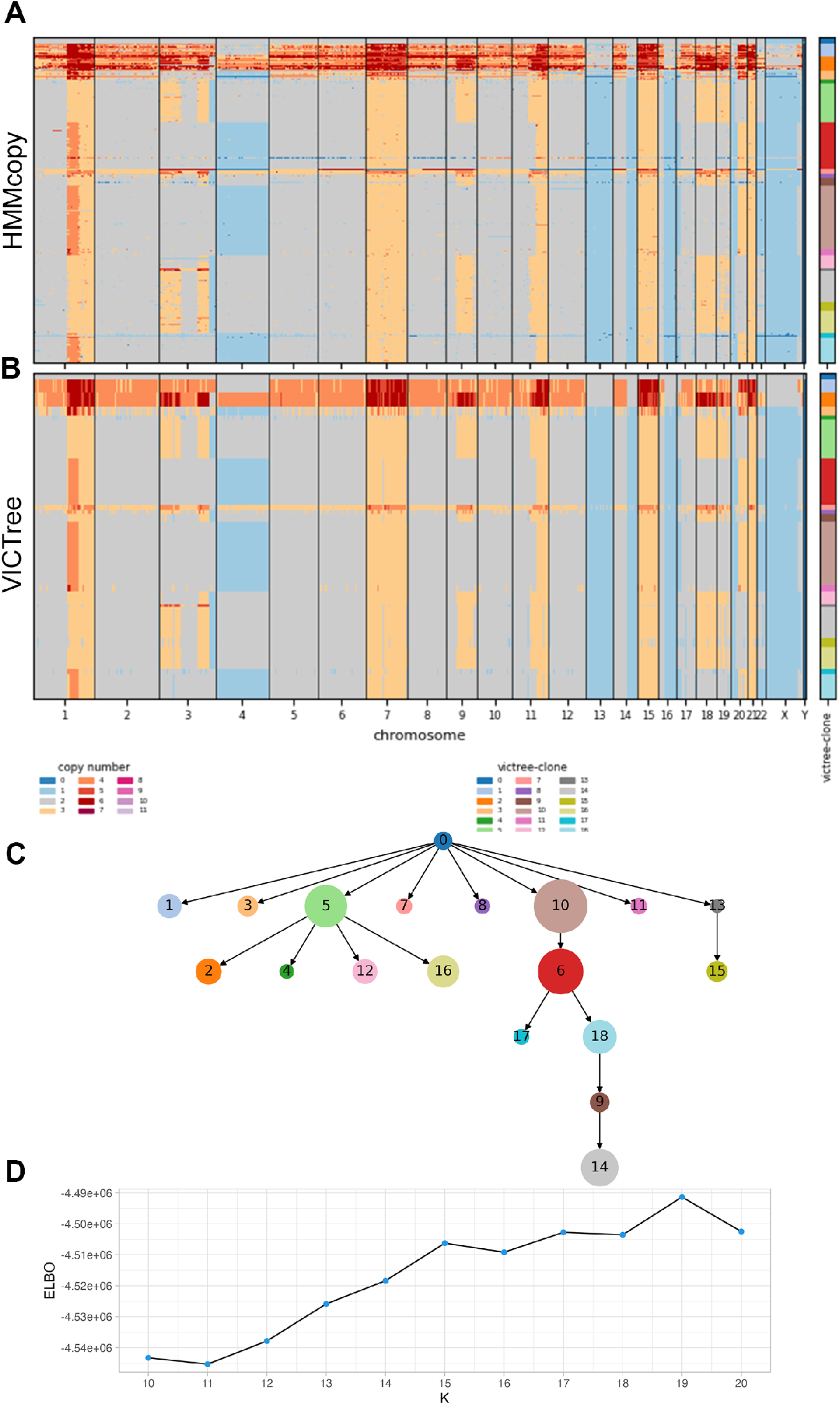
Sample of multiple myeloma MM04. Features 1003 single cells for 5286 bins of width 500kb. **A** HMMcopy copy-number calling. **B** VICTree clonal copy-number calling and clustering of cells annotated on the right. **C** MAP tree from running VICTree with *K* = 19, where each nodes has size proportionally to the clone size according to VICTree result, and color matching the legend on the plots above. **D** Plot of the ELBO for runs with several values of *K*.

**Fig. S6:**
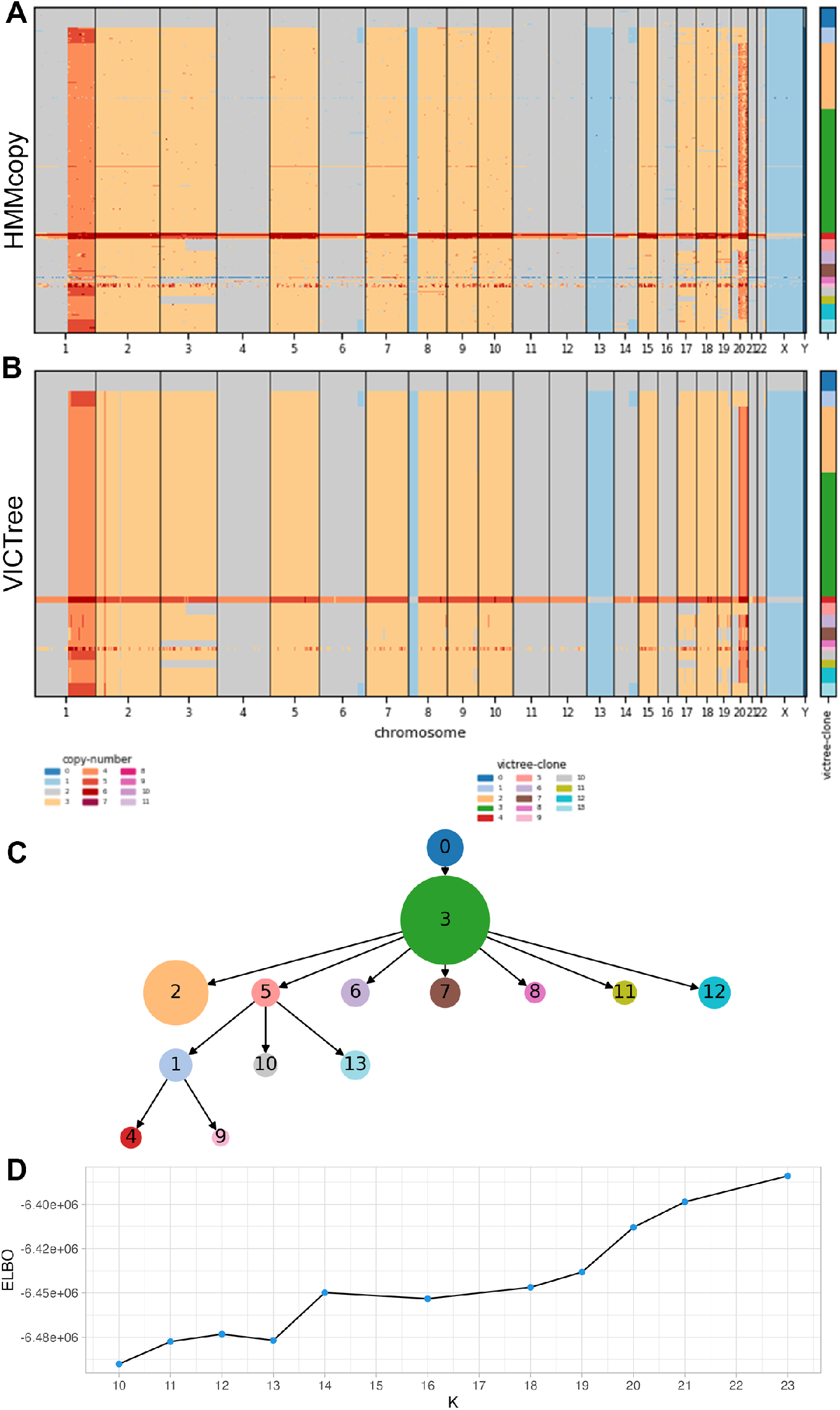
Sample of multiple myeloma MM29. Features 1616 single cells for 5285 bins of width 500kb. **A** HMMcopy copy-number calling. **B** VICTree clonal copy-number calling and clustering of cells annotated on the right. **C** MAP tree from running VICTree with *K* = 14, where each nodes has size proportionally to the clone size according to VICTree result, and color matching the legend on the plots above. **D** Plot of the ELBO for runs with several values of *K*.

**Fig. S7:**
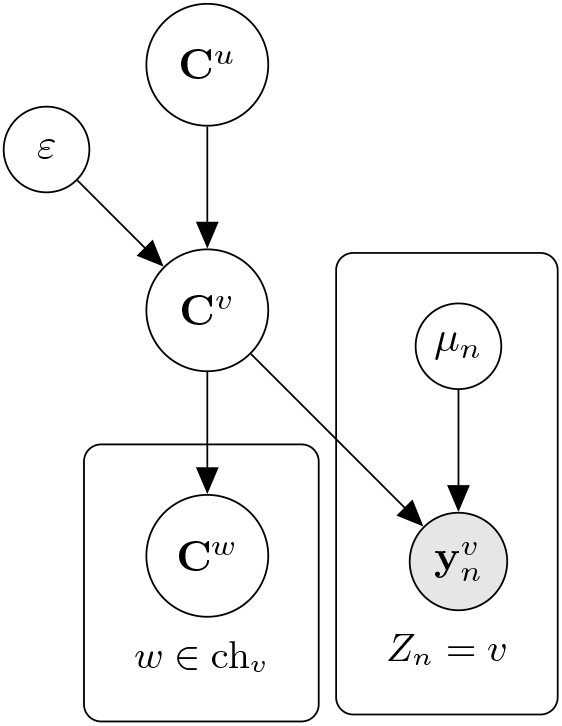
Simple copy-tree Bayes graph representing the neighborhood of a node *v* of the tree. For a generic node *v*, **C**^*v*^ is the latent copy number sequence, 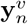 is the sequence of observations for each cell *n* in node *v, ε*^*v*^ the co-mutation parameter and *μ*_*n*_ is the baseline (per-copy expression).

### S2 CopyTree in exponential form

In the following sections we prove that CopyTree model belongs to the exponential family of distributions. We first introduce the notation by showing how standard non-homogeneous HMMs can be written in exponential form and then we extend the proof for the tree-structured HMM which we call CopyTree. The derivations in this sections will also be useful for Section S3 and Section S4 since the notation is first introduced here.

#### S2.1 HMM in exponential form

Let *M* be the length of a discrete Markov chain and *Σ* be the finite discrete set of values for the Markov chain states (alphabet). Let ***X*** be the |*Σ*| *× M* matrix representing a sequence **C** where each column is the one-hot-encoding of a state, i.e.

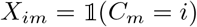

Also, we introduce two mappings which are useful for parameters vectorization:

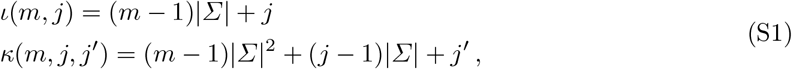

with *j, j*^*′*^ taking values in *Σ*. Note that the vectors are linear and quadratic functions of the alphabet length.

Let us denote with *x*_*m*_ the *m*-th column of *X*, which is the one-hot-encoding for the state at site *m*, and with *Y*_*m*_ the vector of observations for site *m*. Then we can write the joint probability as

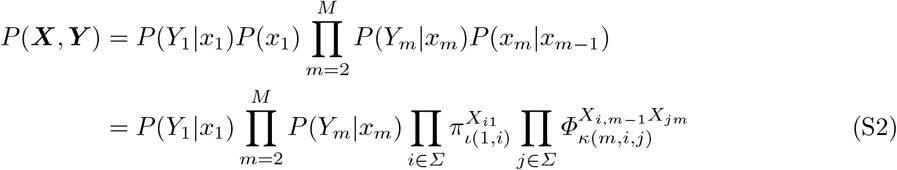

where ***π*** and ***Φ*** are the initial and transition probability vectors respectively, with indices provided by the mapping just described ^6^. For simplicity, we can define the pairwise state column vector 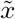 which is indexed by *κ* and has values given by

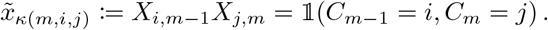

We can then re-write part of (S2) as exponential functions of the parameters.

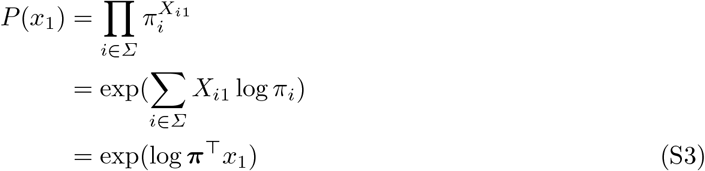

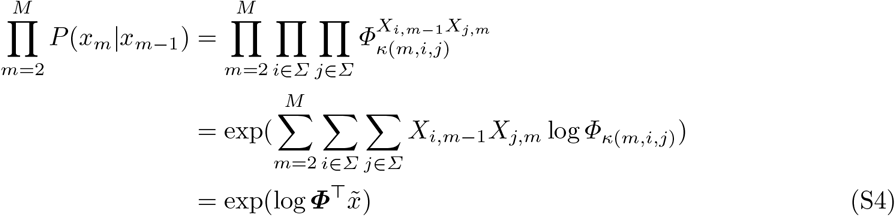

where log ***π*** := (log *π*_1_, .., log *π*_|*Σ*|_). Thus, all together in matrix notation:

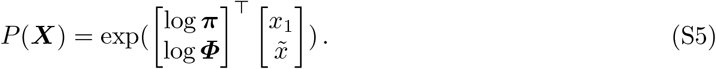

##### Definition 1

**(Exponential family form)**.

*Let X be a random vector whose distribution belongs to the exponential family and θ denote its parameters vector. Then the density function of X is*

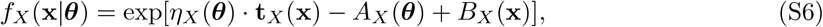

*where* **t**_*X*_ (**x**) *is the sufficient statistic vector, η*_*X*_ (***θ***) *is the natural parameter vector, A*_*X*_ (***θ***) *is the log partition and B*_*X*_ (**x**) *is the log base measure*.

If we assume that the emission probability can be written in exponential form (e.g. product of Gaussians with mean defined by **X**) and that it is conjugate to *P* (**X**), we can write *P* (**Y**|**X**) in terms of **t**_*X*_ (**X**)

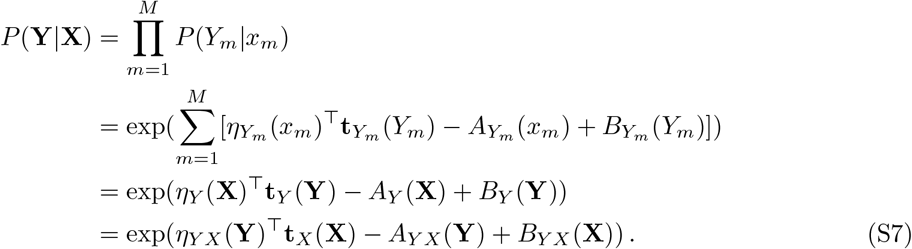

This reparametrization is also shown in [33] when presenting the Conjugate-Exponential models. It is worth noting that *P* (**Y**|**X**) can be thought as a contribution to the likelihood of *X*. Note that **t**_*X*_ is a concatenation of the flattened version of **X** and the 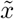 vector previously introduced. The natural parameters vector *η*_*Y X*_ (**Y**) depends on the distribution posit on the observations. More specifically, *η*_*Y X*_ (**Y**)_*ι*(*m,i*)_ = log *p*(*Y*_*m*_|*C*_*m*_ = *i*), function of *Y* with fixed *C*_*m*_ as it is multiplied by *X*_*i,m*_, therefore “activated” or “deactivated”.

Putting altogether, after reparametrizing as in (S7) and rearranging the terms at the exponent, we obtain the complete formulation

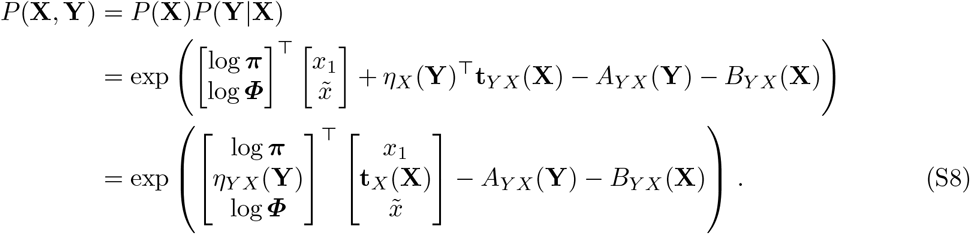

Since *η*_*Y X*_ (**Y**) needs to match the length of **t**_*X*_ (**X**) i.e. *M×*|*Σ*|+*M×*|*Σ*|^2^, we set *η*_*Y X*_ (**Y**)_*κ*(*m,j,j′*_) = **0**. We further notice that *x*_1_ is just the first column of ***X***, hence the first |*Σ*| elements of the flattened vector **t**_*X*_ (***X***). This means that we can group *x*_1_ by incorporating log ***π*** with *η*_*Y X*_ (***Y***). We split the sufficient statistic vector into *t*_1_(**X**), *t*_2_(**X**) and the natural parameter vector into *η*_1_(**Y, *θ, π***), *η*_2_(***Φ***)

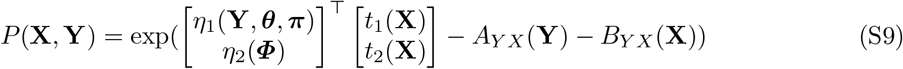

where ***θ*** is the set of extra parameters required in the emission distribution defined on the observations **Y**.

#### S2.2 CopyTree in exponential form

We will now refer to the variables using their biological meaning e.g. denoting by “copy-number profile” the latent states of the Markov chain, but the derivations are context independent and can be applied to any such model. The purpose of this section is to show how the conditional probability in each node of the Bayesian graph Figure S7 can be expressed as a function of the central node *v*’s sufficient statistic. These manipulations, besides allowing for Variational Message Passing [33], better explain the variational updates in section S3.

##### Root node

Let’s denote with **C**^*r*^ the copy number sequence of the root node and with **Y**^*r*^ the observations to it related. Since this does not depend on any other copy number sequence, we say that (**C**^*r*^, **Y**^*r*^) *∼* HMM(***ϕ, ψ***) with ***ϕ*** = {***π, Φ***} being the Markov chain parameters (initial state probability vector and transition matrix), and ***ψ*** being the set of parameters for the chosen emission distribution, whatever it may be.

Following the derivations from the previous section, the joint density function of **C**^*r*^, **Y**^*r*^ can be written in exponential form as

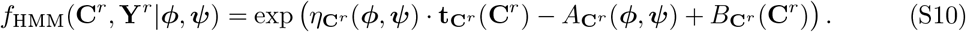

For a more readable notation we simply specify the node to which the parameter is related, which implicitly sets the right variables upon which a certain function depends. E.g. 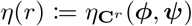. Equation S10 can be therefore rewritten as

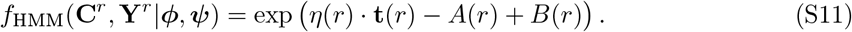

Both the sufficient statistic and the natural parameter vectors are easily represented as the concatenation of two vectors

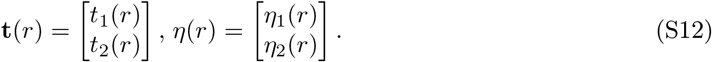

In subsection S2.1 we show that the vectors’ elements are as follows

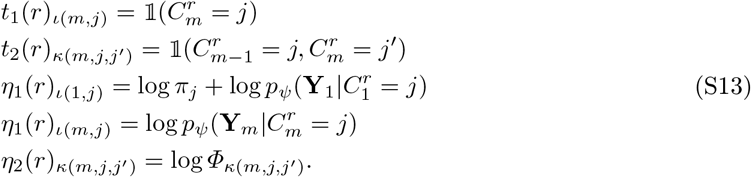

##### Tree HMM joint

We write the root node *r* HMM probability as the standard HMM joint since it does not have a parent node.

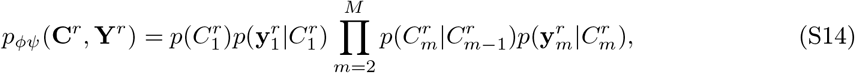

Then, for any internal node *u* with parent *v* we get the probability of the HMM of a child *v* of *u*, given the parent as

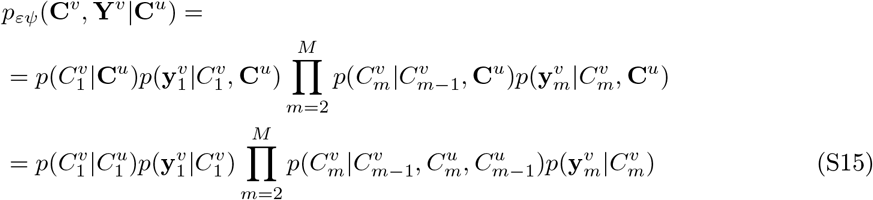

where in step (S15) we assume that the transition probability of the child only depends on the two copy numbers of the parent at bin *m −* 1, *m*.

The complete joint probability *p*_*ϕψε*_(**C, Y**) should be derived from root to leaves given the two quantities above in a chain product.

##### Internal copy-number node

For an internal node of the tree, we introduce the CN coherence function, defined over each single copy number of the sequence. For any node *v* and its parent *u*, the CN coherence function *h* over the copy number values at a specific bin *m* is defined as in (4).

For the first element of the sequence we simply redefine *h* to be

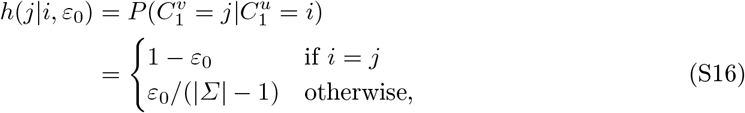

In the context of a variational message passing approach [33], it can be useful to draw a simple graph encoding the dependencies (see Figure S7).

The copy number sequences follow a Markov model. Let *u, v* the parent-child pair of node, as usual, and let ***X***^*u*^, ***X***^*v*^ be the copy number sequences of the respective nodes in the one-hot encoding described above.

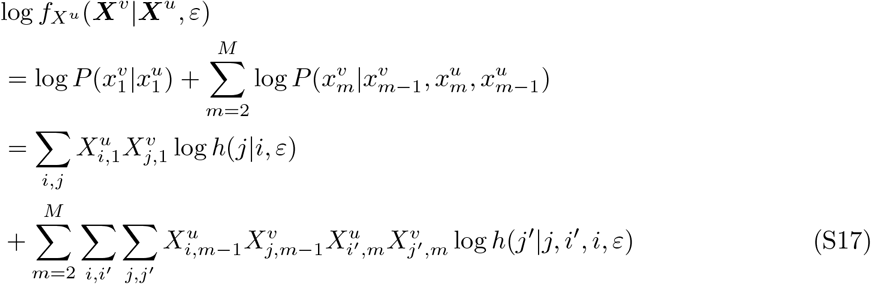

In subsection S2.2 we have decomposed the sufficient statistic vector for HMMs in

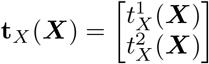

with 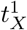 specifically, and 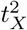 being two column vectors indexed by *ι* and *κ* mappings respectively. More

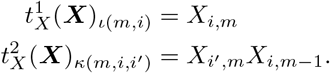

With that in mind, we regroup (S17) to obtain the natural parameter vector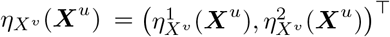, which is then equal to

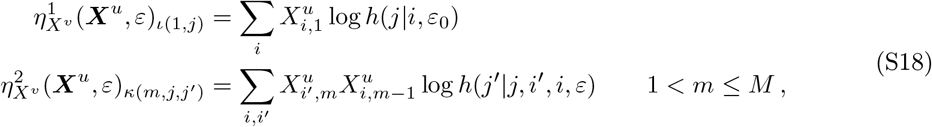

and the rest of the vector elements, namel 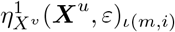 for 1 *< m ≤ M* and 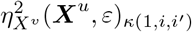, are set to zero.

We can now write the conditional probability in (S17) as

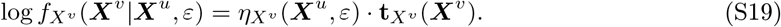

It is then useful to rewrite this as a function of the sufficient statistic of the parent node’s copy number, so that the rest can be used as a message going from child to parent (see [33]). Precisely, we want to find the vector *η*_*Xv*_ *X*_*u*_ function of ***X***^*v*^ such that

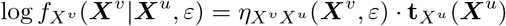

By symmetry of (S17), and since **t**_*Xu*_ has the same form of **t**_*Xv*_ we obtain 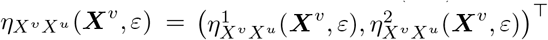 such that

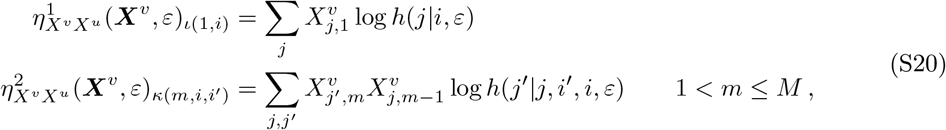

and the rest of the elements equal to zero. Note that even though the presented CN coherence function is symmetric, (S18) and (S20) might differ from the indices of *h* when it is not symmetric on the pairs (*i, i*^*′*^) and (*j, j*^*′*^), therefore in general we have *η*_*Xv*_ ≠ *η*_*Xv*_ *X*_*u*_.

##### Observations

Let ***Y*** ^*u*^ be the set of observation at node *v*. For each bin *m* we have a collection of reads, one for each cell assigned to the node and for each gene belonging to the bin. We denote this collection of reads with 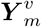.

To simplify the model, we assume that each of these reads range from 1 to *N*, independently of the cell/bin they belong to, and we assume that they are drawn from a Normal distribution, conditioned on the bin’s copy number, with mean 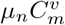.

The conditional log-distribution is then

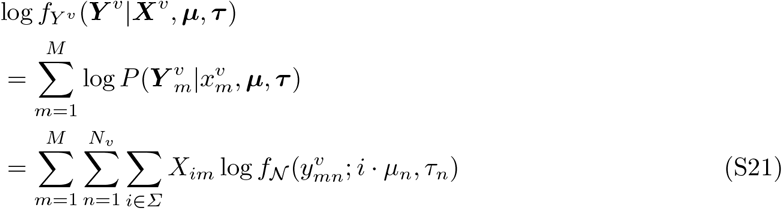

which can be expressed in terms of **t**_*Xv*_ (***X*** ^*v*^)

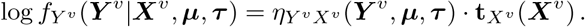

where **t**_*Xv*_ is the same as in (S2.2) and

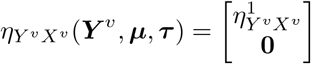

with **0** null vector of length matching 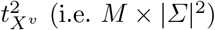 and 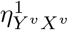 with same length as 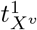, and elements such that

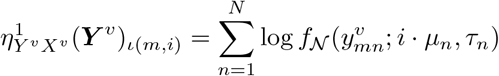

### S3 Variational Updates

In this section we go through the derivations of the CAVI update for each variational distribution. All updates make use of the CAVI update equation (3).

#### S3.1 Copy numbers

##### Copy numbers pre-requisite

We begin by observing that our model for a given *T*, when described as a directed graphical model on the variables

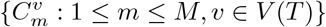

has a cyclic underlying undirected graph (see 2a).

However, it can be formulated as a directed graphical model on the variables

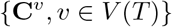

where *p*(**C**^*v*^|**C**^*ρ*^_*T*_ ^(*v*)^) is a conditional Markov model such that the underlying directed graphical model is a tree. In order to draw this conclusion we consider the conditional joint probability of observations and copy numbers

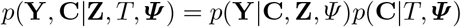

and show how to express the above conditional joint as a product of conditional Markov models:

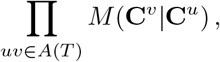

where the conditional Markov models depend on **Z, *Ψ***, **Y**. Notice that

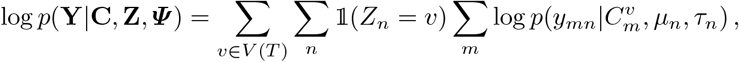

and that

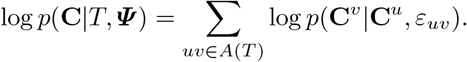

This implies

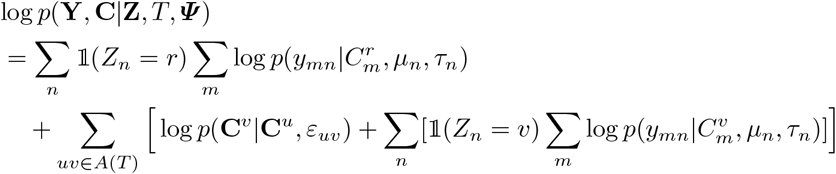

Hence, it is sufficient to find a conditional Markov model *M* (**C**^*v*^|**C**^*u*^) for which the product of sufficient statistics and natural parameter is the outermost summand of the expression above, that is:

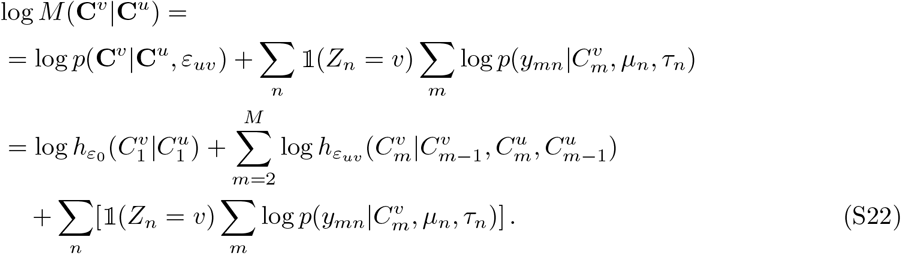

The following shows how to express the Markov model for *M* (**C**^*v*^|**C**^*u*^) in its exponential family form, that is, using the natural parameter ***η***_*T*_ (*v*) and the sufficient statistics **t**_*T*_ (**C**^*v*^). Keep in mind that the sufficient statistics is a function of the copy number sequence, while the natural parameter depends on other parameters e.g. ***Ψ***. Let the parent node of *v* given the tree *T* simply be *ρ* = *ρ*_*T*_ (*v*), and the quantities ***η***_*T*_ (*v*), **t**_*T*_ (**C**^*v*^) be just ***η***(*v*), **t**(**C**^*v*^), then the decomposition of the latters can be expressed in the following way.

Referring to the mappings in (S1), similarly to the notation introduced in section S2, the sufficient statistics vector **t**(**C**^*v*^) is equal to

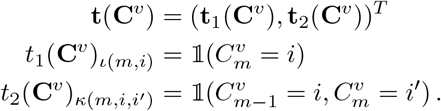

Natural parameters, ***η***(*v*), and associated parameters, ***η***^*′*^(*v*):

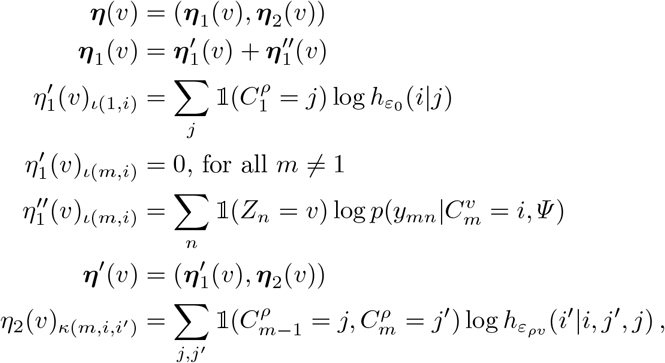

where both ***η***(*v*) and ***η***^*′*^(*v*) are vectors of dimension |*Σ*| *× M* + |*Σ*|^2^ *× M*, and match the dimension of **t**(**C**^*v*^).

Notice that, as desired, the inner product of such natural parameters with sufficient statistic vectors, matches the Markov model in (S22):

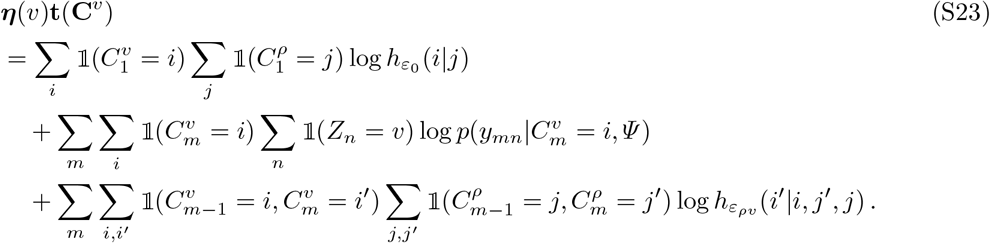

Moreover,

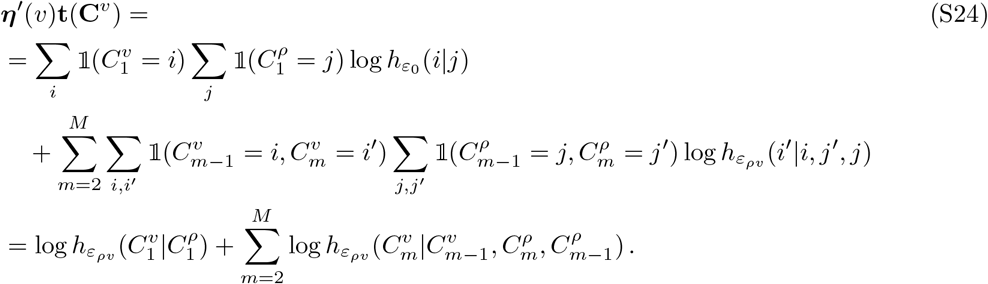

This should clarify the purpose of the natural and associated parameters: while the latter embodies the attributes of the Markov model over the copy number sequence, the former also links the sequence to the observations.

##### Copy numbers update

We begin by noticing that 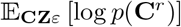 is uniquely determined by the prior that has probability 1 for the reference genome which typically has copy number 2 everywhere. For the the remaining *K −* 1 vertices we provide the details for the standard CAVI updates. Let us first consider a fixed tree *T* and any node *v≠ r*.

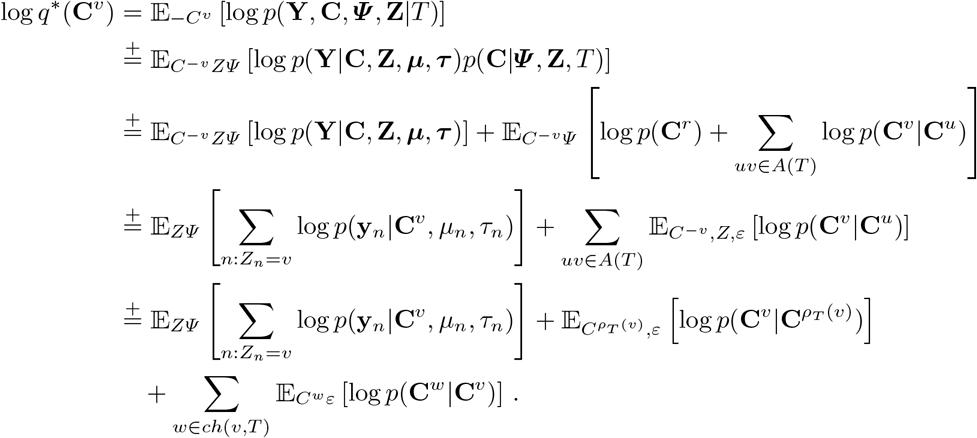

Where *ρ*_*T*_ (*v*) is the parent and ch(v, T) is the set of children of vertex *v* in tree *T*. Moreover, as shown above,

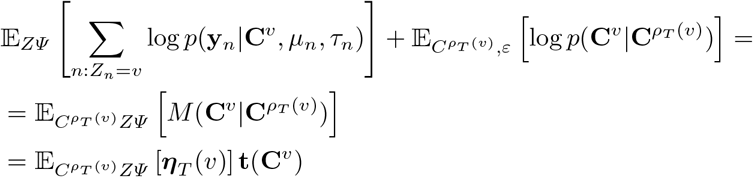

and, which requires some manipulations,

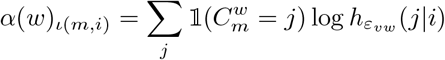

where

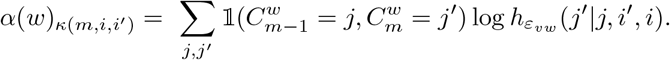

and

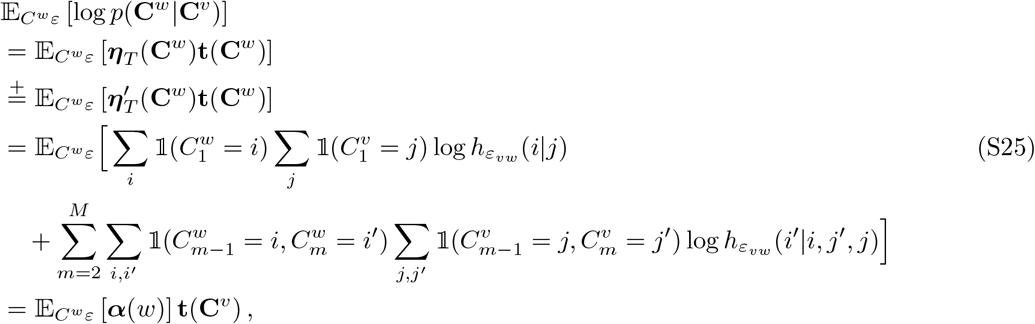

Note that the above two expressions have *j* and *i* inverted in *h*(*·*) because we express the indexing with *i*, but the copy numbers indexed by *j* (which belong to node *w*), are named after *i* in (S25).

Consequently,

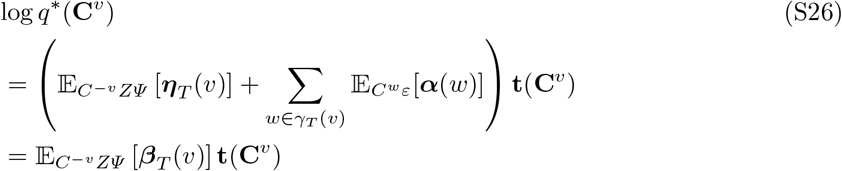

where ***β***_*T*_ (*v*) can be be expressed as the concatenation of two vectors, i.e. *β*_*T*_ (*v*) = (*β*_*T*,1_, *β*_*T*,2_), where *β*_*T*,1_(**C**^*v*^) and *β*_*T*,2_(**C**^*v*^) are defined as follows

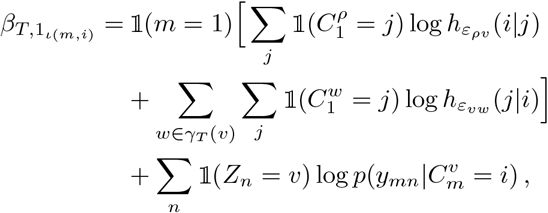

and

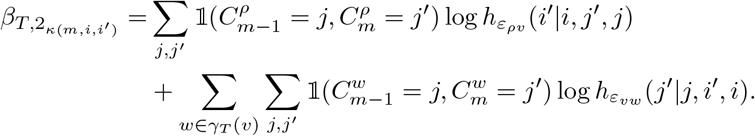

Finally, when considering the VI-distribution over trees *q*(*T*), rather than a fixed tree, we obtain the final update

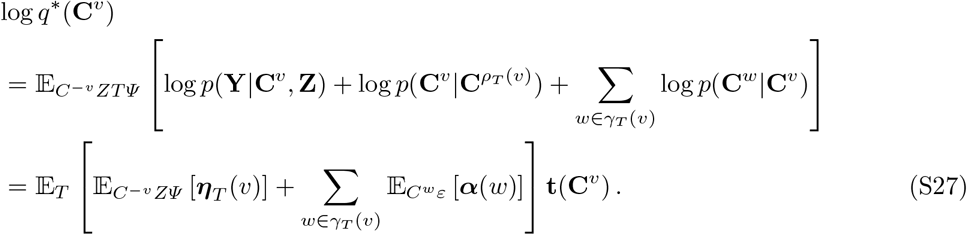

Therefore, we update the natural parameter as

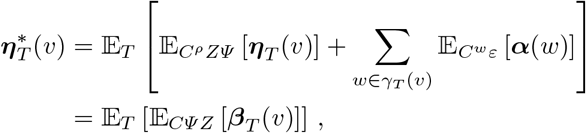

which can be decomposed, element by element, in

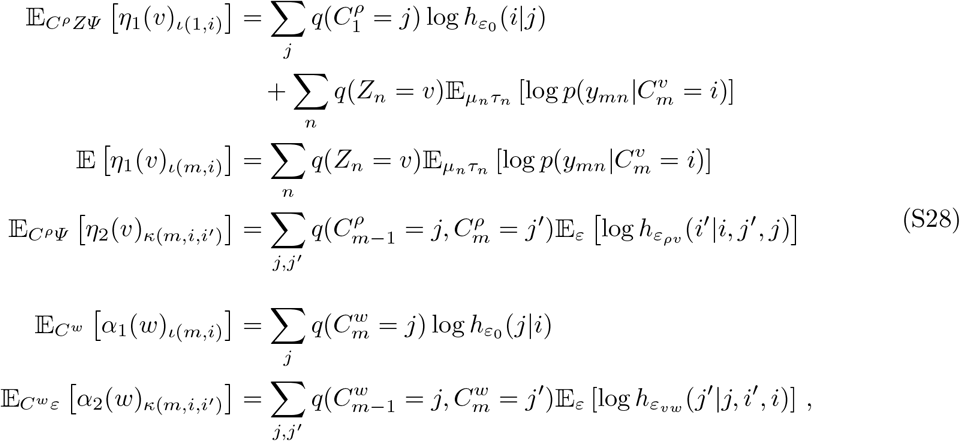

Where

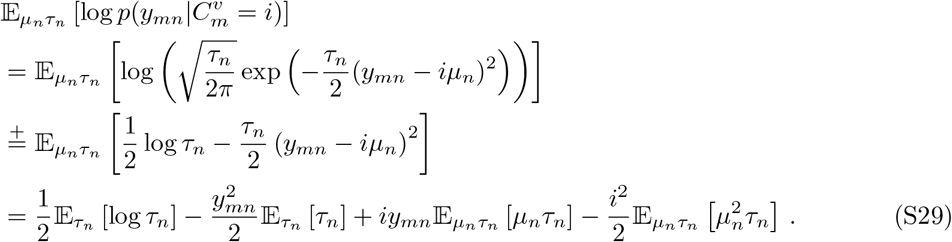

Given that the variational distribution over *μ*_*n*_, *τ*_*n*_ is Normal-Gamma with parameters

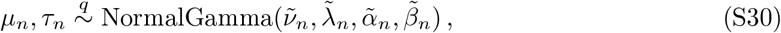

the expectations in the previous formula are equal to:

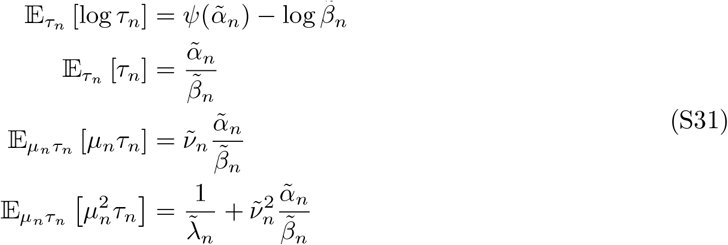

#### S3.2 Tree topology

We begin by using the mean-field VI standard update:

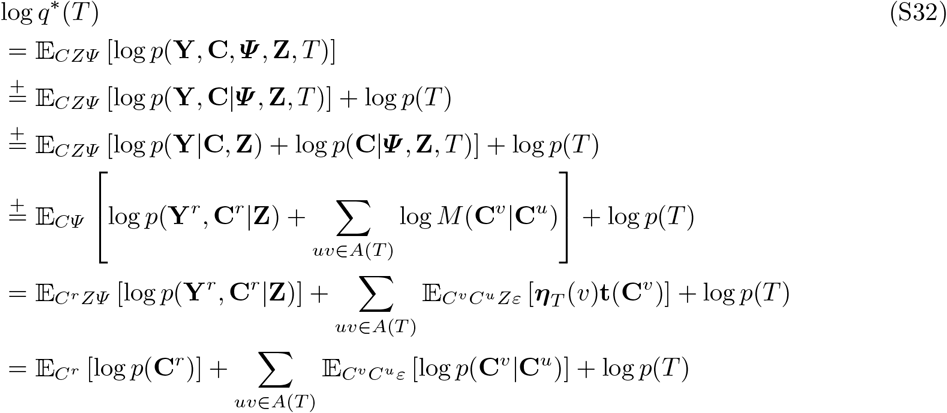

The expression in (S32) contains three terms that we further examine below.

First, 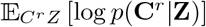, as stated above, is uniquely determined by the prior that has probability 1 for the reference healthy genome, which typically has copy number 2 everywhere. Furthermore, log *p*(*T*) is constant over the support of *T* as we assign a uniform prior.

What varies over *T* is the summand of the remaining term, 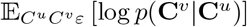, for which we use the notation *w*(*uv*) as it represents the unnormalized weight of each arc in the variational distribution. It can be further simplified as:

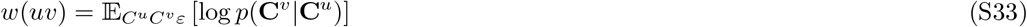

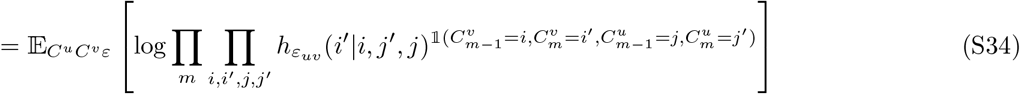

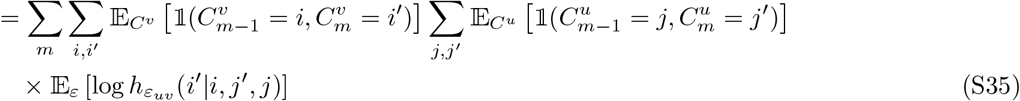

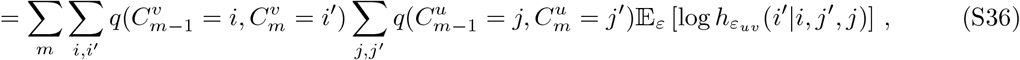

where 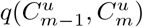 can be computed iteratively for all *m* with a single forward pass, which is all performed in *O*(*KMA*).

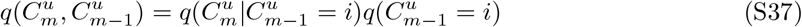

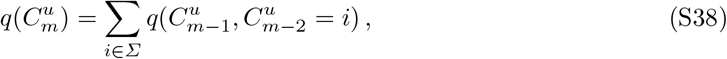

and the expectation of the CN coherence function is piecewise defined as

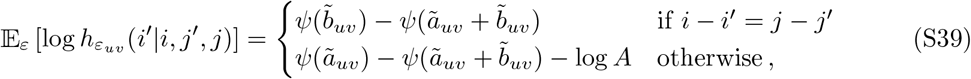

following the expression in (4) and the fact that 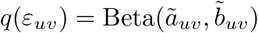.

Note that the weight *w*(*uv*) is the same for all trees that contains the edge *uv*. We use the weight matrix over the fully connected graph, *W* (*𝒢*), containing all *w*(*uv*) : *uv ∈ 𝒢*, for sampling trees using 2 and evaluating our sampled trees on the variational distribution *q*(*T*).

#### S3.3 Arc distances

The update for *q*(*ε*_*uv*_) is derived as follows:

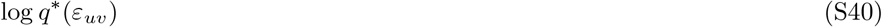

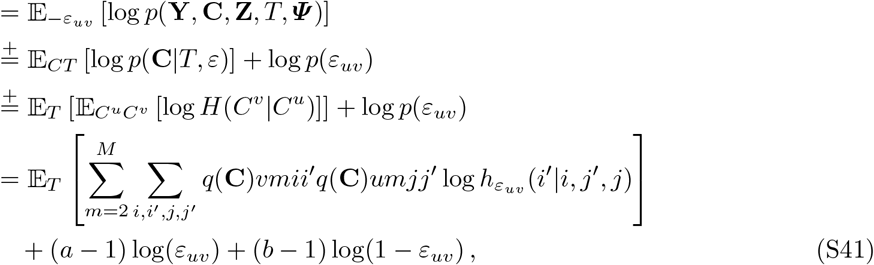

where 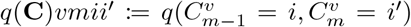. Furthermore, the summation over the four copy numbers can be split into two separate cases, i.e. the two cases of the CN coherence function, which we denote by A := {*i, i*^*′*^, *j, j*^*′*^|*i − i*^*′*^ = *j − j*^*′*^} and its complementary set *¬*A. This being said, we rewrite (S41) as

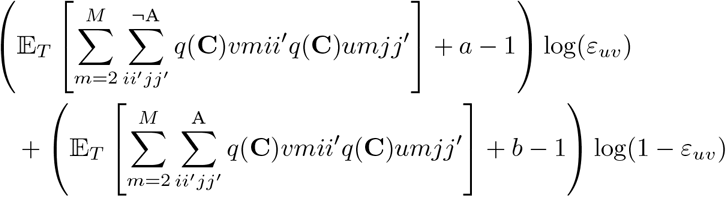

which leaves us with the updated parameters of the variational Beta distribution over *ε*_*uv*_:

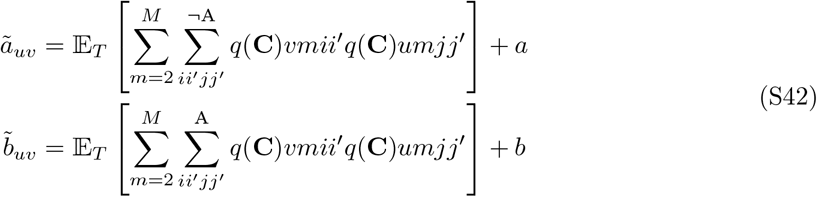

#### S3.4 Assignment concentration

Derivation of *q*(*π*):

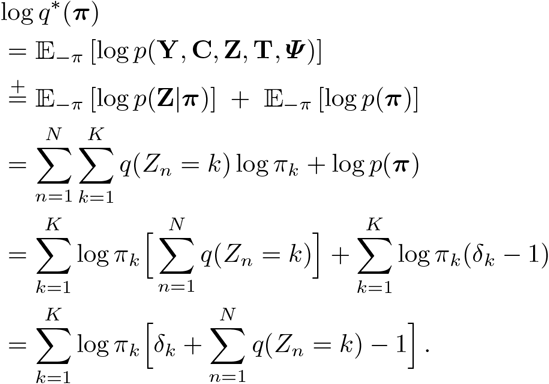

Therefore, *q*^*\**^(***π***) is a Dirichlet distribution with parameters 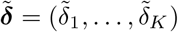, where

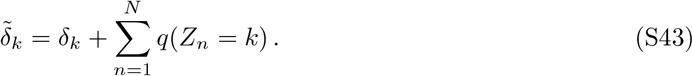

#### S3.5 Cell assignments

Derivation of *q*(*Z*_*n*_):

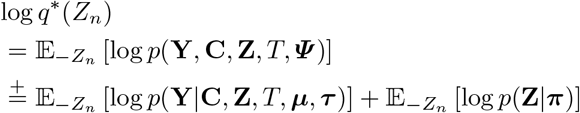

Note that log *p*(**Y**|**Z, C, *Ψ***) and log *p*(**Z**|***π***) can be written as the sum of two terms, one that depends on *Z*_*n*_ and one that does not, i.e.,

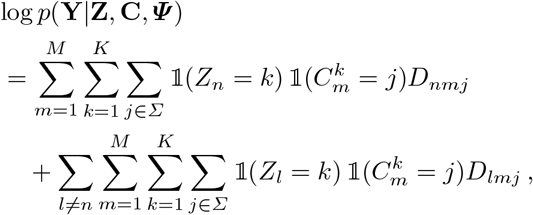

and

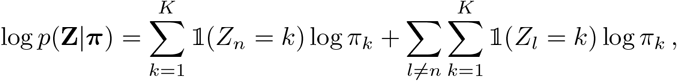

Where

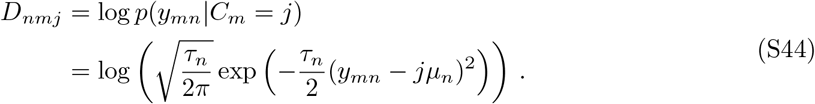

Consequently,

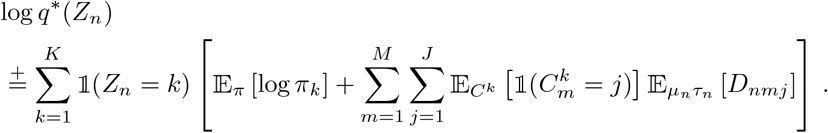

We simplify the expression further by:

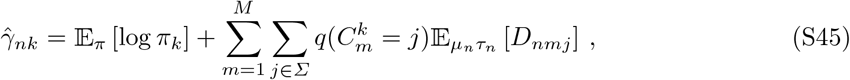

with 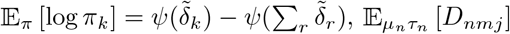 computed as in (S29). Finally, we set:

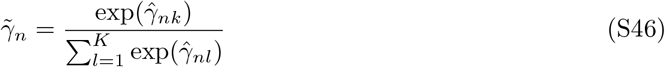

and conclude that *q*^*\**^(*Z*_*n*_) *∼* Categorical 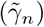 with parameters 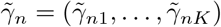

#### S3.6 Baseline mean and precision

We use the Normal-Gamma joint prior on *μ*_*n*_ and *τ*_*n*_, that factorizes over *n*, and introduce a joint variational distribution on the corresponding parameters. The mean-field standard update becomes:

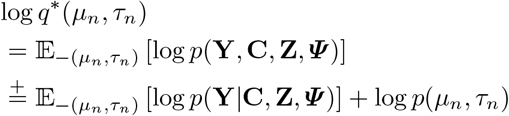

Simplifying each term individually yields:

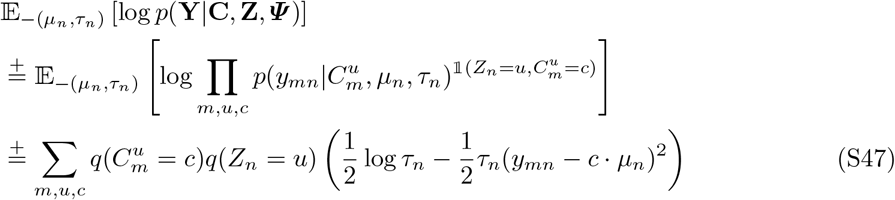

and

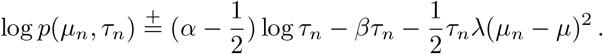

As our model prior is Gamma-Normal distributed, we suspect our variational distribution to be in the same family, and attempt to match the parameters associated with the independent variables of the Gamma-Normal. First we regroup the variables as follows:

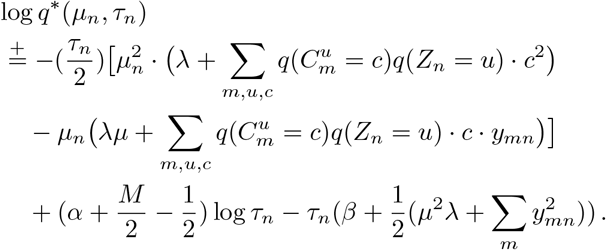

In order to get the expression on Normal-Gamma form we need to complete the square for the cross term. We do this by adding and subtracting the square of the factor of *μ*_*n*_*τ*_*n*_ and then multiplying it with *τ*_*n*_; the negative term is then passed to complete the square and the positive is added to the *τ*_*n*_ term:

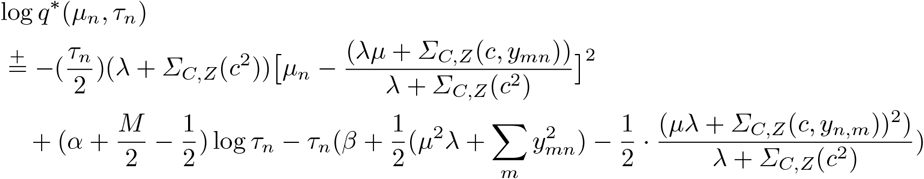

where

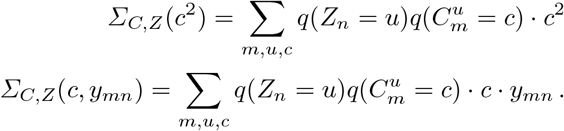

By matching the independent variables with the parameters we see that *q*^*\**^(*μ*_*n*_, *τ*_*n*_) is indeed Normal-Gamma 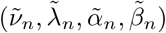 distributed with parameters:

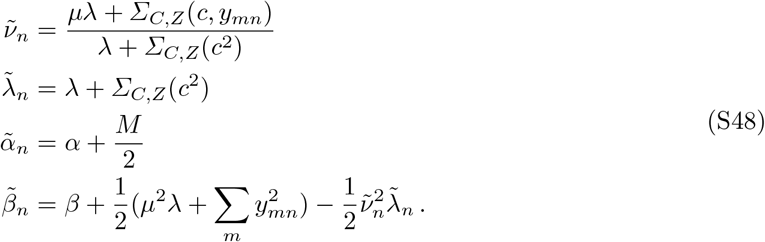

### S4 ELBO computation

In this section we derive the entropy and negative cross-entropy terms of 10 in 4.3.

Since the expectation over the tree topology *T* in 10 requires a sampling technique to be estimated, i.e., it cannot be derived as for the other variables in closed form, we separate the expectation into two main parts:

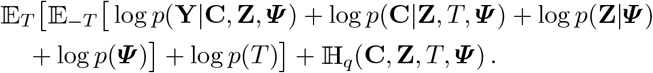

In the subsections below we present each component of the ELBO, by deriving the negative cross entropy terms of in the inner expectation and the variational distribution entropy, the latter which we write following the notation:

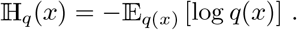

#### S4.1 Observations

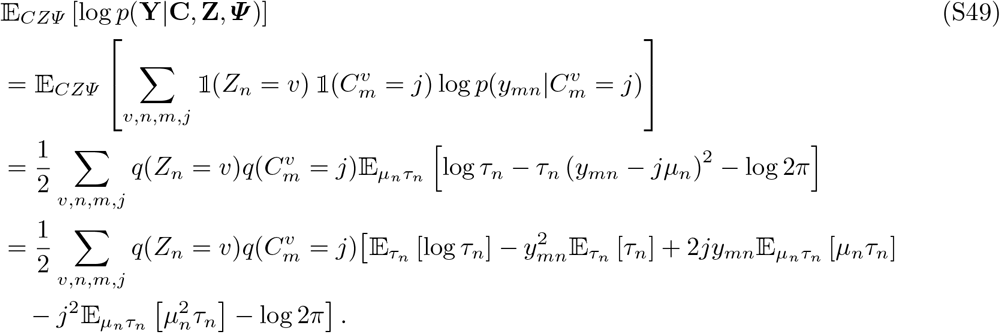

Note that the values of the expectations above are standard expectations for Normal and Gamma random variables and can be found in subsection S3.1.

#### S4.2 Copy numbers

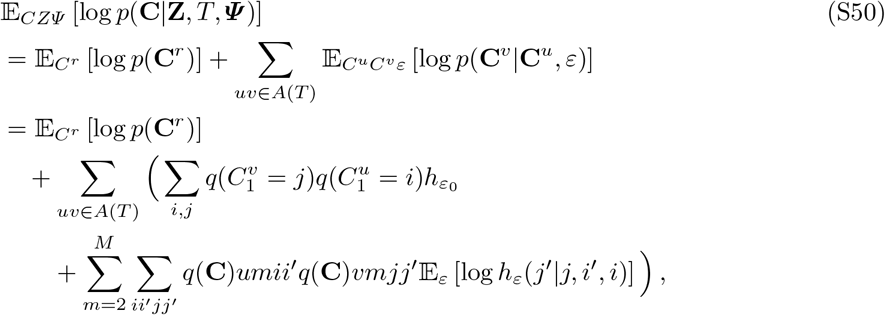

where 𝔼 _*ε*_ [log *h*_*ε*_(*j*^*′*^|*j, i*^*′*^, *i*)] is computed as shown in (S39) and the variational distribution quantities (denoted as in Equation (S41)) for the copy numbers are computed with the forward-algorithm. Furthermore, we can calculate the entropy by use of the marginals *q*(*C*_*m*_) in the following way:

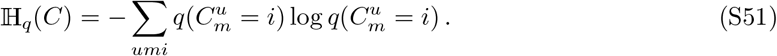

#### S4.3 Cell assignments

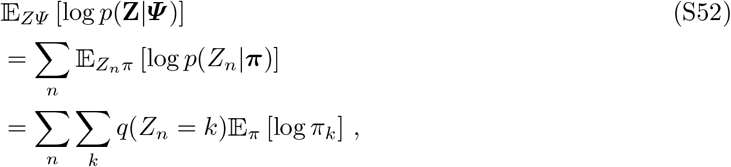

with 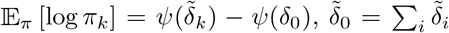 such that 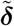 is the parameter of the variational Dirichlet distribution over ***π***.

The entropy is calculated as:

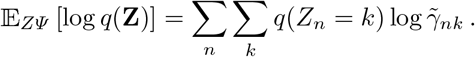

#### S4.4 Concentration parameter

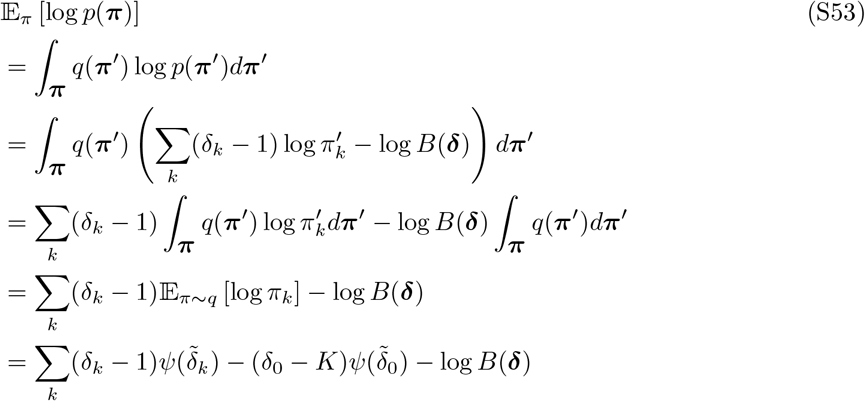

Where

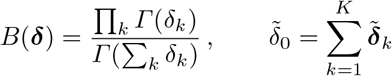

and 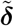 refers to the variational concentration parameter, while ***δ*** is the generative model parameter.

The entropy of *q*(***π***) is found by replacing the model parameter ***δ*** with 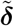.

#### S4.5 Gaussian mean and precision

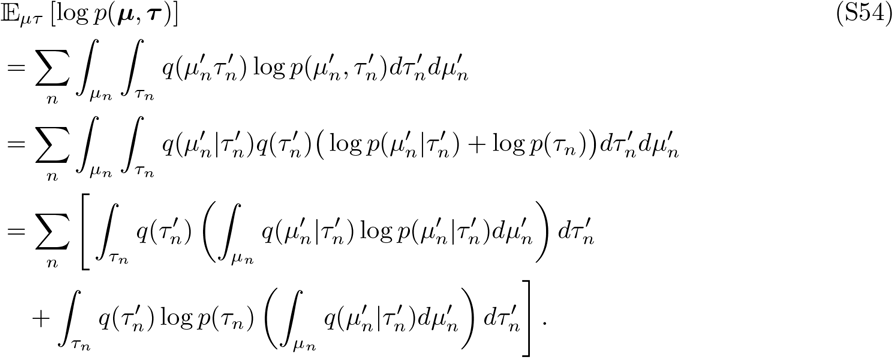

The above expression is made of two terms (excluding the outermost summation): the first double integral which is the expectation over *q*(*τ*_*n*_) of the negative cross-entropy 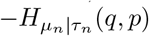, and the second integral which is just the negative cross-entropy over *τ*_*n*_ of *q* and *p*, i.e. 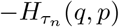.

Therefore we get

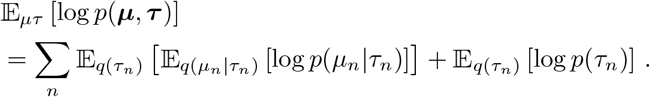

We compute the partial terms below (omitting *q* as all the expectations here are taken with respect to the variational distribution)

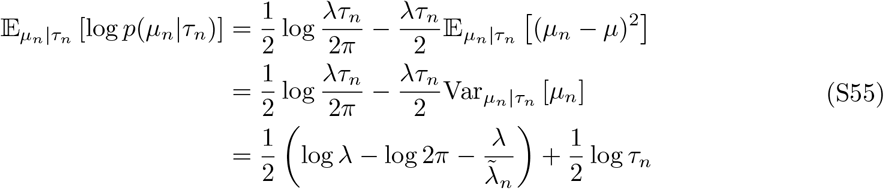

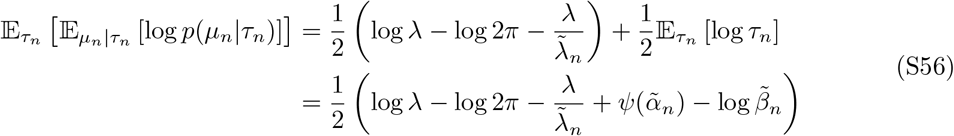

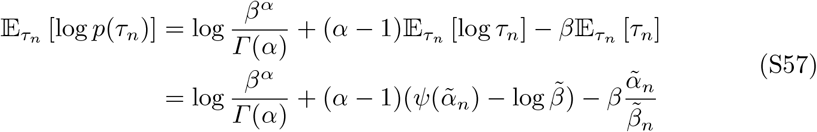

and then put altogether:

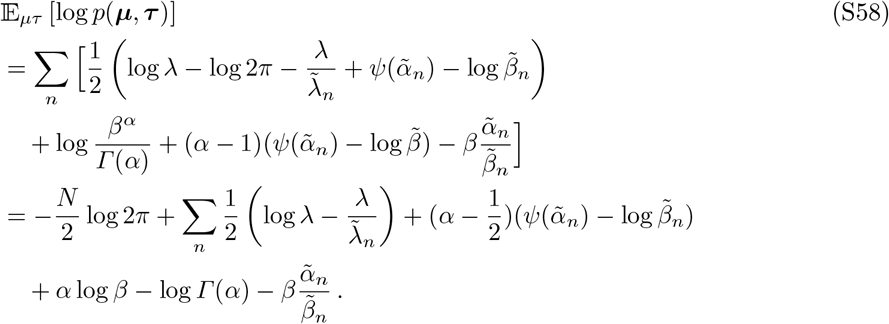

The variational entropy is computed as follows

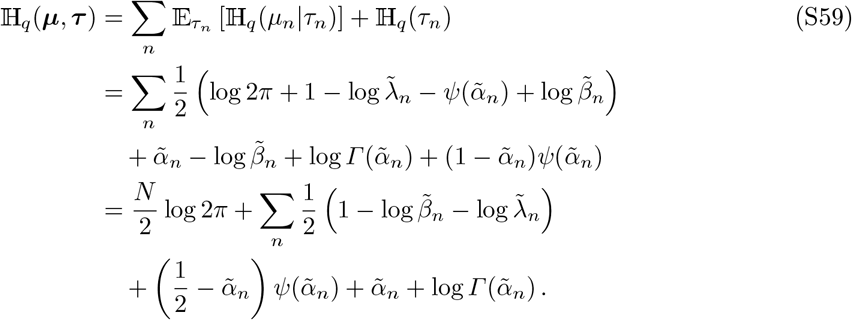

#### S4.6 Epsilon

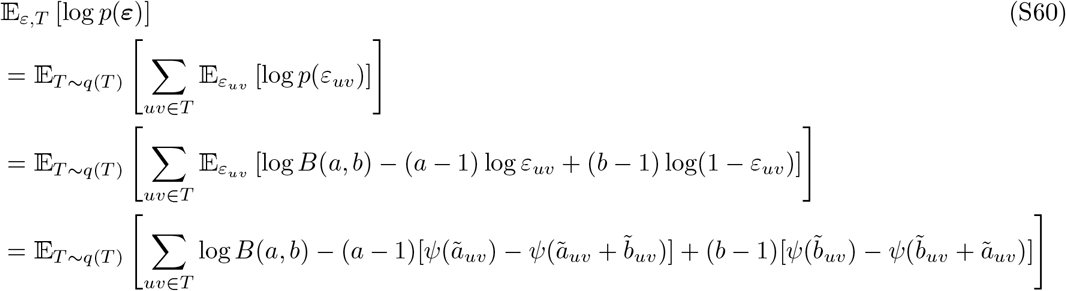

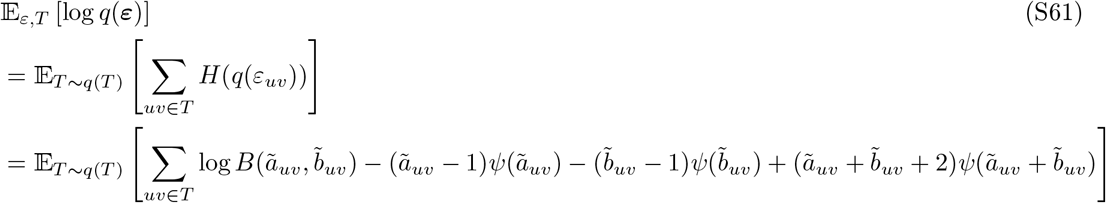

#### S4.7 Tree topology

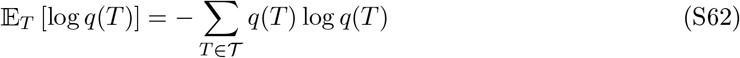

since *q*(*T*) is unnormalized, we use importance sampling to estimate the normalizing constant and evaluate on 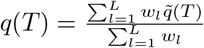

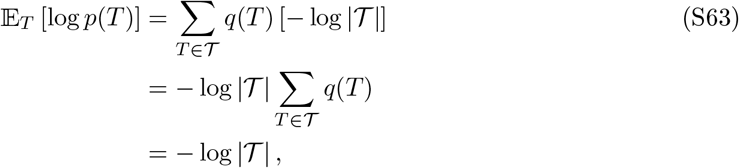

where in case of *K* nodes and considering labeled arborescences, | *𝒯* | = *K*^*K−*2^ [5].

### S5 VICTree

#### S5.1 VICTree pseudocode

The algorithm complexity is dominated by the update of *q*(**C**) parameters, which requires *O*(*KMNA*^2^), at every iteration and for every unique tree in the sample.

##### Algorithm 1

VICTree pseudocode

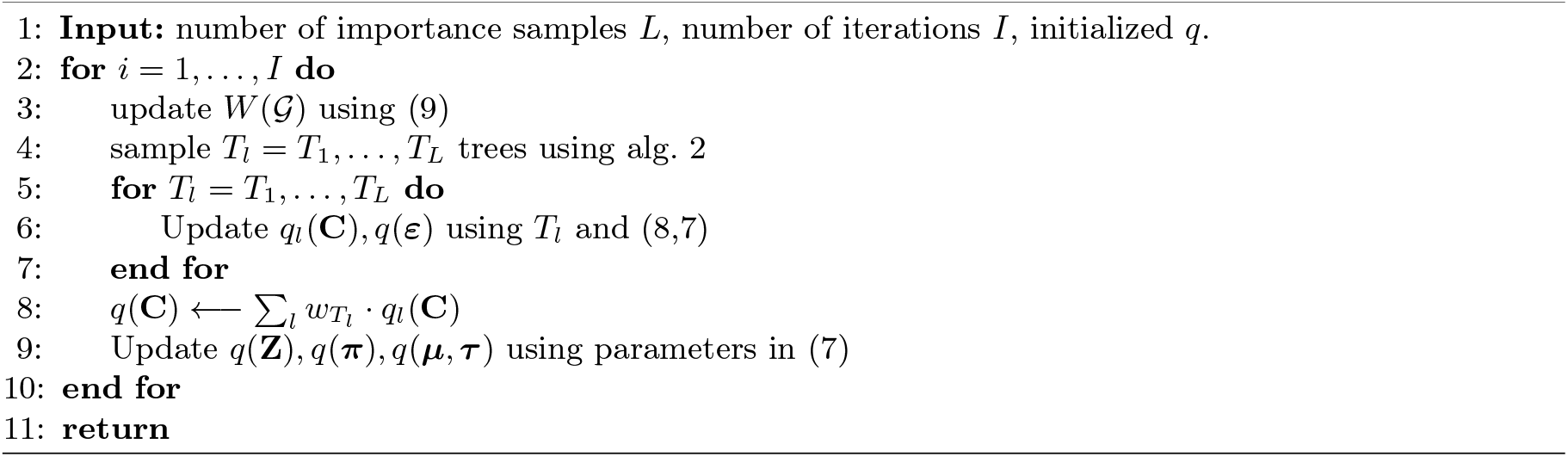

#### S5.2 VICTree Importance sampling details

During inference process, we need to evaluate expectations over *q*(*T*) of some function of the tree, but enumerating over all possible arborescences can be costly when considering directed labeled trees with several nodes. More specifically, by Cayley’s formula [5] and Prüfer sequences it is proven that the number of labeled trees with *n* nodes is *n*^*n−*2^. In our case the trees are rooted and directed, thus the topology space is of size *n*^*n−*1^, which quickly becomes intractable.

Assuming we can draw samples from *q*(*T*) distribution, we can approximate an expectation as follows.

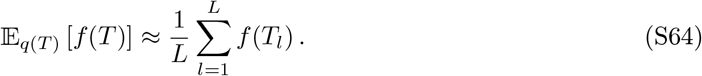

However, sampling from *q*(*T*) is not straightforward, therefore we sample from an alternative distribution *g*(*T*) and apply importance sampling [14] in order to evaluate the desired expectation. Let 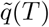 denote the weight of a arborescence *T*, which is defined to be the product over the edge weights and let 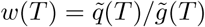 be the importance weights. Then

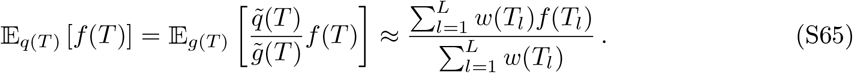

Self-normalization (S65) is required since both *q*(*T*) and *g*(*T*) are evaluated up to a multiplication factor.

### S6 LARS algorithm

The following algorithm runs with computational time *O*(|*E*|*K*^3^) since Edmond’s algorithm requires *O*(*K*^2^). It is then clear that, for bigger trees, the complexity of a tree sample quickly overcomes the complexity of the other costly operations in each VICTree step, especially the copy-number variational distribution update *q*(**C**) as mentioned in Section S5.1.

#### Algorithm 2

Labeled Arborescence Sampler (LARS)

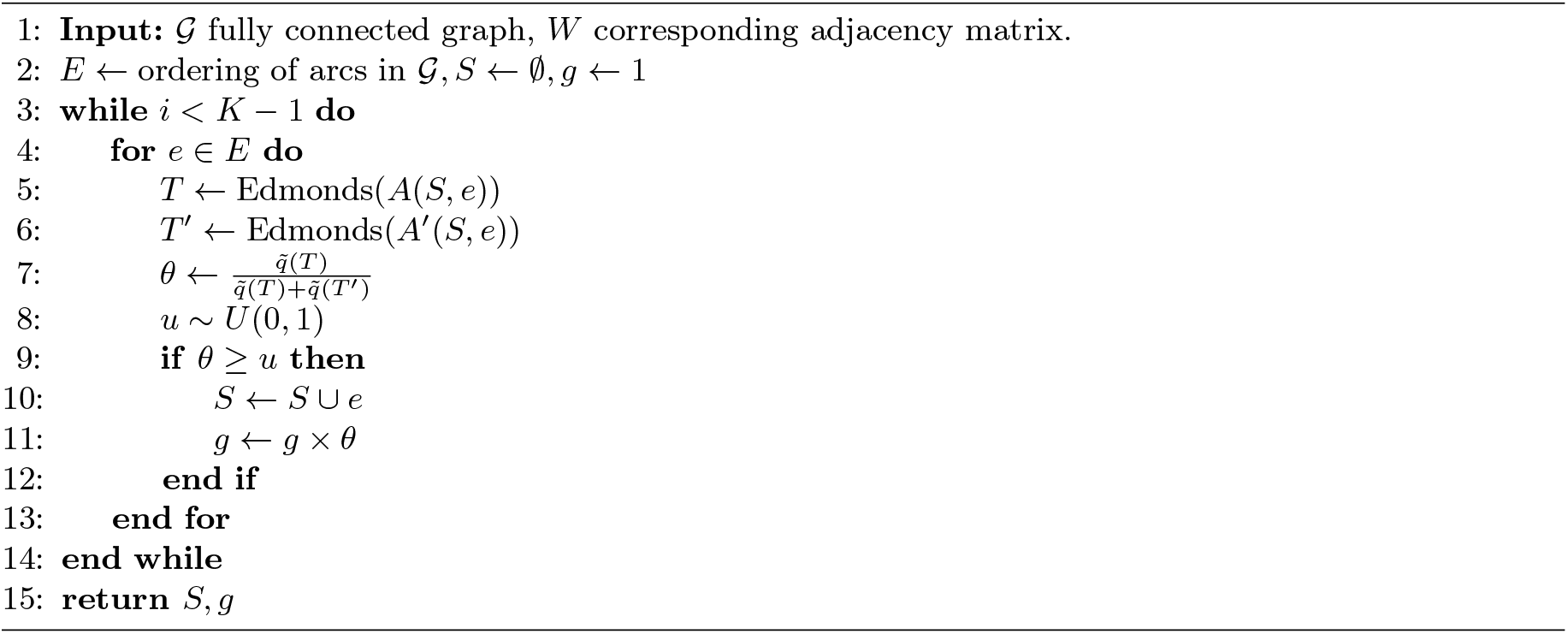

### S7 Split-merge

We run our split-merge algorithm every 10th iteration during optimization. First, we merge by checking the L1 distance between Viterbi paths of all clusters CNPs *q*(*C*^*i*^), *q*(*C*^*j*^). If the distance all candidate clusters *k* with *q*(*Z*_*n*_ = *k*) *>* 0.01 we group the cells with arg max_*j*_ *q*(*Z*_*n*_ = *j*) = *k* is smaller than a threshold, we merge *j* into *i* by re-initializing the parameters of *q*(*C*^*j*^) randomly, setting *q*(*Z*_*n*_ = *j*) = 0 and doubling *q*(*Z*_*n*_ = *i*). Then, if there exists a degenerate cluster, *l*, i.e. a cluster with total probability ∑*q*(*Z*_*n*_ = *l*) smaller than a threshold, we run the split algorithm: For all candidate clusters *k* with ∑*q*(*Z*_*n*_ = *k*)>0.01 we group the cells with arg max_*j*_ *q*(*Z*_*n*_ = *j*) = *k* into two groups, *k*_1_ and *k*_2_ by KMeans. Then, we construct temporary 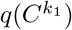 and 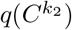 by applying the CAVI update using only the cells in *k*_1_ and *k*_2_ respectively, and then evaluate the ELBO using these new *q*(*C*)s. We select to split the candidate cluster *k* that results in the maximum ELBO into *l*.

## Footnotes

5 the time refers to the execution of VICTree on a CPU Intel Core i7-855U

6 ***π*** = (*π*_1_, …, *π*_|*Σ*|_) is actually a 1-D vector, which means there is no need to use *ι* mapping

