## Supplementary material for "VICTree - a Variational Inference method for Clonal Tree reconstruction"

### S1 Additional experiments

#### S1.1 Synthetic data simulation

#### S1.2 Robustness on high mutation rate

#### S1.3 Breast cancer sample

#### S1.4 Multiple Myeloma samples

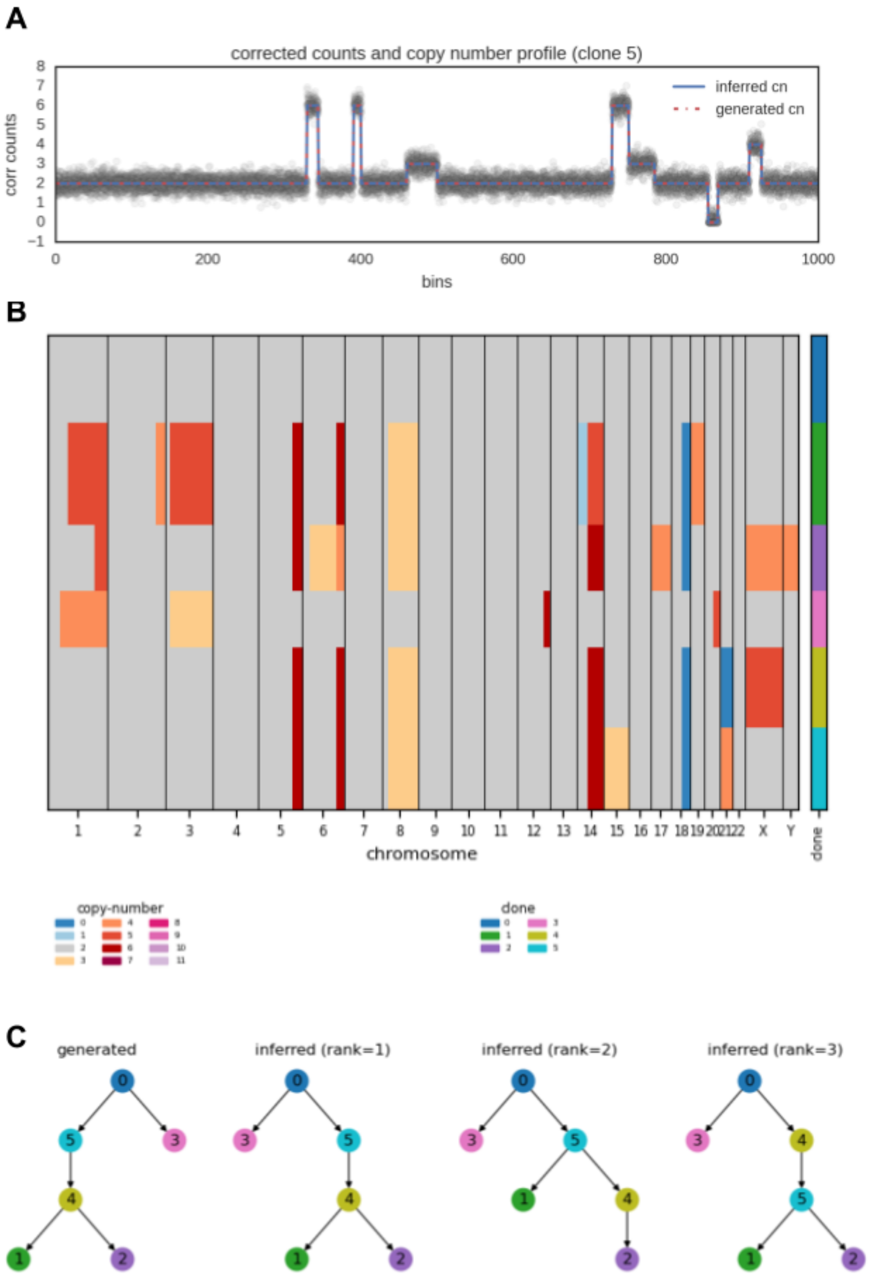


Fig. S1: Example of synthetic dataset with $K = 6$ clones. **A** Corrected counts of cells assigned to clone 5 with ground-truth and inferred copy number profile (CN-MAD $= 0.0$). **B** Clonal copy number profile with cells on the vertical axis, arranged by inferred clonal assignment (ARI $= 1.0$). **C** Generated (ground-truth) tree and set of inferred trees, ranked by importance in the final sample from $q(T)$ using LARS.

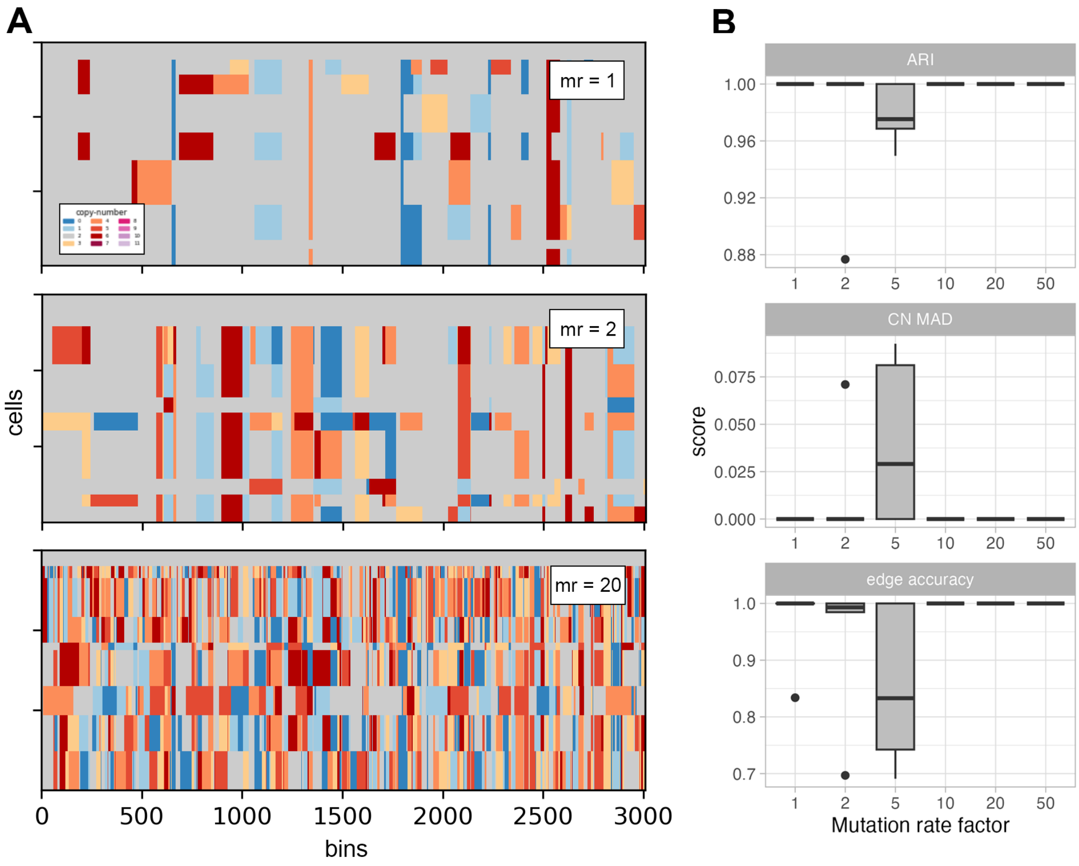


Fig. S2: Illustration on the robustness over increasing rate of copy number mutation. **A** Simulated copy number evolution of 9 clones with three different values for the mutation rate parameter: 1 (default), 2 and 20. **B** ARI (clustering), CN MAD (copy number calling) and edge accuracy scores against ground truth on 10 synthetic datasets for 6 different values of mutation rate parameter.

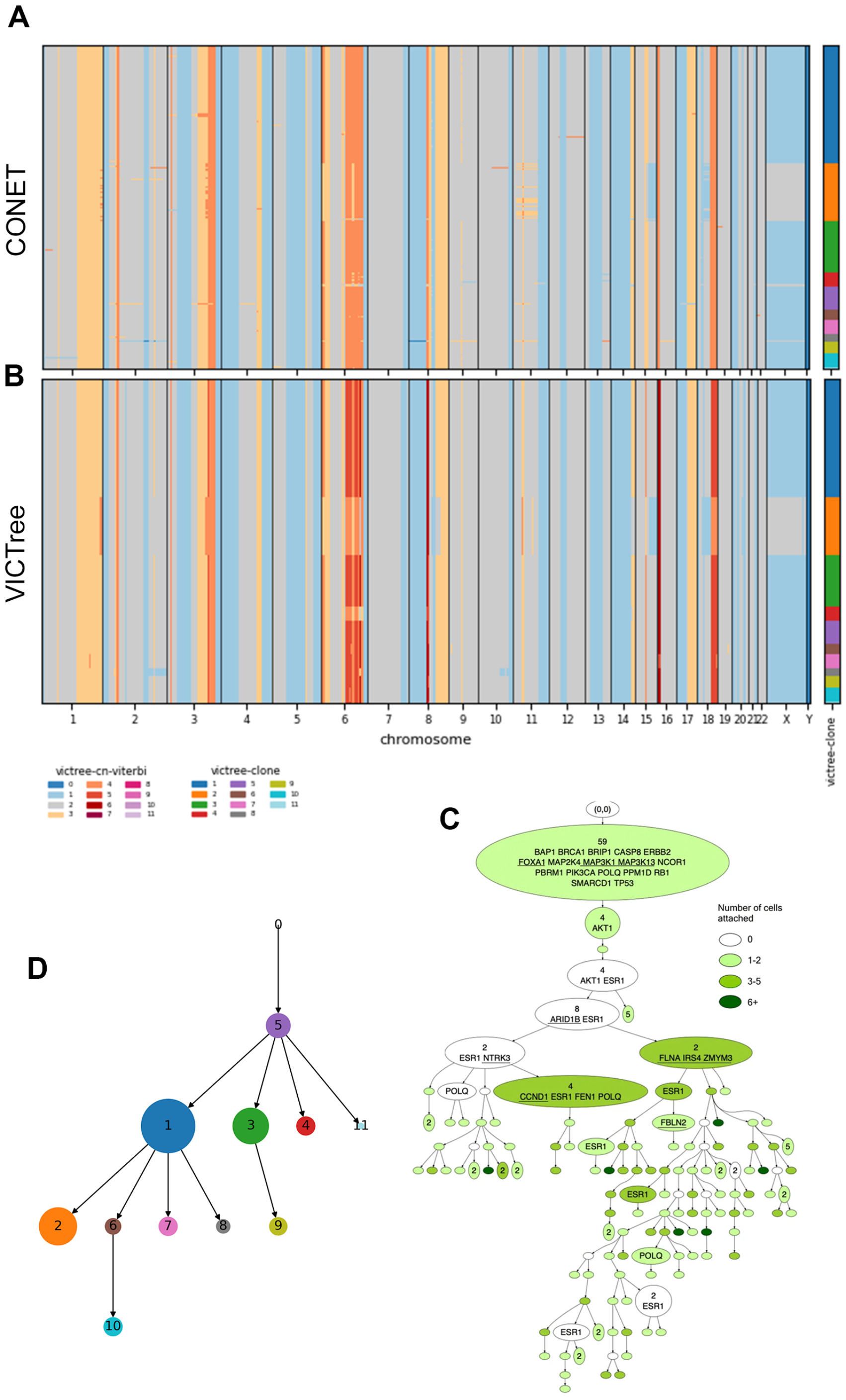


Fig. S3: Sample of xenograft breast cancer SA501X3F. Features 260 single cells for 18175 bins of width 150kb. **A** CONET copy-number calling (sorted and annotated by VICTree clustering). **B** VICTree clonal copy-number calling and clustering of cells annotated on the right. **C** MAP tree from running VICTree with $K = 14$, where each nodes has size proportionally to the clone size according to VICTree result, and color matching the legend on the plots above. **D** CONET tree as shown in [25].

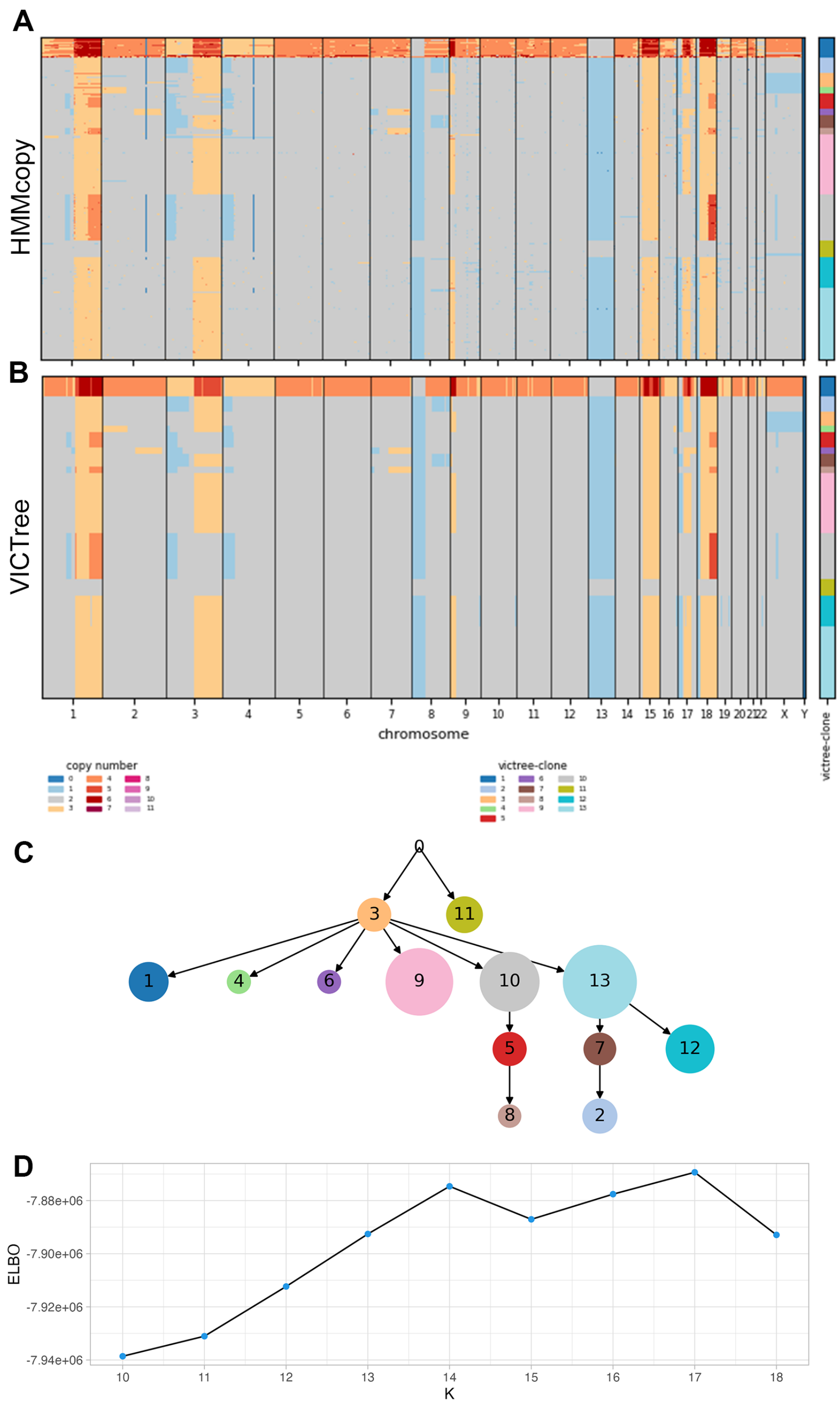


Fig. S4: Sample of multiple myeloma MM03. Features 2208 single cells for 5298 bins of width 500kb. **A** HMMcopy copy-number calling. **B** VICTree clonal copy-number calling and clustering of cells annotated on the right. **C** MAP tree from running VICTree with $K = 14$, where each nodes has size proportionally to the clone size according to VICTree result, and color matching the legend on the plots above. **D** Plot of the ELBO for runs with several values of $K$.

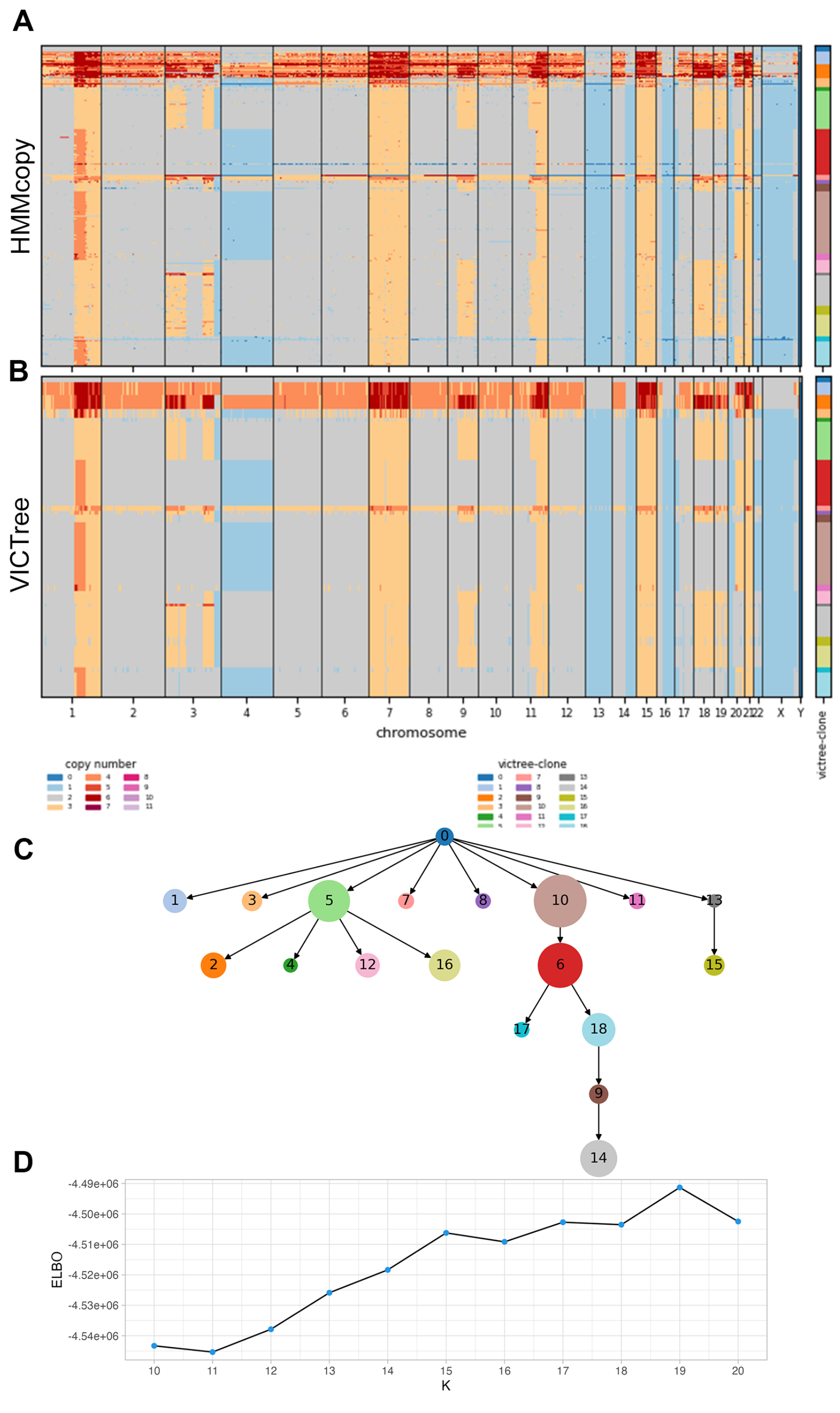


Fig. S5: Sample of multiple myeloma MM04. Features 1003 single cells for 5286 bins of width 500kb. **A** HMMcopy copy-number calling. **B** VICTree clonal copy-number calling and clustering of cells annotated on the right. **C** MAP tree from running VICTree with $K = 19$, where each nodes has size proportionally to the clone size according to VICTree result, and color matching the legend on the plots above. **D** Plot of the ELBO for runs with several values of $K$.

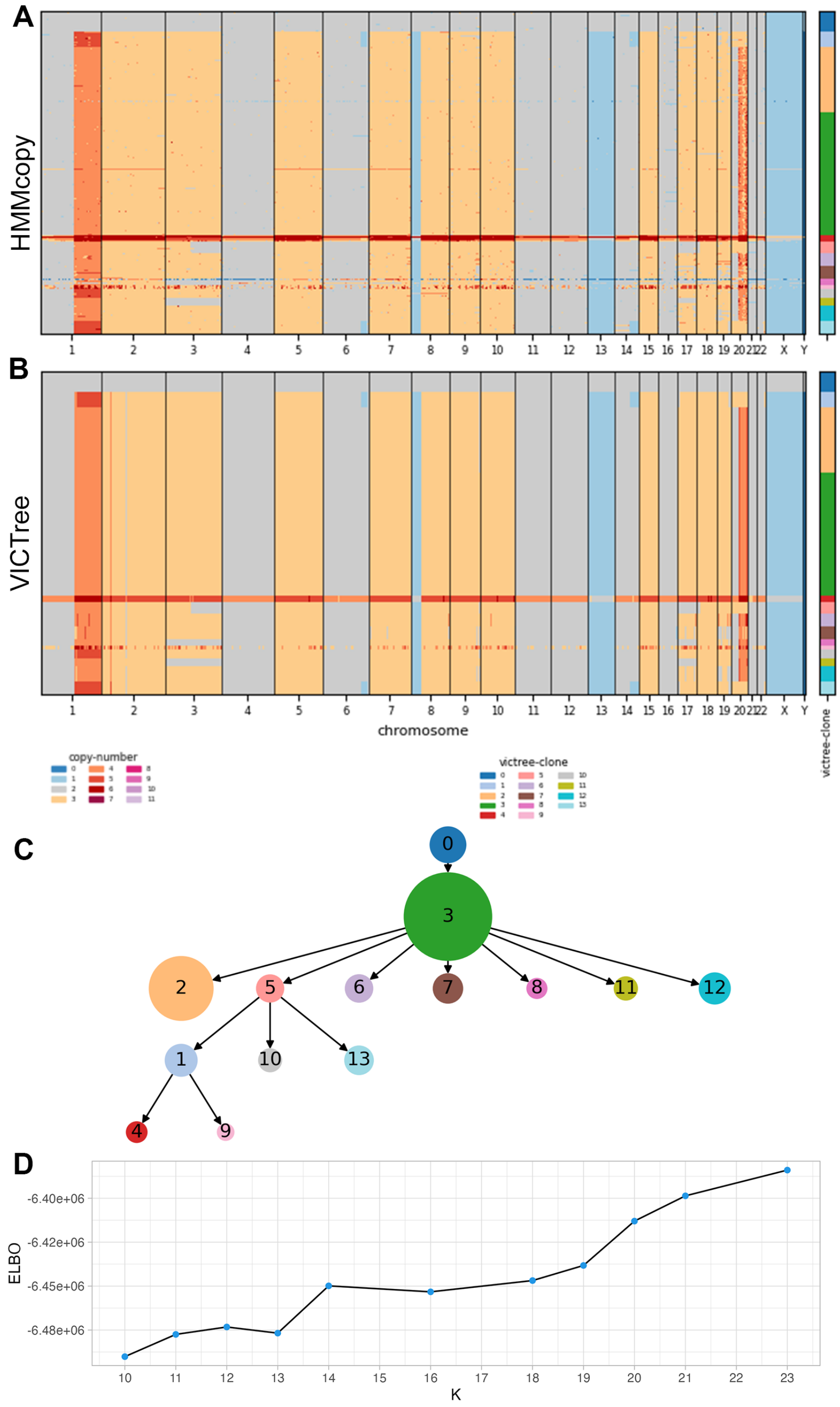


Fig. S6: Sample of multiple myeloma MM29. Features 1616 single cells for 5285 bins of width 500kb. **A** HMMcopy copy-number calling. **B** VICTree clonal copy-number calling and clustering of cells annotated on the right. **C** MAP tree from running VICTree with $K = 14$, where each nodes has size proportionally to the clone size according to VICTree result, and color matching the legend on the plots above. **D** Plot of the ELBO for runs with several values of $K$.

### S2 CopyTree in exponential form

In the following sections we prove that CopyTree model belongs to the exponential family of distributions. We first introduce the notation by showing how standard non-homogeneous HMMs can be written in exponential form and then we extend the proof for the tree-structured HMM which we call CopyTree. The derivations in this sections will also be useful for Section S3 and Section S4 since the notation is first introduced here.

#### S2.1 HMM in exponential form

Let $M$ be the length of a discrete Markov chain and $\Sigma$ be the finite discrete set of values for the Markov chain states (alphabet). Let $\boldsymbol{X}$ be the $|\Sigma| \times M$ matrix representing a sequence $\mathbf{C}$ where each column is the one-hot-encoding of a state, i.e.

$$X_{im} = \mathbb{1}(C_m = i)$$

Also, we introduce two mappings which are useful for parameters vectorization:

$$\begin{aligned} \iota(m, j) &= (m-1)|\Sigma| + j \\ \kappa(m, j, j') &= (m-1)|\Sigma|^2 + (j-1)|\Sigma| + j' , \end{aligned} \tag{S1}$$

with $j, j'$ taking values in $\Sigma$. Note that the vectors are linear and quadratic functions of the alphabet length.

Let us denote with $x_m$ the $m$-th column of $X$, which is the one-hot-encoding for the state at site $m$, and with $Y_m$ the vector of observations for site $m$. Then we can write the joint probability as

$$\begin{aligned} P(\boldsymbol{X}, \boldsymbol{Y}) &= P(Y_1|x_1)P(x_1) \prod_{m=2}^{M} P(Y_m|x_m)P(x_m|x_{m-1}) \\ &= P(Y_1|x_1) \prod_{m=2}^{M} P(Y_m|x_m) \prod_{i\in\Sigma} \pi_{\iota(1,i)}^{X_{i1}} \prod_{j\in\Sigma} \Phi_{\kappa(m,i,j)}^{X_{i,m-1}X_{jm}} \end{aligned} \tag{S2}$$

where $\boldsymbol{\pi}$ and $\boldsymbol{\Phi}$ are the initial and transition probability vectors respectively, with indices provided by the mapping just described [6]. For simplicity, we can define the pairwise state column vector $\tilde{x}$ which is indexed by $\kappa$ and has values given by

$$\tilde{x}_{\kappa(m,i,j)} \coloneqq X_{i,m-1}X_{j,m} = \mathbb{1}(C_{m-1} = i, C_m = j) .$$

We can then re-write part of (S2) as exponential functions of the parameters.

$$\begin{aligned} P(x_1) &= \prod_{i\in\Sigma} \pi_i^{X_{i1}} \\ &= \exp(\sum_{i\in\Sigma} X_{i1} \log \pi_i) \\ &= \exp(\log \boldsymbol{\pi}^\top x_1) \end{aligned} \tag{S3}$$

[6] $\boldsymbol{\pi} = (\pi_1, ..., \pi_{|\Sigma|})$ is actually a 1-D vector, which means there is no need to use $\iota$ mapping

$$\prod_{m=2}^{M} P(x_m|x_{m-1}) = \prod_{m=2}^{M} \prod_{i\in\Sigma} \prod_{j\in\Sigma} \Phi_{\kappa(m,i,j)}^{X_{i,m-1}X_{j,m}}$$
$$= \exp(\sum_{m=2}^{M} \sum_{i\in\Sigma} \sum_{j\in\Sigma} X_{i,m-1}X_{j,m} \log \Phi_{\kappa(m,i,j)})$$
$$= \exp(\log \boldsymbol{\Phi}^\top \tilde{x}) \tag{S4}$$

where $\log \boldsymbol{\pi} := (\log \pi_1, .., \log \pi_{|\Sigma|})$. Thus, all together in matrix notation:

$$P(\boldsymbol{X}) = \exp(\begin{bmatrix}\log \boldsymbol{\pi} \\ \log \boldsymbol{\Phi}\end{bmatrix}^\top \begin{bmatrix} x_1 \\ \tilde{x}\end{bmatrix}) \,. \tag{S5}$$

**Definition 1 (Exponential family form).**

*Let $X$ be a random vector whose distribution belongs to the exponential family and $\theta$ denote its parameters vector. Then the density function of $X$ is*

$$f_X(\mathbf{x}|\boldsymbol{\theta}) = \exp[\eta_X(\boldsymbol{\theta}) \cdot \mathbf{t}_X(\mathbf{x}) - A_X(\boldsymbol{\theta}) + B_X(\mathbf{x})], \tag{S6}$$

*where $\mathbf{t}_X(\mathbf{x})$ is the sufficient statistic vector, $\eta_X(\boldsymbol{\theta})$ is the natural parameter vector, $A_X(\boldsymbol{\theta})$ is the log partition and $B_X(\mathbf{x})$ is the log base measure.*

If we assume that the emission probability can be written in exponential form (e.g. product of Gaussians with mean defined by $\mathbf{X}$) and that it is conjugate to $P(\mathbf{X})$, we can write $P(\mathbf{Y}|\mathbf{X})$ in terms of $\mathbf{t}_X(\mathbf{X})$

$$P(\mathbf{Y}|\mathbf{X}) = \prod_{m=1}^{M} P(Y_m|x_m)$$
$$= \exp(\sum_{m=1}^{M} [\eta_{Y_m}(x_m)^\top \mathbf{t}_{Y_m}(Y_m) - A_{Y_m}(x_m) + B_{Y_m}(Y_m)])$$
$$= \exp(\eta_Y(\mathbf{X})^\top \mathbf{t}_Y(\mathbf{Y}) - A_Y(\mathbf{X}) + B_Y(\mathbf{Y}))$$
$$= \exp(\eta_{YX}(\mathbf{Y})^\top \mathbf{t}_X(\mathbf{X}) - A_{YX}(\mathbf{Y}) + B_{YX}(\mathbf{X}))\,. \tag{S7}$$

This reparametrization is also shown in [33] when presenting the Conjugate-Exponential models. It is worth noting that $P(\mathbf{Y}|\mathbf{X})$ can be thought as a contribution to the likelihood of $X$. Note that $\mathbf{t}_X$ is a concatenation of the flattened version of $\mathbf{X}$ and the $\tilde{x}$ vector previously introduced. The natural parameters vector $\eta_{YX}(\mathbf{Y})$ depends on the distribution posit on the observations. More specifically, $\eta_{YX}(\mathbf{Y})_{\iota(m,i)} = \log p(Y_m|C_m = i)$, function of $Y$ with fixed $C_m$ as it is multiplied by $X_{i,m}$, therefore "activated" or "deactivated".

Putting altogether, after reparametrizing as in (S7) and rearranging the terms at the exponent, we obtain the complete formulation

$$
\begin{aligned}
P(\mathbf{X},\mathbf{Y}) &= P(\mathbf{X})P(\mathbf{Y}|\mathbf{X}) \\
&= \exp\left(\begin{bmatrix}\log\boldsymbol{\pi}\\ \log\boldsymbol{\Phi}\end{bmatrix}^\top \begin{bmatrix}x_1\\ \tilde{x}\end{bmatrix} + \eta_X(\mathbf{Y})^\top \mathbf{t}_{YX}(\mathbf{X}) - A_{YX}(\mathbf{Y}) - B_{YX}(\mathbf{X})\right) \\
&= \exp\left(\begin{bmatrix}\log\boldsymbol{\pi}\\ \eta_{YX}(\mathbf{Y})\\ \log\boldsymbol{\Phi}\end{bmatrix}^\top \begin{bmatrix}x_1\\ \mathbf{t}_X(\mathbf{X})\\ \tilde{x}\end{bmatrix} - A_{YX}(\mathbf{Y}) - B_{YX}(\mathbf{X})\right). \qquad \text{(S8)}
\end{aligned}
$$

Since $\eta_{YX}(\mathbf{Y})$ needs to match the length of $\mathbf{t}_X(\mathbf{X})$ i.e. $M\times|\Sigma|+M\times|\Sigma|^2$, we set $\eta_{YX}(\mathbf{Y})_{\kappa(m,j,j')} = \mathbf{0}$. We further notice that $x_1$ is just the first column of $\boldsymbol{X}$, hence the first $|\Sigma|$ elements of the flattened vector $\mathbf{t}_X(\boldsymbol{X})$. This means that we can group $x_1$ by incorporating $\log\boldsymbol{\pi}$ with $\eta_{YX}(\boldsymbol{Y})$. We split the sufficient statistic vector into $t_1(\mathbf{X}), t_2(\mathbf{X})$ and the natural parameter vector into $\eta_1(\mathbf{Y},\boldsymbol{\theta},\boldsymbol{\pi}), \eta_2(\boldsymbol{\Phi})$

$$
P(\mathbf{X},\mathbf{Y}) = \exp(\begin{bmatrix}\eta_1(\mathbf{Y},\boldsymbol{\theta},\boldsymbol{\pi})\\ \eta_2(\boldsymbol{\Phi})\end{bmatrix}^\top \begin{bmatrix}t_1(\mathbf{X})\\ t_2(\mathbf{X})\end{bmatrix} - A_{YX}(\mathbf{Y}) - B_{YX}(\mathbf{X})) \qquad \text{(S9)}
$$

where $\boldsymbol{\theta}$ is the set of extra parameters required in the emission distribution defined on the observations $\mathbf{Y}$.

#### S2.2 CopyTree in exponential form

We will now refer to the variables using their biological meaning e.g. denoting by "copy-number profile" the latent states of the Markov chain, but the derivations are context independent and can be applied to any such model. The purpose of this section is to show how the conditional probability in each node of the Bayesian graph Figure S7 can be expressed as a function of the central node $v$'s sufficient statistic. These manipulations, besides allowing for Variational Message Passing [33], better explain the variational updates in section S3.

**Root node** Let's denote with $\mathbf{C}^r$ the copy number sequence of the root node and with $\mathbf{Y}^r$ the observations to it related. Since this does not depend on any other copy number sequence, we say that $(\mathbf{C}^r,\mathbf{Y}^r) \sim \text{HMM}(\boldsymbol{\phi},\boldsymbol{\psi})$ with $\boldsymbol{\phi} = \{\boldsymbol{\pi},\boldsymbol{\Phi}\}$ being the Markov chain parameters (initial state probability vector and transition matrix), and $\boldsymbol{\psi}$ being the set of parameters for the chosen emission distribution, whatever it may be.

Following the derivations from the previous section, the joint density function of $\mathbf{C}^r, \mathbf{Y}^r$ can be written in exponential form as

$$
f_{\text{HMM}}(\mathbf{C}^r,\mathbf{Y}^r|\boldsymbol{\phi},\boldsymbol{\psi}) = \exp\left(\eta_{\mathbf{C}^r}(\boldsymbol{\phi},\boldsymbol{\psi})\cdot \mathbf{t}_{\mathbf{C}^r}(\mathbf{C}^r) - A_{\mathbf{C}^r}(\boldsymbol{\phi},\boldsymbol{\psi}) + B_{\mathbf{C}^r}(\mathbf{C}^r)\right). \qquad \text{(S10)}
$$

For a more readable notation we simply specify the node to which the parameter is related, which implicitly sets the right variables upon which a certain function depends. E.g. $\eta(r) \coloneqq \eta_{\mathbf{C}^r}(\boldsymbol{\phi},\boldsymbol{\psi})$. Equation S10 can be therefore rewritten as

$$
f_{\text{HMM}}(\mathbf{C}^r,\mathbf{Y}^r|\boldsymbol{\phi},\boldsymbol{\psi}) = \exp\left(\eta(r)\cdot \mathbf{t}(r) - A(r) + B(r)\right). \qquad \text{(S11)}
$$

Both the sufficient statistic and the natural parameter vectors are easily represented as the concatenation of two vectors

$$\mathbf{t}(r) = \begin{bmatrix} t_1(r) \\ t_2(r) \end{bmatrix}, \eta(r) = \begin{bmatrix} \eta_1(r) \\ \eta_2(r) \end{bmatrix}. \tag{S12}$$

In subsection S2.1 we show that the vectors' elements are as follows

$$\begin{aligned} t_1(r)_{\iota(m,j)} &= \mathbb{1}(C^r_m = j) \\ t_2(r)_{\kappa(m,j,j')} &= \mathbb{1}(C^r_{m-1} = j, C^r_m = j') \\ \eta_1(r)_{\iota(1,j)} &= \log \pi_j + \log p_\psi(\mathbf{Y}_1 | C^r_1 = j) \\ \eta_1(r)_{\iota(m,j)} &= \log p_\psi(\mathbf{Y}_m | C^r_m = j) \\ \eta_2(r)_{\kappa(m,j,j')} &= \log \Phi_{\kappa(m,j,j')}. \end{aligned} \tag{S13}$$

**Tree HMM joint** We write the root node $r$ HMM probability as the standard HMM joint since it does not have a parent node.

$$p_{\phi\psi}(\mathbf{C}^r, \mathbf{Y}^r) = p(C^r_1)p(\mathbf{y}^r_1|C^r_1) \prod_{m=2}^{M} p(C^r_m|C^r_{m-1})p(\mathbf{y}^r_m|C^r_m), \tag{S14}$$

Then, for any internal node $u$ with parent $v$ we get the probability of the HMM of a child $v$ of $u$, given the parent as

$$\begin{aligned} &p_{\varepsilon\psi}(\mathbf{C}^v, \mathbf{Y}^v|\mathbf{C}^u) = \\ &= p(C^v_1|\mathbf{C}^u)p(\mathbf{y}^v_1|C^v_1, \mathbf{C}^u) \prod_{m=2}^{M} p(C^v_m|C^v_{m-1}, \mathbf{C}^u)p(\mathbf{y}^v_m|C^v_m, \mathbf{C}^u) \\ &= p(C^v_1|C^u_1)p(\mathbf{y}^v_1|C^v_1) \prod_{m=2}^{M} p(C^v_m|C^v_{m-1}, C^u_m, C^u_{m-1})p(\mathbf{y}^v_m|C^v_m) \end{aligned} \tag{S15}$$

where in step (S15) we assume that the transition probability of the child only depends on the two copy numbers of the parent at bin $m-1, m$.

The complete joint probability $p_{\phi\psi\varepsilon}(\mathbf{C}, \mathbf{Y})$ should be derived from root to leaves given the two quantities above in a chain product.

**Internal copy-number node** For an internal node of the tree, we introduce the CN coherence function, defined over each single copy number of the sequence. For any node $v$ and its parent $u$, the CN coherence function $h$ over the copy number values at a specific bin $m$ is defined as in (4).

For the first element of the sequence we simply redefine $h$ to be

$$\begin{aligned} h(j|i, \varepsilon_0) &= P(C^v_1 = j | C^u_1 = i) \\ &= \begin{cases} 1 - \varepsilon_0 & \text{if } i = j \\ \varepsilon_0/(|\Sigma| - 1) & \text{otherwise,} \end{cases} \end{aligned} \tag{S16}$$

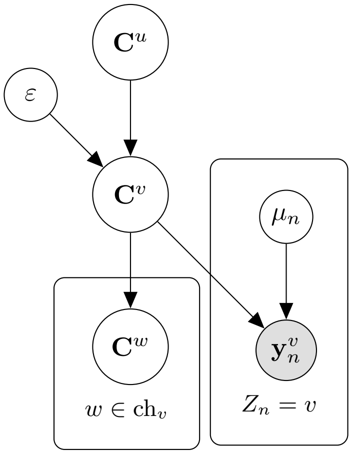


Fig. S7: Simple copy-tree Bayes graph representing the neighborhood of a node $v$ of the tree. For a generic node $v$, $\mathbf{C}^v$ is the latent copy number sequence, $\mathbf{y}_n^v$ is the sequence of observations for each cell $n$ in node $v$, $\varepsilon^v$ the co-mutation parameter and $\mu_n$ is the baseline (per-copy expression).

In the context of a variational message passing approach [33], it can be useful to draw a simple graph encoding the dependencies (see Figure S7).

The copy number sequences follow a Markov model. Let $u, v$ the parent-child pair of node, as usual, and let $\boldsymbol{X}^u, \boldsymbol{X}^v$ be the copy number sequences of the respective nodes in the one-hot encoding described above.

$$
\begin{aligned}
&\log f_{X^u}(\boldsymbol{X}^v|\boldsymbol{X}^u, \varepsilon) \\
&= \log P(x_1^v|x_1^u) + \sum_{m=2}^{M} \log P(x_m^v|x_{m-1}^v, x_m^u, x_{m-1}^u) \\
&= \sum_{i,j} X_{i,1}^u X_{j,1}^v \log h(j|i, \varepsilon) \\
&+ \sum_{m=2}^{M} \sum_{i,i'} \sum_{j,j'} X_{i,m-1}^u X_{j,m-1}^v X_{i',m}^u X_{j',m}^v \log h(j'|j, i', i, \varepsilon) \qquad \text{(S17)}
\end{aligned}
$$

In subsection S2.2 we have decomposed the sufficient statistic vector for HMMs in

$$\mathbf{t}_X(\boldsymbol{X}) = \begin{bmatrix} t_X^1(\boldsymbol{X}) \\ t_X^2(\boldsymbol{X}) \end{bmatrix}$$

with $t_X^1$ and $t_X^2$ being two column vectors indexed by $\iota$ and $\kappa$ mappings respectively. More specifically,

$$t^1_X(\boldsymbol{X})_{\iota(m,i)} = X_{i,m}$$
$$t^2_X(\boldsymbol{X})_{\kappa(m,i,i')} = X_{i',m}X_{i,m-1}.$$

With that in mind, we regroup (S17) to obtain the natural parameter vector $\eta_{X^v}(\boldsymbol{X}^u) = \left(\eta^1_{X^v}(\boldsymbol{X}^u), \eta^2_{X^v}(\boldsymbol{X}^u)\right)^\top$, which is then equal to

$$\begin{aligned} \eta^1_{X^v}(\boldsymbol{X}^u, \varepsilon)_{\iota(1,j)} &= \sum_i X^u_{i,1} \log h(j|i, \varepsilon_0) \\ \eta^2_{X^v}(\boldsymbol{X}^u, \varepsilon)_{\kappa(m,j,j')} &= \sum_{i,i'} X^u_{i',m}X^u_{i,m-1} \log h(j'|j, i', i, \varepsilon) \qquad 1 < m \leq M\,, \end{aligned} \tag{S18}$$

and the rest of the vector elements, namely $\eta^1_{X^v}(\boldsymbol{X}^u, \varepsilon)_{\iota(m,i)}$ for $1 < m \leq M$ and $\eta^2_{X^v}(\boldsymbol{X}^u, \varepsilon)_{\kappa(1,i,i')}$, are set to zero.

We can now write the conditional probability in (S17) as

$$\log f_{X^v}(\boldsymbol{X}^v|\boldsymbol{X}^u, \varepsilon) = \eta_{X^v}(\boldsymbol{X}^u, \varepsilon) \cdot \mathbf{t}_{X^v}(\boldsymbol{X}^v). \tag{S19}$$

It is then useful to rewrite this as a function of the sufficient statistic of the parent node's copy number, so that the rest can be used as a message going from child to parent (see [33]). Precisely, we want to find the vector $\eta_{X^vX^u}$ function of $\boldsymbol{X}^v$ such that

$$\log f_{X^v}(\boldsymbol{X}^v|\boldsymbol{X}^u, \varepsilon) = \eta_{X^vX^u}(\boldsymbol{X}^v, \varepsilon) \cdot \mathbf{t}_{X^u}(\boldsymbol{X}^u)$$

By symmetry of (S17), and since $\mathbf{t}_{X^u}$ has the same form of $\mathbf{t}_{X^v}$ we obtain $\eta_{X^vX^u}(\boldsymbol{X}^v, \varepsilon) = \left(\eta^1_{X^vX^u}(\boldsymbol{X}^v, \varepsilon), \eta^2_{X^vX^u}(\boldsymbol{X}^v, \varepsilon)\right)^\top$ such that

$$\begin{aligned} \eta^1_{X^vX^u}(\boldsymbol{X}^v, \varepsilon)_{\iota(1,i)} &= \sum_j X^v_{j,1} \log h(j|i, \varepsilon) \\ \eta^2_{X^vX^u}(\boldsymbol{X}^v, \varepsilon)_{\kappa(m,i,i')} &= \sum_{j,j'} X^v_{j',m}X^v_{j,m-1} \log h(j'|j, i', i, \varepsilon) \qquad 1 < m \leq M\,, \end{aligned} \tag{S20}$$

and the rest of the elements equal to zero. Note that even though the presented CN coherence function is symmetric, (S18) and (S20) might differ from the indices of $h$ when it is not symmetric on the pairs $(i, i')$ and $(j, j')$, therefore in general we have $\eta_{X^v} \neq \eta_{X^vX^u}$.

**Observations** Let $\boldsymbol{Y}^u$ be the set of observation at node $v$. For each bin $m$ we have a collection of reads, one for each cell assigned to the node and for each gene belonging to the bin. We denote this collection of reads with $\boldsymbol{Y}^v_m$.

To simplify the model, we assume that each of these reads range from 1 to $N$, independently of the cell/bin they belong to, and we assume that they are drawn from a Normal distribution, conditioned on the bin's copy number, with mean $\mu_n C^v_m$.

The conditional log-distribution is then

$$\begin{aligned}
&\log f_{Y^v}(\boldsymbol{Y}^v|\boldsymbol{X}^v, \boldsymbol{\mu}, \boldsymbol{\tau}) \\
&= \sum_{m=1}^{M} \log P(\boldsymbol{Y}_m^v|x_m^v, \boldsymbol{\mu}, \boldsymbol{\tau}) \\
&= \sum_{m=1}^{M} \sum_{n=1}^{N_v} \sum_{i \in \Sigma} X_{im} \log f_{\mathcal{N}}(y_{mn}^v; i \cdot \mu_n, \tau_n) \qquad \text{(S21)}
\end{aligned}$$

which can be expressed in terms of $\mathbf{t}_{X^v}(\boldsymbol{X}^v)$

$$\log f_{Y^v}(\boldsymbol{Y}^v|\boldsymbol{X}^v, \boldsymbol{\mu}, \boldsymbol{\tau}) = \eta_{Y^v X^v}(\boldsymbol{Y}^v, \boldsymbol{\mu}, \boldsymbol{\tau}) \cdot \mathbf{t}_{X^v}(\boldsymbol{X}^v) \,.$$

where $\mathbf{t}_{X^v}$ is the same as in (S2.2) and

$$\eta_{Y^v X^v}(\boldsymbol{Y}^v, \boldsymbol{\mu}, \boldsymbol{\tau}) = \begin{bmatrix} \eta^1_{Y^v X^v} \\ \mathbf{0} \end{bmatrix}$$

with $\mathbf{0}$ null vector of length matching $t^2_{X^v}$ (i.e. $M \times |\Sigma|^2$) and $\eta^1_{Y^v X^v}$ with same length as $t^1_{X^v}$, and elements such that

$$\eta^1_{Y^v X^v}(\boldsymbol{Y}^v)_{\iota(m,i)} = \sum_{n=1}^{N} \log f_{\mathcal{N}}(y_{mn}^v; i \cdot \mu_n, \tau_n)$$

### S3 Variational Updates

In this section we go through the derivations of the CAVI update for each variational distribution. All updates make use of the CAVI update equation (3).

#### S3.1 Copy numbers

**Copy numbers pre-requisite** We begin by observing that our model for a given $T$, when described as a directed graphical model on the variables

$$\{C_m^v : 1 \leq m \leq M, v \in V(T)\}$$

has a cyclic underlying undirected graph (see 2a).

However, it can be formulated as a directed graphical model on the variables

$$\{\mathbf{C}^v, v \in V(T)\}$$

where $p(\mathbf{C}^v|\mathbf{C}^{\rho_T(v)})$ is a conditional Markov model such that the underlying directed graphical model is a tree. In order to draw this conclusion we consider the conditional joint probability of observations and copy numbers

$$p(\mathbf{Y}, \mathbf{C}|\mathbf{Z}, T, \boldsymbol{\Psi}) = p(\mathbf{Y}|\mathbf{C}, \mathbf{Z}, \Psi)p(\mathbf{C}|T, \boldsymbol{\Psi})$$

and show how to express the above conditional joint as a product of conditional Markov models:

$$\prod_{uv \in A(T)} M(\mathbf{C}^v|\mathbf{C}^u)\,,$$

where the conditional Markov models depend on $\mathbf{Z}, \boldsymbol{\Psi}, \mathbf{Y}$. Notice that

$$\log p(\mathbf{Y}|\mathbf{C}, \mathbf{Z}, \boldsymbol{\Psi}) = \sum_{v \in V(T)} \sum_n \mathbb{1}(Z_n = v) \sum_m \log p(y_{mn}|C_m^v, \mu_n, \tau_n)\,,$$

and that

$$\log p(\mathbf{C}|T, \boldsymbol{\Psi}) = \sum_{uv \in A(T)} \log p(\mathbf{C}^v|\mathbf{C}^u, \varepsilon_{uv}).$$

This implies

$$\begin{aligned}
&\log p(\mathbf{Y}, \mathbf{C}|\mathbf{Z}, T, \boldsymbol{\Psi}) \\
&= \sum_n \mathbb{1}(Z_n = r) \sum_m \log p(y_{mn}|C_m^r, \mu_n, \tau_n) \\
&\quad + \sum_{uv \in A(T)} \Big[\log p(\mathbf{C}^v|\mathbf{C}^u, \varepsilon_{uv}) + \sum_n [\mathbb{1}(Z_n = v) \sum_m \log p(y_{mn}|C_m^v, \mu_n, \tau_n)]\Big]
\end{aligned}$$

Hence, it is sufficient to find a conditional Markov model $M(\mathbf{C}^v|\mathbf{C}^u)$ for which the product of sufficient statistics and natural parameter is the outermost summand of the expression above, that is:

$$
\begin{aligned}
&\log M(\mathbf{C}^v|\mathbf{C}^u) = \\
&= \log p(\mathbf{C}^v|\mathbf{C}^u, \varepsilon_{uv}) + \sum_n \mathbb{1}(Z_n = v) \sum_m \log p(y_{mn}|C^v_m, \mu_n, \tau_n) \\
&= \log h_{\varepsilon_0}(C^v_1|C^u_1) + \sum_{m=2}^{M} \log h_{\varepsilon_{uv}}(C^v_m|C^v_{m-1}, C^u_m, C^u_{m-1}) \\
&\quad + \sum_n [\mathbb{1}(Z_n = v) \sum_m \log p(y_{mn}|C^v_m, \mu_n, \tau_n)]\,. 
\end{aligned} \tag{S22}
$$

The following shows how to express the Markov model for $M(\mathbf{C}^v|\mathbf{C}^u)$ in its exponential family form, that is, using the natural parameter $\boldsymbol{\eta}_T(v)$ and the sufficient statistics $\mathbf{t}_T(\mathbf{C}^v)$. Keep in mind that the sufficient statistics is a function of the copy number sequence, while the natural parameter depends on other parameters e.g. $\boldsymbol{\Psi}$. Let the parent node of $v$ given the tree $T$ simply be $\rho = \rho_T(v)$, and the quantities $\boldsymbol{\eta}_T(v), \mathbf{t}_T(\mathbf{C}^v)$ be just $\boldsymbol{\eta}(v), \mathbf{t}(\mathbf{C}^v)$, then the decomposition of the latters can be expressed in the following way.
Referring to the mappings in (S1), similarly to the notation introduced in section S2, the sufficient statistics vector $\mathbf{t}(\mathbf{C}^v)$ is equal to

$$
\begin{aligned}
\mathbf{t}(\mathbf{C}^v) &= (\mathbf{t}_1(\mathbf{C}^v), \mathbf{t}_2(\mathbf{C}^v))^T \\
t_1(\mathbf{C}^v)_{\iota(m,i)} &= \mathbb{1}(C^v_m = i) \\
t_2(\mathbf{C}^v)_{\kappa(m,i,i')} &= \mathbb{1}(C^v_{m-1} = i, C^v_m = i')\,.
\end{aligned}
$$

Natural parameters, $\boldsymbol{\eta}(v)$, and associated parameters, $\boldsymbol{\eta}'(v)$:

$$
\begin{aligned}
\boldsymbol{\eta}(v) &= (\boldsymbol{\eta}_1(v), \boldsymbol{\eta}_2(v)) \\
\boldsymbol{\eta}_1(v) &= \boldsymbol{\eta}'_1(v) + \boldsymbol{\eta}''_1(v) \\
\eta'_1(v)_{\iota(1,i)} &= \sum_j \mathbb{1}(C^\rho_1 = j) \log h_{\varepsilon_0}(i|j) \\
\eta'_1(v)_{\iota(m,i)} &= 0, \text{ for all } m \neq 1 \\
\eta''_1(v)_{\iota(m,i)} &= \sum_n \mathbb{1}(Z_n = v) \log p(y_{mn}|C^v_m = i, \Psi) \\
\boldsymbol{\eta}'(v) &= (\boldsymbol{\eta}'_1(v), \boldsymbol{\eta}_2(v)) \\
\eta_2(v)_{\kappa(m,i,i')} &= \sum_{j,j'} \mathbb{1}(C^\rho_{m-1} = j, C^\rho_m = j') \log h_{\varepsilon_{\rho v}}(i'|i, j', j)\,,
\end{aligned}
$$

where both $\boldsymbol{\eta}(v)$ and $\boldsymbol{\eta}'(v)$ are vectors of dimension $|\Sigma| \times M + |\Sigma|^2 \times M$, and match the dimension of $\mathbf{t}(\mathbf{C}^v)$.

Notice that, as desired, the inner product of such natural parameters with sufficient statistic vectors, matches the Markov model in (S22):

$$
\begin{aligned}
&\boldsymbol{\eta}(v)\mathbf{t}(\mathbf{C}^v) \\
&= \sum_i \mathbb{1}(C_1^v = i) \sum_j \mathbb{1}(C_1^\rho = j) \log h_{\varepsilon_0}(i|j) \\
&\quad + \sum_m \sum_i \mathbb{1}(C_m^v = i) \sum_n \mathbb{1}(Z_n = v) \log p(y_{mn}|C_m^v = i, \Psi) \\
&\quad + \sum_m \sum_{i,i'} \mathbb{1}(C_{m-1}^v = i, C_m^v = i') \sum_{j,j'} \mathbb{1}(C_{m-1}^\rho = j, C_m^\rho = j') \log h_{\varepsilon_{\rho v}}(i'|i, j', j)\,.
\end{aligned}
\tag{S23}
$$

Moreover,

$$
\begin{aligned}
&\boldsymbol{\eta}'(v)\mathbf{t}(\mathbf{C}^v) = \\
&= \sum_i \mathbb{1}(C_1^v = i) \sum_j \mathbb{1}(C_1^\rho = j) \log h_{\varepsilon_0}(i|j) \\
&\quad + \sum_{m=2}^{M} \sum_{i,i'} \mathbb{1}(C_{m-1}^v = i, C_m^v = i') \sum_{j,j'} \mathbb{1}(C_{m-1}^\rho = j, C_m^\rho = j') \log h_{\varepsilon_{\rho v}}(i'|i, j', j) \\
&= \log h_{\varepsilon_{\rho v}}(C_1^v|C_1^\rho) + \sum_{m=2}^{M} \log h_{\varepsilon_{\rho v}}(C_m^v|C_{m-1}^v, C_m^\rho, C_{m-1}^\rho)\,.
\end{aligned}
\tag{S24}
$$

This should clarify the purpose of the natural and associated parameters: while the latter embodies the attributes of the Markov model over the copy number sequence, the former also links the sequence to the observations.

**Copy numbers update** We begin by noticing that $\mathbb{E}_{\mathbf{CZ}\varepsilon}\left[\log p(\mathbf{C}^r)\right]$ is uniquely determined by the prior that has probability 1 for the reference genome which typically has copy number 2 everywhere. For the the remaining $K-1$ vertices we provide the details for the standard CAVI updates. Let us first consider a fixed tree $T$ and any node $v \neq r$.

$$
\begin{aligned}
\log q^*(\mathbf{C}^v) &= \mathbb{E}_{-C^v}\left[\log p(\mathbf{Y}, \mathbf{C}, \boldsymbol{\Psi}, \mathbf{Z}|T)\right] \\
&\stackrel{+}{=} \mathbb{E}_{C^{-v}Z\Psi}\left[\log p(\mathbf{Y}|\mathbf{C}, \mathbf{Z}, \boldsymbol{\mu}, \boldsymbol{\tau}) p(\mathbf{C}|\boldsymbol{\Psi}, \mathbf{Z}, T)\right] \\
&\stackrel{+}{=} \mathbb{E}_{C^{-v}Z\Psi}\left[\log p(\mathbf{Y}|\mathbf{C}, \mathbf{Z}, \boldsymbol{\mu}, \boldsymbol{\tau})\right] + \mathbb{E}_{C^{-v}\Psi}\left[\log p(\mathbf{C}^r) + \sum_{uv \in A(T)} \log p(\mathbf{C}^v|\mathbf{C}^u)\right] \\
&\stackrel{+}{=} \mathbb{E}_{Z\Psi}\left[\sum_{n:Z_n=v} \log p(\mathbf{y}_n|\mathbf{C}^v, \mu_n, \tau_n)\right] + \sum_{uv \in A(T)} \mathbb{E}_{C^{-v},Z,\varepsilon}\left[\log p(\mathbf{C}^v|\mathbf{C}^u)\right] \\
&\stackrel{+}{=} \mathbb{E}_{Z\Psi}\left[\sum_{n:Z_n=v} \log p(\mathbf{y}_n|\mathbf{C}^v, \mu_n, \tau_n)\right] + \mathbb{E}_{C^{\rho_T(v)},\varepsilon}\left[\log p(\mathbf{C}^v|\mathbf{C}^{\rho_T(v)})\right] \\
&\quad + \sum_{w \in ch(v,T)} \mathbb{E}_{C^w\varepsilon}\left[\log p(\mathbf{C}^w|\mathbf{C}^v)\right] .
\end{aligned}
$$

Where $\rho_T(v)$ is the parent and ch(v, T) is the set of children of vertex $v$ in tree $T$. Moreover, as shown above,

$$
\begin{aligned}
&\mathbb{E}_{Z\Psi}\left[\sum_{n:Z_n=v} \log p(\mathbf{y}_n|\mathbf{C}^v, \mu_n, \tau_n)\right] + \mathbb{E}_{C^{\rho_T(v)},\varepsilon}\left[\log p(\mathbf{C}^v|\mathbf{C}^{\rho_T(v)})\right] = \\
&= \mathbb{E}_{C^{\rho_T(v)}Z\Psi}\left[M(\mathbf{C}^v|\mathbf{C}^{\rho_T(v)})\right] \\
&= \mathbb{E}_{C^{\rho_T(v)}Z\Psi}\left[\boldsymbol{\eta}_T(v)\right]\mathbf{t}(\mathbf{C}^v)
\end{aligned}
$$

and, which requires some manipulations,

$$
\begin{aligned}
&\mathbb{E}_{C^w\varepsilon}\left[\log p(\mathbf{C}^w|\mathbf{C}^v)\right] \\
&= \mathbb{E}_{C^w\varepsilon}\left[\boldsymbol{\eta}_T(\mathbf{C}^w)\mathbf{t}(\mathbf{C}^w)\right] \\
&\stackrel{+}{=} \mathbb{E}_{C^w\varepsilon}\left[\boldsymbol{\eta}'_T(\mathbf{C}^w)\mathbf{t}(\mathbf{C}^w)\right] \\
&= \mathbb{E}_{C^w\varepsilon}\Big[\sum_i \mathbb{1}(C_1^w = i) \sum_j \mathbb{1}(C_1^v = j) \log h_{\varepsilon_{vw}}(i|j) \\
&\quad + \sum_{m=2}^{M} \sum_{i,i'} \mathbb{1}(C_{m-1}^w = i, C_m^w = i') \sum_{j,j'} \mathbb{1}(C_{m-1}^v = j, C_m^v = j') \log h_{\varepsilon_{vw}}(i'|i, j', j)\Big] \\
&= \mathbb{E}_{C^w\varepsilon}\left[\boldsymbol{\alpha}(w)\right]\mathbf{t}(\mathbf{C}^v) ,
\end{aligned}
\tag{S25}
$$

where

$$
\alpha(w)_{\iota(m,i)} = \sum_j \mathbb{1}(C_m^w = j) \log h_{\varepsilon_{vw}}(j|i)
$$

and

$$
\alpha(w)_{\kappa(m,i,i')} = \sum_{j,j'} \mathbb{1}(C_{m-1}^w = j, C_m^w = j') \log h_{\varepsilon_{vw}}(j'|j, i', i).
$$

Note that the above two expressions have $j$ and $i$ inverted in $h(\cdot)$ because we express the indexing with $i$, but the copy numbers indexed by $j$ (which belong to node $w$), are named after $i$ in (S25).

Consequently,

$$\begin{aligned}
&\log q^*(\mathbf{C}^v) \qquad \text{(S26)}\\
&= \left( \mathbb{E}_{C^{-v}Z\Psi}\left[\boldsymbol{\eta}_T(v)\right] + \sum_{w\in\gamma_T(v)} \mathbb{E}_{C^w\varepsilon}[\boldsymbol{\alpha}(w)] \right) \mathbf{t}(\mathbf{C}^v)\\
&= \mathbb{E}_{C^{-v}Z\Psi}\left[\boldsymbol{\beta}_T(v)\right] \mathbf{t}(\mathbf{C}^v)
\end{aligned}$$

where $\boldsymbol{\beta}_T(v)$ can be be expressed as the concatenation of two vectors, i.e. $\beta_T(v) = (\beta_{T,1}, \beta_{T,2})$, where $\beta_{T,1}(\mathbf{C}^v)$ and $\beta_{T,2}(\mathbf{C}^v)$ are defined as follows

$$\begin{aligned}
\beta_{T,1_{\iota(m,i)}} = \mathbb{1}(m=1)\Big[ &\sum_j \mathbb{1}(C_1^\rho = j) \log h_{\varepsilon_{\rho v}}(i|j)\\
&+ \sum_{w\in\gamma_T(v)} \sum_j \mathbb{1}(C_1^w = j) \log h_{\varepsilon_{vw}}(j|i) \Big]\\
&+ \sum_n \mathbb{1}(Z_n = v) \log p(y_{mn}|C_m^v = i)\,,
\end{aligned}$$

and

$$\begin{aligned}
\beta_{T,2_{\kappa(m,i,i')}} = &\sum_{j,j'} \mathbb{1}(C_{m-1}^\rho = j, C_m^\rho = j') \log h_{\varepsilon_{\rho v}}(i'|i,j',j)\\
&+ \sum_{w\in\gamma_T(v)} \sum_{j,j'} \mathbb{1}(C_{m-1}^w = j, C_m^w = j') \log h_{\varepsilon_{vw}}(j'|j,i',i).
\end{aligned}$$

Finally, when considering the VI-distribution over trees $q(T)$, rather than a fixed tree, we obtain the final update

$$\begin{aligned}
&\log q^*(\mathbf{C}^v)\\
&= \mathbb{E}_{C^{-v}ZT\Psi}\left[ \log p(\mathbf{Y}|\mathbf{C}^v, \mathbf{Z}) + \log p(\mathbf{C}^v|\mathbf{C}^{\rho_T(v)}) + \sum_{w\in\gamma_T(v)} \log p(\mathbf{C}^w|\mathbf{C}^v) \right]\\
&= \mathbb{E}_T\left[ \mathbb{E}_{C^{-v}Z\Psi}\left[\boldsymbol{\eta}_T(v)\right] + \sum_{w\in\gamma_T(v)} \mathbb{E}_{C^w\varepsilon}\left[\boldsymbol{\alpha}(w)\right] \right] \mathbf{t}(\mathbf{C}^v)\,. \qquad \text{(S27)}
\end{aligned}$$

Therefore, we update the natural parameter as

$$\begin{aligned}
\boldsymbol{\eta}_T^*(v) &= \mathbb{E}_T\left[ \mathbb{E}_{C^\rho Z\Psi}\left[\boldsymbol{\eta}_T(v)\right] + \sum_{w\in\gamma_T(v)} \mathbb{E}_{C^w\varepsilon}\left[\boldsymbol{\alpha}(w)\right] \right]\\
&= \mathbb{E}_T\left[\mathbb{E}_{C\Psi Z}\left[\boldsymbol{\beta}_T(v)\right]\right]\,,
\end{aligned}$$

which can be decomposed, element by element, in

$$\begin{aligned}
\mathbb{E}_{C^\rho Z \Psi}\left[\eta_1(v)_{\iota(1,i)}\right] &= \sum_j q(C_1^\rho = j) \log h_{\varepsilon_0}(i|j) \\
&\quad + \sum_n q(Z_n = v) \mathbb{E}_{\mu_n \tau_n}\left[\log p(y_{mn}|C_m^v = i)\right] \\
\mathbb{E}\left[\eta_1(v)_{\iota(m,i)}\right] &= \sum_n q(Z_n = v) \mathbb{E}_{\mu_n \tau_n}\left[\log p(y_{mn}|C_m^v = i)\right] \\
\mathbb{E}_{C^\rho \Psi}\left[\eta_2(v)_{\kappa(m,i,i')}\right] &= \sum_{j,j'} q(C_{m-1}^\rho = j, C_m^\rho = j') \mathbb{E}_\varepsilon\left[\log h_{\varepsilon_{\rho v}}(i'|i,j',j)\right] \\
\mathbb{E}_{C^w}\left[\alpha_1(w)_{\iota(m,i)}\right] &= \sum_j q(C_m^w = j) \log h_{\varepsilon_0}(j|i) \\
\mathbb{E}_{C^w \varepsilon}\left[\alpha_2(w)_{\kappa(m,i,i')}\right] &= \sum_{j,j'} q(C_{m-1}^w = j, C_m^w = j') \mathbb{E}_\varepsilon\left[\log h_{\varepsilon_{vw}}(j'|j,i',i)\right] ,
\end{aligned} \tag{S28}$$

where

$$\begin{aligned}
&\mathbb{E}_{\mu_n \tau_n}\left[\log p(y_{mn}|C_m^v = i)\right] \\
&= \mathbb{E}_{\mu_n \tau_n}\left[\log\left(\sqrt{\frac{\tau_n}{2\pi}} \exp\left(-\frac{\tau_n}{2}(y_{mn} - i\mu_n)^2\right)\right)\right] \\
&\stackrel{+}{=} \mathbb{E}_{\mu_n \tau_n}\left[\frac{1}{2}\log \tau_n - \frac{\tau_n}{2}\left(y_{mn} - i\mu_n\right)^2\right] \\
&= \frac{1}{2}\mathbb{E}_{\tau_n}\left[\log \tau_n\right] - \frac{y_{mn}^2}{2}\mathbb{E}_{\tau_n}\left[\tau_n\right] + i y_{mn} \mathbb{E}_{\mu_n \tau_n}\left[\mu_n \tau_n\right] - \frac{i^2}{2}\mathbb{E}_{\mu_n \tau_n}\left[\mu_n^2 \tau_n\right] .
\end{aligned} \tag{S29}$$

Given that the variational distribution over $\mu_n, \tau_n$ is Normal-Gamma with parameters

$$\mu_n, \tau_n \stackrel{q}{\sim} \text{NormalGamma}(\tilde{\nu}_n, \tilde{\lambda}_n, \tilde{\alpha}_n, \tilde{\beta}_n) , \tag{S30}$$

the expectations in the previous formula are equal to:

$$\begin{aligned}
\mathbb{E}_{\tau_n}\left[\log \tau_n\right] &= \psi(\tilde{\alpha}_n) - \log \tilde{\beta}_n \\
\mathbb{E}_{\tau_n}\left[\tau_n\right] &= \frac{\tilde{\alpha}_n}{\tilde{\beta}_n} \\
\mathbb{E}_{\mu_n \tau_n}\left[\mu_n \tau_n\right] &= \tilde{\nu}_n \frac{\tilde{\alpha}_n}{\tilde{\beta}_n} \\
\mathbb{E}_{\mu_n \tau_n}\left[\mu_n^2 \tau_n\right] &= \frac{1}{\tilde{\lambda}_n} + \tilde{\nu}_n^2 \frac{\tilde{\alpha}_n}{\tilde{\beta}_n}
\end{aligned} \tag{S31}$$

#### S3.2 Tree topology

We begin by using the mean-field VI standard update:

$$\begin{aligned}
&\log q^*(T) &&\text{(S32)}\\
&= \mathbb{E}_{CZ\Psi}\left[\log p(\mathbf{Y}, \mathbf{C}, \boldsymbol{\Psi}, \mathbf{Z}, T)\right]\\
&\stackrel{+}{=} \mathbb{E}_{CZ\Psi}\left[\log p(\mathbf{Y}, \mathbf{C}|\boldsymbol{\Psi}, \mathbf{Z}, T)\right] + \log p(T)\\
&\stackrel{+}{=} \mathbb{E}_{CZ\Psi}\left[\log p(\mathbf{Y}|\mathbf{C}, \mathbf{Z}) + \log p(\mathbf{C}|\boldsymbol{\Psi}, \mathbf{Z}, T)\right] + \log p(T)\\
&\stackrel{+}{=} \mathbb{E}_{C\Psi}\left[\log p(\mathbf{Y}^r, \mathbf{C}^r|\mathbf{Z}) + \sum_{uv\in A(T)} \log M(\mathbf{C}^v|\mathbf{C}^u)\right] + \log p(T)\\
&= \mathbb{E}_{C^r Z\Psi}\left[\log p(\mathbf{Y}^r, \mathbf{C}^r|\mathbf{Z})\right] + \sum_{uv\in A(T)} \mathbb{E}_{C^v C^u Z\varepsilon}\left[\boldsymbol{\eta}_T(v)\mathbf{t}(\mathbf{C}^v)\right] + \log p(T)\\
&= \mathbb{E}_{C^r}\left[\log p(\mathbf{C}^r)\right] + \sum_{uv\in A(T)} \mathbb{E}_{C^v C^u \varepsilon}\left[\log p(\mathbf{C}^v|\mathbf{C}^u)\right] + \log p(T)
\end{aligned}$$

The expression in (S32) contains three terms that we further examine below.

First, $\mathbb{E}_{C^r Z}\left[\log p(\mathbf{C}^r|\mathbf{Z})\right]$, as stated above, is uniquely determined by the prior that has probability 1 for the reference healthy genome, which typically has copy number 2 everywhere. Furthermore, $\log p(T)$ is constant over the support of $T$ as we assign a uniform prior.

What varies over $T$ is the summand of the remaining term, $\mathbb{E}_{C^u C^v \varepsilon}\left[\log p(\mathbf{C}^v|\mathbf{C}^u)\right]$, for which we use the notation $w(uv)$ as it represents the unnormalized weight of each arc in the variational distribution. It can be further simplified as:

$$\begin{aligned}
&w(uv) = \mathbb{E}_{C^u C^v \varepsilon}\left[\log p(\mathbf{C}^v|\mathbf{C}^u)\right] &&\text{(S33)}\\
&= \mathbb{E}_{C^u C^v \varepsilon}\left[\log \prod_m \prod_{i,i',j,j'} h_{\varepsilon_{uv}}(i'|i,j',j)^{\mathbb{1}(C^v_{m-1}=i, C^v_m=i', C^u_{m-1}=j, C^u_m=j')}\right] &&\text{(S34)}\\
&= \sum_m \sum_{i,i'} \mathbb{E}_{C^v}\left[\mathbb{1}(C^v_{m-1}=i, C^v_m=i')\right] \sum_{j,j'} \mathbb{E}_{C^u}\left[\mathbb{1}(C^u_{m-1}=j, C^u_m=j')\right]\\
&\quad \times \mathbb{E}_{\varepsilon}\left[\log h_{\varepsilon_{uv}}(i'|i,j',j)\right] &&\text{(S35)}\\
&= \sum_m \sum_{i,i'} q(C^v_{m-1}=i, C^v_m=i') \sum_{j,j'} q(C^u_{m-1}=j, C^u_m=j')\mathbb{E}_{\varepsilon}\left[\log h_{\varepsilon_{uv}}(i'|i,j',j)\right], &&\text{(S36)}
\end{aligned}$$

where $q(C^u_{m-1}, C^u_m)$ can be computed iteratively for all $m$ with a single forward pass, which is all performed in $O(KMA)$.

$$q(C^u_m, C^u_{m-1}) = q(C^u_m|C^u_{m-1}=i)q(C^u_{m-1}=i) \qquad \text{(S37)}$$

$$q(C^u_m) = \sum_{i\in\Sigma} q(C^u_{m-1}, C^u_{m-2}=i), \qquad \text{(S38)}$$

and the expectation of the CN coherence function is piecewise defined as

$$\mathbb{E}_\varepsilon \left[\log h_{\varepsilon_{uv}}(i'|i, j', j)\right] = \begin{cases} \psi(\tilde{b}_{uv}) - \psi(\tilde{a}_{uv} + \tilde{b}_{uv}) & \text{if } i - i' = j - j' \\ \psi(\tilde{a}_{uv}) - \psi(\tilde{a}_{uv} + \tilde{b}_{uv}) - \log A & \text{otherwise}, \end{cases} \tag{S39}$$

following the expression in (4) and the fact that $q(\varepsilon_{uv}) = \text{Beta}(\tilde{a}_{uv}, \tilde{b}_{uv})$.

Note that the weight $w(uv)$ is the same for all trees that contains the edge $uv$. We use the weight matrix over the fully connected graph, $W(\mathcal{G})$, containing all $w(uv) : uv \in \mathcal{G}$, for sampling trees using 2 and evaluating our sampled trees on the variational distribution $q(T)$.

#### S3.3 Arc distances

The update for $q(\varepsilon_{uv})$ is derived as follows:

$$\begin{aligned} &\log q^*(\varepsilon_{uv}) && \text{(S40)} \\ &= \mathbb{E}_{-\varepsilon_{uv}} \left[\log p(\mathbf{Y}, \mathbf{C}, \mathbf{Z}, T, \boldsymbol{\Psi})\right] \\ &\stackrel{\pm}{=} \mathbb{E}_{CT} \left[\log p(\mathbf{C}|T, \varepsilon)\right] + \log p(\varepsilon_{uv}) \\ &\stackrel{\pm}{=} \mathbb{E}_T \left[\mathbb{E}_{C^u C^v} \left[\log H(C^v|C^u)\right]\right] + \log p(\varepsilon_{uv}) \\ &= \mathbb{E}_T \left[ \sum_{m=2}^{M} \sum_{i,i',j,j'} q(\mathbf{C})vmii' q(\mathbf{C})umjj' \log h_{\varepsilon_{uv}}(i'|i, j', j) \right] \\ &\quad + (a-1)\log(\varepsilon_{uv}) + (b-1)\log(1-\varepsilon_{uv}), && \text{(S41)} \end{aligned}$$

where $q(\mathbf{C})vmii' \coloneqq q(C^v_{m-1} = i, C^v_m = i')$. Furthermore, the summation over the four copy numbers can be split into two separate cases, i.e. the two cases of the CN coherence function, which we denote by $\text{A} \coloneqq \{i, i', j, j' | i - i' = j - j'\}$ and its complementary set $\neg\text{A}$. This being said, we rewrite (S41) as

$$\left( \mathbb{E}_T \left[ \sum_{m=2}^{M} \sum_{ii'jj'}^{\neg \text{A}} q(\mathbf{C})vmii' q(\mathbf{C})umjj' \right] + a - 1 \right) \log(\varepsilon_{uv}) \\ + \left( \mathbb{E}_T \left[ \sum_{m=2}^{M} \sum_{ii'jj'}^{\text{A}} q(\mathbf{C})vmii' q(\mathbf{C})umjj' \right] + b - 1 \right) \log(1 - \varepsilon_{uv})$$

which leaves us with the updated parameters of the variational Beta distribution over $\varepsilon_{uv}$:

$$\begin{aligned} \tilde{a}_{uv} &= \mathbb{E}_T \left[ \sum_{m=2}^{M} \sum_{ii'jj'}^{\neg \text{A}} q(\mathbf{C})vmii' q(\mathbf{C})umjj' \right] + a \\ \tilde{b}_{uv} &= \mathbb{E}_T \left[ \sum_{m=2}^{M} \sum_{ii'jj'}^{\text{A}} q(\mathbf{C})vmii' q(\mathbf{C})umjj' \right] + b \end{aligned} \tag{S42}$$

#### S3.4 Assignment concentration

Derivation of $q(\pi)$:

$$\begin{aligned}
&\log q^*(\boldsymbol{\pi}) \\
&= \mathbb{E}_{-\pi}\left[\log p(\mathbf{Y}, \mathbf{C}, \mathbf{Z}, \mathbf{T}, \boldsymbol{\Psi})\right] \\
&\stackrel{\pm}{=} \mathbb{E}_{-\pi}\left[\log p(\mathbf{Z}|\boldsymbol{\pi})\right] \; + \; \mathbb{E}_{-\pi}\left[\log p(\boldsymbol{\pi})\right] \\
&= \sum_{n=1}^{N}\sum_{k=1}^{K} q(Z_n = k) \log \pi_k + \log p(\boldsymbol{\pi}) \\
&= \sum_{k=1}^{K} \log \pi_k \Big[\sum_{n=1}^{N} q(Z_n = k)\Big] + \sum_{k=1}^{K} \log \pi_k (\delta_k - 1) \\
&= \sum_{k=1}^{K} \log \pi_k \Big[\delta_k + \sum_{n=1}^{N} q(Z_n = k) - 1\Big] .
\end{aligned}$$

Therefore, $q^*(\boldsymbol{\pi})$ is a Dirichlet distribution with parameters $\tilde{\boldsymbol{\delta}} = (\tilde{\delta}_1, \ldots, \tilde{\delta}_K)$, where

$$\tilde{\delta}_k = \delta_k + \sum_{n=1}^{N} q(Z_n = k) \,. \tag{S43}$$

#### S3.5 Cell assignments

Derivation of $q(Z_n)$:

$$\begin{aligned}
&\log q^*(Z_n) \\
&= \mathbb{E}_{-Z_n}\left[\log p(\mathbf{Y}, \mathbf{C}, \mathbf{Z}, T, \boldsymbol{\Psi})\right] \\
&\stackrel{\pm}{=} \mathbb{E}_{-Z_n}\left[\log p(\mathbf{Y}|\mathbf{C}, \mathbf{Z}, T, \boldsymbol{\mu}, \boldsymbol{\tau})\right] + \mathbb{E}_{-Z_n}\left[\log p(\mathbf{Z}|\boldsymbol{\pi})\right]
\end{aligned}$$

Note that $\log p(\mathbf{Y}|\mathbf{Z}, \mathbf{C}, \boldsymbol{\Psi})$ and $\log p(\mathbf{Z}|\boldsymbol{\pi})$ can be written as the sum of two terms, one that depends on $Z_n$ and one that does not, i.e.,

$$\begin{aligned}
&\log p(\mathbf{Y}|\mathbf{Z}, \mathbf{C}, \boldsymbol{\Psi}) \\
&= \sum_{m=1}^{M}\sum_{k=1}^{K}\sum_{j\in\Sigma} \mathbb{1}(Z_n = k)\, \mathbb{1}(C_m^k = j) D_{nmj} \\
&\quad + \sum_{l\neq n}\sum_{m=1}^{M}\sum_{k=1}^{K}\sum_{j\in\Sigma} \mathbb{1}(Z_l = k)\, \mathbb{1}(C_m^k = j) D_{lmj} \,,
\end{aligned}$$

and

$$\log p(\mathbf{Z}|\boldsymbol{\pi}) = \sum_{k=1}^{K} \mathbb{1}(Z_n = k) \log \pi_k + \sum_{l\neq n}\sum_{k=1}^{K} \mathbb{1}(Z_l = k) \log \pi_k \,,$$

where

$$\begin{aligned} D_{nmj} &= \log p(y_{mn} | C_m = j) \\ &= \log \left( \sqrt{\frac{\tau_n}{2\pi}} \exp\left( -\frac{\tau_n}{2} (y_{mn} - j\mu_n)^2 \right) \right) . \end{aligned} \tag{S44}$$

Consequently,

$$\begin{aligned} &\log q^*(Z_n) \\ &\stackrel{\pm}{=} \sum_{k=1}^{K} \mathbb{1}(Z_n = k) \left[ \mathbb{E}_\pi \left[\log \pi_k\right] + \sum_{m=1}^{M} \sum_{j=1}^{J} \mathbb{E}_{C^k} \left[ \mathbb{1}(C_m^k = j) \right] \mathbb{E}_{\mu_n \tau_n} \left[ D_{nmj} \right] \right] . \end{aligned}$$

We simplify the expression further by:

$$\hat{\gamma}_{nk} = \mathbb{E}_\pi \left[\log \pi_k\right] + \sum_{m=1}^{M} \sum_{j \in \Sigma} q(C_m^k = j) \mathbb{E}_{\mu_n \tau_n} \left[ D_{nmj} \right] , \tag{S45}$$

with $\mathbb{E}_\pi \left[\log \pi_k\right] = \psi(\tilde{\delta}_k) - \psi(\sum_r \tilde{\delta}_r)$, $\mathbb{E}_{\mu_n \tau_n} \left[ D_{nmj} \right]$ computed as in (S29). Finally, we set:

$$\tilde{\gamma}_n = \frac{\exp(\hat{\gamma}_{nk})}{\sum_{l=1}^{K} \exp(\hat{\gamma}_{nl})} \tag{S46}$$

and conclude that $q^*(Z_n) \sim \text{Categorical}(\tilde{\gamma}_n)$ with parameters $\tilde{\gamma}_n = (\tilde{\gamma}_{n1}, \ldots, \tilde{\gamma}_{nK})$

#### S3.6 Baseline mean and precision

We use the Normal-Gamma joint prior on $\mu_n$ and $\tau_n$, that factorizes over $n$, and introduce a joint variational distribution on the corresponding parameters. The mean-field standard update becomes:

$$\begin{aligned} &\log q^*(\mu_n, \tau_n) \\ &= \mathbb{E}_{-(\mu_n, \tau_n)} \left[ \log p(\mathbf{Y}, \mathbf{C}, \mathbf{Z}, \boldsymbol{\Psi}) \right] \\ &\stackrel{\pm}{=} \mathbb{E}_{-(\mu_n, \tau_n)} \left[ \log p(\mathbf{Y} | \mathbf{C}, \mathbf{Z}, \boldsymbol{\Psi}) \right] + \log p(\mu_n, \tau_n) \end{aligned}$$

Simplifying each term individually yields:

$$\begin{aligned} &\mathbb{E}_{-(\mu_n, \tau_n)} \left[ \log p(\mathbf{Y} | \mathbf{C}, \mathbf{Z}, \boldsymbol{\Psi}) \right] \\ &\stackrel{\pm}{=} \mathbb{E}_{-(\mu_n, \tau_n)} \left[ \log \prod_{m,u,c} p(y_{mn} | C_m^u, \mu_n, \tau_n)^{\mathbb{1}(Z_n = u, C_m^u = c)} \right] \\ &\stackrel{\pm}{=} \sum_{m,u,c} q(C_m^u = c) q(Z_n = u) \left( \frac{1}{2} \log \tau_n - \frac{1}{2} \tau_n (y_{mn} - c \cdot \mu_n)^2 \right) \end{aligned} \tag{S47}$$

and

$$\log p(\mu_n, \tau_n) \stackrel{\pm}{=} (\alpha - \frac{1}{2}) \log \tau_n - \beta \tau_n - \frac{1}{2} \tau_n \lambda (\mu_n - \mu)^2 .$$

As our model prior is Gamma-Normal distributed, we suspect our variational distribution to be in the same family, and attempt to match the parameters associated with the independent variables of the Gamma-Normal. First we regroup the variables as follows:

$$\begin{aligned}
&\log q^*(\mu_n, \tau_n) \\
&\stackrel{+}{=} -(\frac{\tau_n}{2})\big[\mu_n^2 \cdot \big(\lambda + \sum_{m,u,c} q(C_m^u = c) q(Z_n = u) \cdot c^2\big) \\
&\quad - \mu_n \big(\lambda\mu + \sum_{m,u,c} q(C_m^u = c) q(Z_n = u) \cdot c \cdot y_{mn}\big)\big] \\
&\quad + (\alpha + \frac{M}{2} - \frac{1}{2}) \log \tau_n - \tau_n (\beta + \frac{1}{2}(\mu^2\lambda + \sum_m y_{mn}^2))\,.
\end{aligned}$$

In order to get the expression on Normal-Gamma form we need to complete the square for the cross term. We do this by adding and subtracting the square of the factor of $\mu_n \tau_n$ and then multiplying it with $\tau_n$; the negative term is then passed to complete the square and the positive is added to the $\tau_n$ term:

$$\begin{aligned}
&\log q^*(\mu_n, \tau_n) \\
&\stackrel{+}{=} -(\frac{\tau_n}{2})(\lambda + \Sigma_{C,Z}(c^2))\big[\mu_n - \frac{(\lambda\mu + \Sigma_{C,Z}(c, y_{mn}))}{\lambda + \Sigma_{C,Z}(c^2)}\big]^2 \\
&\quad + (\alpha + \frac{M}{2} - \frac{1}{2}) \log \tau_n - \tau_n (\beta + \frac{1}{2}(\mu^2\lambda + \sum_m y_{mn}^2) - \frac{1}{2} \cdot \frac{(\mu\lambda + \Sigma_{C,Z}(c, y_{n,m}))^2)}{\lambda + \Sigma_{C,Z}(c^2)})
\end{aligned}$$

where

$$\begin{aligned}
\Sigma_{C,Z}(c^2) &= \sum_{m,u,c} q(Z_n = u) q(C_m^u = c) \cdot c^2 \\
\Sigma_{C,Z}(c, y_{mn}) &= \sum_{m,u,c} q(Z_n = u) q(C_m^u = c) \cdot c \cdot y_{mn}\,.
\end{aligned}$$

By matching the independent variables with the parameters we see that $q^*(\mu_n, \tau_n)$ is indeed Normal-Gamma$(\tilde{\nu}_n, \tilde{\lambda}_n, \tilde{\alpha}_n, \tilde{\beta}_n)$ distributed with parameters:

$$\begin{aligned}
\tilde{\nu}_n &= \frac{\mu\lambda + \Sigma_{C,Z}(c, y_{mn})}{\lambda + \Sigma_{C,Z}(c^2)} \\
\tilde{\lambda}_n &= \lambda + \Sigma_{C,Z}(c^2) \\
\tilde{\alpha}_n &= \alpha + \frac{M}{2} \\
\tilde{\beta}_n &= \beta + \frac{1}{2}(\mu^2\lambda + \sum_m y_{mn}^2) - \frac{1}{2}\tilde{\nu}_n^2 \tilde{\lambda}_n\,.
\end{aligned} \tag{S48}$$

### S4 ELBO computation

In this section we derive the entropy and negative cross-entropy terms of 10 in 4.3.

Since the expectation over the tree topology $T$ in 10 requires a sampling technique to be estimated, i.e., it cannot be derived as for the other variables in closed form, we separate the expectation into two main parts:

$$\mathbb{E}_T\Big[\mathbb{E}_{-T}\Big[\log p(\mathbf{Y}|\mathbf{C},\mathbf{Z},\boldsymbol{\Psi}) + \log p(\mathbf{C}|\mathbf{Z},T,\boldsymbol{\Psi}) + \log p(\mathbf{Z}|\boldsymbol{\Psi}) + \log p(\boldsymbol{\Psi})\Big] + \log p(T)\Big] + \mathbb{H}_q(\mathbf{C},\mathbf{Z},T,\boldsymbol{\Psi})\,.$$

In the subsections below we present each component of the ELBO, by deriving the negative cross entropy terms of in the inner expectation and the variational distribution entropy, the latter which we write following the notation:

$$\mathbb{H}_q(x) = -\mathbb{E}_{q(x)}\left[\log q(x)\right]\,.$$

### S4.1 Observations

$$\begin{aligned}
&\mathbb{E}_{CZ\Psi}\left[\log p(\mathbf{Y}|\mathbf{C},\mathbf{Z},\boldsymbol{\Psi})\right] \qquad \text{(S49)}\\
&= \mathbb{E}_{CZ\Psi}\left[\sum_{v,n,m,j} \mathbb{1}(Z_n = v)\,\mathbb{1}(C_m^v = j)\log p(y_{mn}|C_m^v = j)\right]\\
&= \frac{1}{2}\sum_{v,n,m,j} q(Z_n = v)q(C_m^v = j)\mathbb{E}_{\mu_n\tau_n}\left[\log\tau_n - \tau_n\left(y_{mn} - j\mu_n\right)^2 - \log 2\pi\right]\\
&= \frac{1}{2}\sum_{v,n,m,j} q(Z_n = v)q(C_m^v = j)\big[\mathbb{E}_{\tau_n}\left[\log\tau_n\right] - y_{mn}^2\mathbb{E}_{\tau_n}\left[\tau_n\right] + 2jy_{mn}\mathbb{E}_{\mu_n\tau_n}\left[\mu_n\tau_n\right]\\
&\quad - j^2\mathbb{E}_{\mu_n\tau_n}\left[\mu_n^2\tau_n\right] - \log 2\pi\big]\,.
\end{aligned}$$

Note that the values of the expectations above are standard expectations for Normal and Gamma random variables and can be found in subsection S3.1.

### S4.2 Copy numbers

$$\begin{aligned}
&\mathbb{E}_{CZ\Psi}\left[\log p(\mathbf{C}|\mathbf{Z},T,\boldsymbol{\Psi})\right] \qquad \text{(S50)}\\
&= \mathbb{E}_{C^r}\left[\log p(\mathbf{C}^r)\right] + \sum_{uv\in A(T)}\mathbb{E}_{C^uC^v\varepsilon}\left[\log p(\mathbf{C}^v|\mathbf{C}^u,\varepsilon)\right]\\
&= \mathbb{E}_{C^r}\left[\log p(\mathbf{C}^r)\right]\\
&\quad + \sum_{uv\in A(T)}\Big(\sum_{i,j} q(C_1^v = j)q(C_1^u = i)h_{\varepsilon_0}\\
&\qquad + \sum_{m=2}^{M}\sum_{ii'jj'} q(\mathbf{C})umii'q(\mathbf{C})vmjj'\mathbb{E}_{\varepsilon}\left[\log h_{\varepsilon}(j'|j,i',i)\right]\Big)\,,
\end{aligned}$$

where $\mathbb{E}_\varepsilon\left[\log h_\varepsilon(j'|j,i',i)\right]$ is computed as shown in (S39) and the variational distribution quantities (denoted as in Equation (S41)) for the copy numbers are computed with the forward-algorithm.

Furthermore, we can calculate the entropy by use of the marginals $q(C_m)$ in the following way:

$$\mathbb{H}_q(C) = -\sum_{umi} q(C_m^u = i) \log q(C_m^u = i)\,. \tag{S51}$$

#### S4.3 Cell assignments

$$\begin{aligned}
&\mathbb{E}_{Z\Psi}\left[\log p(\mathbf{Z}|\boldsymbol{\Psi})\right] \\
&= \sum_n \mathbb{E}_{Z_n\pi}\left[\log p(Z_n|\boldsymbol{\pi})\right] \\
&= \sum_n \sum_k q(Z_n = k)\mathbb{E}_\pi\left[\log \pi_k\right]\,,
\end{aligned} \tag{S52}$$

with $\mathbb{E}_\pi\left[\log \pi_k\right] = \psi(\tilde{\delta}_k) - \psi(\delta_0)$, $\tilde{\delta}_0 = \sum_i \tilde{\delta}_i$ such that $\tilde{\boldsymbol{\delta}}$ is the parameter of the variational Dirichlet distribution over $\boldsymbol{\pi}$.

The entropy is calculated as:

$$\mathbb{E}_{Z\Psi}\left[\log q(\mathbf{Z})\right] = \sum_n \sum_k q(Z_n = k) \log \tilde{\gamma}_{nk}\,.$$

#### S4.4 Concentration parameter

$$\begin{aligned}
&\mathbb{E}_\pi\left[\log p(\boldsymbol{\pi})\right] \\
&= \int_{\boldsymbol{\pi}} q(\boldsymbol{\pi}') \log p(\boldsymbol{\pi}') d\boldsymbol{\pi}' \\
&= \int_{\boldsymbol{\pi}} q(\boldsymbol{\pi}') \left( \sum_k (\delta_k - 1) \log \pi'_k - \log B(\boldsymbol{\delta}) \right) d\boldsymbol{\pi}' \\
&= \sum_k (\delta_k - 1) \int_{\boldsymbol{\pi}} q(\boldsymbol{\pi}') \log \pi'_k d\boldsymbol{\pi}' - \log B(\boldsymbol{\delta}) \int_{\boldsymbol{\pi}} q(\boldsymbol{\pi}') d\boldsymbol{\pi}' \\
&= \sum_k (\delta_k - 1) \mathbb{E}_{\pi\sim q}\left[\log \pi_k\right] - \log B(\boldsymbol{\delta}) \\
&= \sum_k (\delta_k - 1) \psi(\tilde{\delta}_k) - (\delta_0 - K)\psi(\tilde{\delta}_0) - \log B(\boldsymbol{\delta})
\end{aligned} \tag{S53}$$

where

$$B(\boldsymbol{\delta}) = \frac{\prod_k \mathit{\Gamma}(\delta_k)}{\mathit{\Gamma}(\sum_k \delta_k)}\,, \qquad \tilde{\delta}_0 = \sum_{k=1}^{K} \tilde{\boldsymbol{\delta}}_k$$

and $\tilde{\boldsymbol{\delta}}$ refers to the variational concentration parameter, while $\boldsymbol{\delta}$ is the generative model parameter. The entropy of $q(\boldsymbol{\pi})$ is found by replacing the model parameter $\boldsymbol{\delta}$ with $\tilde{\boldsymbol{\delta}}$.

#### S4.5 Gaussian mean and precision

$$
\begin{aligned}
&\mathbb{E}_{\mu\tau}\left[\log p(\boldsymbol{\mu}, \boldsymbol{\tau})\right] \\
&= \sum_n \int_{\mu_n} \int_{\tau_n} q(\mu_n' \tau_n') \log p(\mu_n', \tau_n') d\tau_n' d\mu_n' \\
&= \sum_n \int_{\mu_n} \int_{\tau_n} q(\mu_n' | \tau_n') q(\tau_n') \big( \log p(\mu_n' | \tau_n') + \log p(\tau_n) \big) d\tau_n' d\mu_n' \\
&= \sum_n \Bigg[ \int_{\tau_n} q(\tau_n') \left( \int_{\mu_n} q(\mu_n' | \tau_n') \log p(\mu_n' | \tau_n') d\mu_n' \right) d\tau_n' \\
&\quad + \int_{\tau_n} q(\tau_n') \log p(\tau_n) \left( \int_{\mu_n} q(\mu_n' | \tau_n') d\mu_n' \right) d\tau_n' \Bigg] .
\end{aligned} \tag{S54}
$$

The above expression is made of two terms (excluding the outermost summation): the first double integral which is the expectation over $q(\tau_n)$ of the negative cross-entropy $-H_{\mu_n|\tau_n}(q,p)$, and the second integral which is just the negative cross-entropy over $\tau_n$ of $q$ and $p$, i.e. $-H_{\tau_n}(q,p)$.

Therefore we get

$$
\begin{aligned}
&\mathbb{E}_{\mu\tau}\left[\log p(\boldsymbol{\mu}, \boldsymbol{\tau})\right] \\
&= \sum_n \mathbb{E}_{q(\tau_n)}\left[ \mathbb{E}_{q(\mu_n|\tau_n)}\left[\log p(\mu_n|\tau_n)\right]\right] + \mathbb{E}_{q(\tau_n)}\left[\log p(\tau_n)\right] .
\end{aligned}
$$

We compute the partial terms below (omitting $q$ as all the expectations here are taken with respect to the variational distribution)

$$
\begin{aligned}
\mathbb{E}_{\mu_n|\tau_n}\left[\log p(\mu_n|\tau_n)\right] &= \frac{1}{2}\log\frac{\lambda\tau_n}{2\pi} - \frac{\lambda\tau_n}{2}\mathbb{E}_{\mu_n|\tau_n}\left[(\mu_n - \mu)^2\right] \\
&= \frac{1}{2}\log\frac{\lambda\tau_n}{2\pi} - \frac{\lambda\tau_n}{2}\mathrm{Var}_{\mu_n|\tau_n}\left[\mu_n\right] \\
&= \frac{1}{2}\left(\log\lambda - \log 2\pi - \frac{\lambda}{\tilde{\lambda}_n}\right) + \frac{1}{2}\log\tau_n
\end{aligned} \tag{S55}
$$

$$
\begin{aligned}
\mathbb{E}_{\tau_n}\left[\mathbb{E}_{\mu_n|\tau_n}\left[\log p(\mu_n|\tau_n)\right]\right] &= \frac{1}{2}\left(\log\lambda - \log 2\pi - \frac{\lambda}{\tilde{\lambda}_n}\right) + \frac{1}{2}\mathbb{E}_{\tau_n}\left[\log\tau_n\right] \\
&= \frac{1}{2}\left(\log\lambda - \log 2\pi - \frac{\lambda}{\tilde{\lambda}_n} + \psi(\tilde{\alpha}_n) - \log\tilde{\beta}_n\right)
\end{aligned} \tag{S56}
$$

$$
\begin{aligned}
\mathbb{E}_{\tau_n}\left[\log p(\tau_n)\right] &= \log\frac{\beta^\alpha}{\Gamma(\alpha)} + (\alpha - 1)\mathbb{E}_{\tau_n}\left[\log\tau_n\right] - \beta\mathbb{E}_{\tau_n}\left[\tau_n\right] \\
&= \log\frac{\beta^\alpha}{\Gamma(\alpha)} + (\alpha - 1)(\psi(\tilde{\alpha}_n) - \log\tilde{\beta}) - \beta\frac{\tilde{\alpha}_n}{\tilde{\beta}_n}
\end{aligned} \tag{S57}
$$

and then put altogether:

$$
\begin{aligned}
&\mathbb{E}_{\mu\tau}\left[\log p(\boldsymbol{\mu}, \boldsymbol{\tau})\right] \\
&= \sum_n \Big[ \frac{1}{2} \left( \log \lambda - \log 2\pi - \frac{\lambda}{\tilde{\lambda}_n} + \psi(\tilde{\alpha}_n) - \log \tilde{\beta}_n \right) \\
&\quad + \log \frac{\beta^\alpha}{\Gamma(\alpha)} + (\alpha - 1)(\psi(\tilde{\alpha}_n) - \log \tilde{\beta}) - \beta \frac{\tilde{\alpha}_n}{\tilde{\beta}_n} \Big] \\
&= -\frac{N}{2} \log 2\pi + \sum_n \frac{1}{2} \left( \log \lambda - \frac{\lambda}{\tilde{\lambda}_n} \right) + (\alpha - \frac{1}{2})(\psi(\tilde{\alpha}_n) - \log \tilde{\beta}_n) \\
&\quad + \alpha \log \beta - \log \Gamma(\alpha) - \beta \frac{\tilde{\alpha}_n}{\tilde{\beta}_n} \,.
\end{aligned} \tag{S58}
$$

The variational entropy is computed as follows

$$
\begin{aligned}
\mathbb{H}_q(\boldsymbol{\mu}, \boldsymbol{\tau}) &= \sum_n \mathbb{E}_{\tau_n} \left[ \mathbb{H}_q(\mu_n | \tau_n) \right] + \mathbb{H}_q(\tau_n) \\
&= \sum_n \frac{1}{2} \left( \log 2\pi + 1 - \log \tilde{\lambda}_n - \psi(\tilde{\alpha}_n) + \log \tilde{\beta}_n \right) \\
&\quad + \tilde{\alpha}_n - \log \tilde{\beta}_n + \log \Gamma(\tilde{\alpha}_n) + (1 - \tilde{\alpha}_n)\psi(\tilde{\alpha}_n) \\
&= \frac{N}{2} \log 2\pi + \sum_n \frac{1}{2} \left( 1 - \log \tilde{\beta}_n - \log \tilde{\lambda}_n \right) \\
&\quad + \left( \frac{1}{2} - \tilde{\alpha}_n \right) \psi(\tilde{\alpha}_n) + \tilde{\alpha}_n + \log \Gamma(\tilde{\alpha}_n) \,.
\end{aligned} \tag{S59}
$$

#### S4.6 Epsilon

$$
\begin{aligned}
&\mathbb{E}_{\varepsilon, T}\left[\log p(\boldsymbol{\varepsilon})\right] \\
&= \mathbb{E}_{T \sim q(T)} \left[ \sum_{uv \in T} \mathbb{E}_{\varepsilon_{uv}} \left[ \log p(\varepsilon_{uv}) \right] \right] \\
&= \mathbb{E}_{T \sim q(T)} \left[ \sum_{uv \in T} \mathbb{E}_{\varepsilon_{uv}} \left[ \log B(a, b) - (a - 1) \log \varepsilon_{uv} + (b - 1) \log(1 - \varepsilon_{uv}) \right] \right] \\
&= \mathbb{E}_{T \sim q(T)} \left[ \sum_{uv \in T} \log B(a, b) - (a - 1)[\psi(\tilde{a}_{uv}) - \psi(\tilde{a}_{uv} + \tilde{b}_{uv})] + (b - 1)[\psi(\tilde{b}_{uv}) - \psi(\tilde{b}_{uv} + \tilde{a}_{uv})] \right]
\end{aligned} \tag{S60}
$$

$$
\begin{aligned}
&\mathbb{E}_{\varepsilon, T}\left[\log q(\boldsymbol{\varepsilon})\right] \\
&= \mathbb{E}_{T \sim q(T)} \left[ \sum_{uv \in T} H(q(\varepsilon_{uv})) \right] \\
&= \mathbb{E}_{T \sim q(T)} \left[ \sum_{uv \in T} \log B(\tilde{a}_{uv}, \tilde{b}_{uv}) - (\tilde{a}_{uv} - 1)\psi(\tilde{a}_{uv}) - (\tilde{b}_{uv} - 1)\psi(\tilde{b}_{uv}) + (\tilde{a}_{uv} + \tilde{b}_{uv} + 2)\psi(\tilde{a}_{uv} + \tilde{b}_{uv}) \right]
\end{aligned} \tag{S61}
$$

#### S4.7 Tree topology

$$\mathbb{E}_T\left[\log q(T)\right] = -\sum_{T \in \mathcal{T}} q(T) \log q(T) \tag{S62}$$

since $q(T)$ is unnormalized, we use importance sampling to estimate the normalizing constant and evaluate on $q(T) = \frac{\sum_{l=1}^{L} w_l \tilde{q}(T)}{\sum_{l=1}^{L} w_l}$

$$\begin{aligned} \mathbb{E}_T\left[\log p(T)\right] &= \sum_{T \in \mathcal{T}} q(T) \left[-\log |\mathcal{T}|\right] \\ &= -\log |\mathcal{T}| \sum_{T \in \mathcal{T}} q(T) \\ &= -\log |\mathcal{T}|, \end{aligned} \tag{S63}$$

where in case of $K$ nodes and considering labeled arborescences, $|\mathcal{T}| = K^{K-2}$ [5].

### S5 VICTree

#### S5.1 VICTree pseudocode

The algorithm complexity is dominated by the update of $q(\mathbf{C})$ parameters, which requires $O(KMNA^2)$, at every iteration and for every unique tree in the sample.

**Algorithm 1** VICTree pseudocode

**Input:** number of importance samples $L$, number of iterations $I$, initialized $q$.
**for** $i = 1, \ldots, I$ **do**
  update $W(\mathcal{G})$ using (9)
  sample $T_l = T_1, \ldots, T_L$ trees using alg. 2
  **for** $T_l = T_1, \ldots, T_L$ **do**
    Update $q_l(\mathbf{C}), q(\boldsymbol{\varepsilon})$ using $T_l$ and (8,7)
  **end for**
  $q(\mathbf{C}) \longleftarrow \sum_l w_{T_l} \cdot q_l(\mathbf{C})$
  Update $q(\mathbf{Z}), q(\boldsymbol{\pi}), q(\boldsymbol{\mu}, \boldsymbol{\tau})$ using parameters in (7)
**end for**
**return**

#### S5.2 VICTree Importance sampling details

During inference process, we need to evaluate expectations over $q(T)$ of some function of the tree, but enumerating over all possible arborescences can be costly when considering directed labeled trees with several nodes. More specifically, by Cayley's formula [5] and Prüfer sequences it is proven that the number of labeled trees with $n$ nodes is $n^{n-2}$. In our case the trees are rooted and directed, thus the topology space is of size $n^{n-1}$, which quickly becomes intractable.

Assuming we can draw samples from $q(T)$ distribution, we can approximate an expectation as follows.

$$\mathbb{E}_{q(T)}\left[f(T)\right] \approx \frac{1}{L}\sum_{l=1}^{L} f(T_l)\,. \tag{S64}$$

However, sampling from $q(T)$ is not straightforward, therefore we sample from an alternative distribution $g(T)$ and apply importance sampling [14] in order to evaluate the desired expectation. Let $\tilde{q}(T)$ denote the weight of a arborescence $T$, which is defined to be the product over the edge weights and let $w(T) = \tilde{q}(T)/\tilde{g}(T)$ be the importance weights. Then

$$\mathbb{E}_{q(T)}\left[f(T)\right] = \mathbb{E}_{g(T)}\left[\frac{\tilde{q}(T)}{\tilde{g}(T)} f(T)\right] \approx \frac{\sum_{l=1}^{L} w(T_l) f(T_l)}{\sum_{l=1}^{L} w(T_l)}\,. \tag{S65}$$

Self-normalization (S65) is required since both $q(T)$ and $g(T)$ are evaluated up to a multiplication factor.

### S6 LARS algorithm

The following algorithm runs with computational time $O(|E|K^3)$ since Edmond's algorithm requires $O(K^2)$. It is then clear that, for bigger trees, the complexity of a tree sample quickly overcomes the complexity of the other costly operations in each VICTree step, especially the copy-number variational distribution update $q(\mathbf{C})$ as mentioned in Section S5.1.

**Algorithm 2** Labeled Arborescence Sampler (LARS)

**Input:** $\mathcal{G}$ fully connected graph, $W$ corresponding adjacency matrix.
$E \leftarrow$ ordering of arcs in $\mathcal{G}, S \leftarrow \emptyset, g \leftarrow 1$
**while** $i < K - 1$ **do**
  **for** $e \in E$ **do**
    $T \leftarrow \text{Edmonds}(A(S, e))$
    $T' \leftarrow \text{Edmonds}(A'(S, e))$
    $\theta \leftarrow \frac{\tilde{q}(T)}{\tilde{q}(T)+\tilde{q}(T')}$
    $u \sim U(0, 1)$
    **if** $\theta \geq u$ **then**
      $S \leftarrow S \cup e$
      $g \leftarrow g \times \theta$
    **end if**
  **end for**
**end while**
**return** $S, g$

### S7 Split-merge

We run our split-merge algorithm every 10th iteration during optimization. First, we merge by checking the L1 distance between Viterbi paths of all clusters CNPs $q(C^i)$, $q(C^j)$. If the distance is smaller than a threshold, we merge $j$ into $i$ by re-initializing the parameters of $q(C^j)$ randomly, setting $q(Z_n = j) = 0$ and doubling $q(Z_n = i)$. Then, if there exists a degenerate cluster, $l$, i.e. a cluster with total probability $\sum q(Z_n = l)$ smaller than a threshold, we run the split algorithm: For all candidate clusters $k$ with $\sum q(Z_n = k) > 0.01$ we group the cells with $\arg\max_j q(Z_n = j) = k$ into two groups, $k_1$ and $k_2$ by KMeans. Then, we construct temporary $q(C^{k_1})$ and $q(C^{k_2})$ by applying the CAVI update using only the cells in $k_1$ and $k_2$ respectively, and then evaluate the ELBO using these new $q(C)$s. We select to split the candidate cluster $k$ that results in the maximum ELBO into $l$.
